# Exploiting fluctuations in gene expression to detect causal interactions between genes

**DOI:** 10.1101/2023.09.01.555799

**Authors:** Euan Joly-Smith, Mir Mikdad Talpur, Paige Allard, Fotini Papazotos, Laurent Potvin-Trottier, Andreas Hilfinger

## Abstract

Characterizing and manipulating cellular behaviour requires a mechanistic understanding of the causal interactions between cellular components. We present an approach to detect causal interactions between genes without the need to perturb the physiological state of cells. This approach exploits naturally occurring cell-to-cell variability which is experimentally accessible from static population snapshots of genetically identical cells without the need to follow cells over time. Our main contribution is a simple mathematical relation that constrains the propagation of gene expression noise through biochemical reaction networks. This relation allows us to rigorously interpret fluctuation data even when only a small part of a complex gene regulatory process can be observed. We show how this relation can, in theory, be exploited to detect causal interactions by synthetically engineering a passive reporter of gene expression, akin to the established “dual reporter assay”. While the focus of our contribution is theoretical, we also present an experimental proof-of-principle to demonstrate the real-world applicability of our approach in certain circumstances. Our experimental data suggest that the method can detect causal interactions in specific synthetic gene regulatory circuits in *E. coli* confirming our theoretical result in a narrow set of controlled experimental settings. Further work is needed to show that the approach is practical on a large scale, with naturally occurring gene regulatory networks, or in organisms other than *E. coli*.

---

Translating molecular abundance data into mechanistic models of cellular processes remains a major challenge of systems biology. High-throughput experimental techniques routinely produce statistical associations between genes through correlation measurements of gene expression variability [1–5]. While such statistical associations can provide insightful clues, they do not directly identify causal interactions: even if two components *X* and *Z* are non-spuriously correlated, we cannot conclude that *X* affects *Z* because the causal connection could be reversed or a confounding factor could be regulating both components, even with statistical associations that go beyond correlations such as Granger causality [6, 7].

Perturbation experiments are a conceptually simple solution to avoid this problem and directly infer causal relationships in gene regulatory networks [8–10]. However, they come with practical challenges: drug perturbations can affect multiple targets at once [11] and genetic perturbations, e.g., through changing gene copy numbers, are not guaranteed to keep cells in their physiologically relevant regime [12, 13]. For large enough perturbations, everything is expected to affect everything else in the cell.

Here we present a novel mathematical relation which can, in theory, be exploited to infer directional causal interactions between cellular components by utilizing stochastic fluctuations of cellular abundances in unperturbed cells. Existing work on analyzing non-genetic variability has focused on testing completely specified mechanistic models against such fluctuation data [1, 14– 17]. However, completely specifying mechanistic models of cellular processes often requires making a large number of assumptions about unknown details, which can make inferences based on such an approach unreliable [18, 19].

We thus propose a novel inference method to detect whether a gene *X* causally affects a gene *Z* by exploiting a mathematical identity that constrains an entire class of models. Under certain assumptions, we show how covariability measurements of molecular abundances can, in theory, be used to detect causal interactions by specifying only “local” aspects of the underlying gene expression dynamics [19–21].

Experimentally, our proposed inference method is based on an approach similar to the “dual reporter assay” [22–25] previously established to quantify stochastic fluctuations in expression differences between copies of a gene of interest. However, instead of analyzing the covariance between the dual reporters, we focus on the covariance of each reporter with a third component of biological interest. We prove that under broad conditions these covariances must be identical in the absence of the causal interaction we wish to establish. Violations of this relation can thus be used to detect causal interactions.

To illustrate the generality of our analytically proven covariance identity, we numerically verify the result in a wide variety of example systems. Additionally, we present an experimental proof-of-principle for a specific set of synthetically constructed gene regulatory circuits in *E. coli* with known interactions. The data establish a baseline estimate for the accuracy of our approach with current experimental methods. Despite the arguable occurrence of a false positive result, the experimental data support our hypothesis that single-cell variability measurements might prove useful in detecting causal interactions in gene regulatory networks.

However, we have not shown that the approach is practical on a large scale, we have not tested our approach using naturally occurring gene regulatory networks, and we have not tested our approach in organisms other than *E. coli*. Additional experimental tests beyond our proof-of-principle demonstration are thus needed to establish the scalability of our method and to show that the method reliably detects causal interactions between endogenous genes in more general experimental settings.

## Results

### Invariant covariance relation in the absence of causal interactions

Our main mathematical result is a covariance invariant that constrains the fluctuations of two molecular species in a partially specified reaction network. We consider a molecular species of interest *X* that is subject to firstorder degradation and is made with an unspecified and time-varying production rate. A reporter species *Y* acts as an identical but separate copy of *X* that is subject to the same control input. This specifies the following stochastic reactions within a larger reaction network

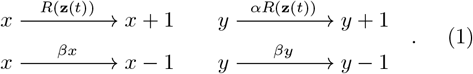

Here *x* and *y* denote the number of copies of molecular species *X* and *Y* respectively, *α* is an arbitrary proportionality constant, and the shared production rate *R* can depend in arbitrary ways on the abundances *z*_*k*_ of any cellular component (including *X* and *Y* themselves), collectively denoted as **Z** := {*Z*_1_, *Z*_2_, …}. The birth-death events denoted in Eq. (1) produce or remove one molecule of *X* or *Y* at a time and occur stochastically with the state-dependent rates indicated above the arrows.

If the level of *X* affects the rate of production or degradation of another component, say *Z*_3_, we say *X* causally affects *Z*_3_ and we define causal effects transitively, i.e., if *X* affects *Z*_3_, and *Z*_3_ affects *Z*_5_, then *X* affects *Z*_5_. In other words, *X* causally affects a cellular component if a directed path can be drawn from *X* to said component in the biochemical reaction network as illustrated in Fig. 1A

**Figure 1.**
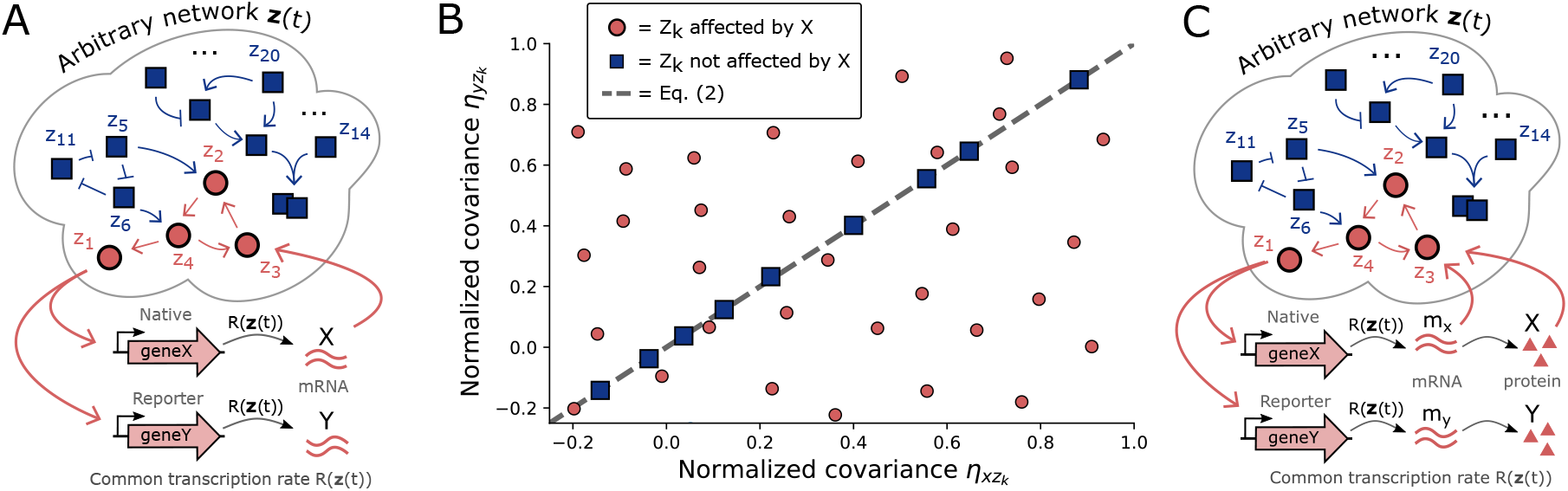
Causal interactions between genes can be inferred when non-genetic cell-to-cell variability violates the covariance invariant of Eq. (2). **A)** Consider an arbitrary network of interacting cellular components in which an engineered reporter *Y* is introduced to act as a passive read-out of the transcriptional signal that regulates a gene of interest *geneX*, but itself does not regulate other cellular components. Any cellular component belongs to one of two groups: components affected by *X* (red circles), and components not affected by *X* (blue squares), where the arrows indicate directed causal biochemical effects. Specifically, an arrow from *Z*_*i*_ to *Z*_*j*_ denotes the existence of a reaction that changes *Z*_*j*_ with a rate that depends on *Z*_*i*_ (see SI for details). **B)** The dashed line of Eq. (2) constrains the normalized covariance between *X* and any cellular component *Z*_*k*_ not affected by *X* (blue squares). In contrast, components *Z*_*k*_ affected by *X* (red circles), are not constrained by Eq. (2). A violation of Eq. (2) thus implies the existence of a causal interaction from *X* to *Z*_*k*_. Data points are numerical simulations of specific example networks (see SI for details) to illustrate the analytically proven theorem of Eq. (2). **C)** The invariant of Eq. (2) applies not only to transcriptional reporters but also when *X* and *Y* correspond to co-regulated fluorescent proteins with translation rates that scale with transcript abundances.

*Theorem*: if neither *X* nor *Y* causally affect a third cellular component of interest *Z*_*k*_ ∈ **Z**, then

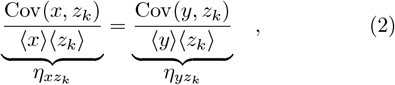

where angular brackets denote ensemble averages, and 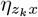 denotes the normalized covariance between *Z*_*k*_ and *X* in a population of genetically identical cells that have reached a time-independent distribution of cell-to-cell variability and do not exhibit permanently zero degradation rates for *X* and *Y*.

Eq. (2) makes no assumptions on the interactions of all the unspecified cellular components and allows arbitrary effects from the component of interest *Z*_*k*_ onto *X* and *Y*. The invariant constrains any cellular component *Z*_*k*_ that is not affected by *X* and *Y* as long as the “local” dynamics of *X* and *Y* is given by Eq. (1). The detailed proof of Eq. (2) is presented in SI. The key condition under which it holds is that the non-genetic population variability has reached a stationary state. In systems in which there is uncertainty about this condition it can be directly verified experimentally.

Note, a violation of Eq. (2) implies the presence of a causal connection but not vice versa. This is a logical necessity because the dynamics of causally connected genes can be arbitrarily close to that of non-interacting genes.

In a later section, we discuss further generalizations of the dual reporter class defined in Eq. (1), allowing for growing and dividing cells, measurement noise, and fluctuations in degradation rates as would be relevant for its application to experimental single-cell data. Note that due to the possibility of feedback, the dynamics of *X* and *Y* is not generally symmetric even though *X* and *Y* are co-regulated [20], see Fig. S1A. Eq. (2) is a statement about stochastic covariability and not a trivial relation based on the (incorrect) assumption that *x*(*t*) = *y*(*t*). For example, the Pearson correlation coefficients 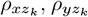 are not necessarily equal even when *X* and *Y* do not affect *Z*_*k*_, see Fig. 2 and Fig. S1B.

**Figure 2.**
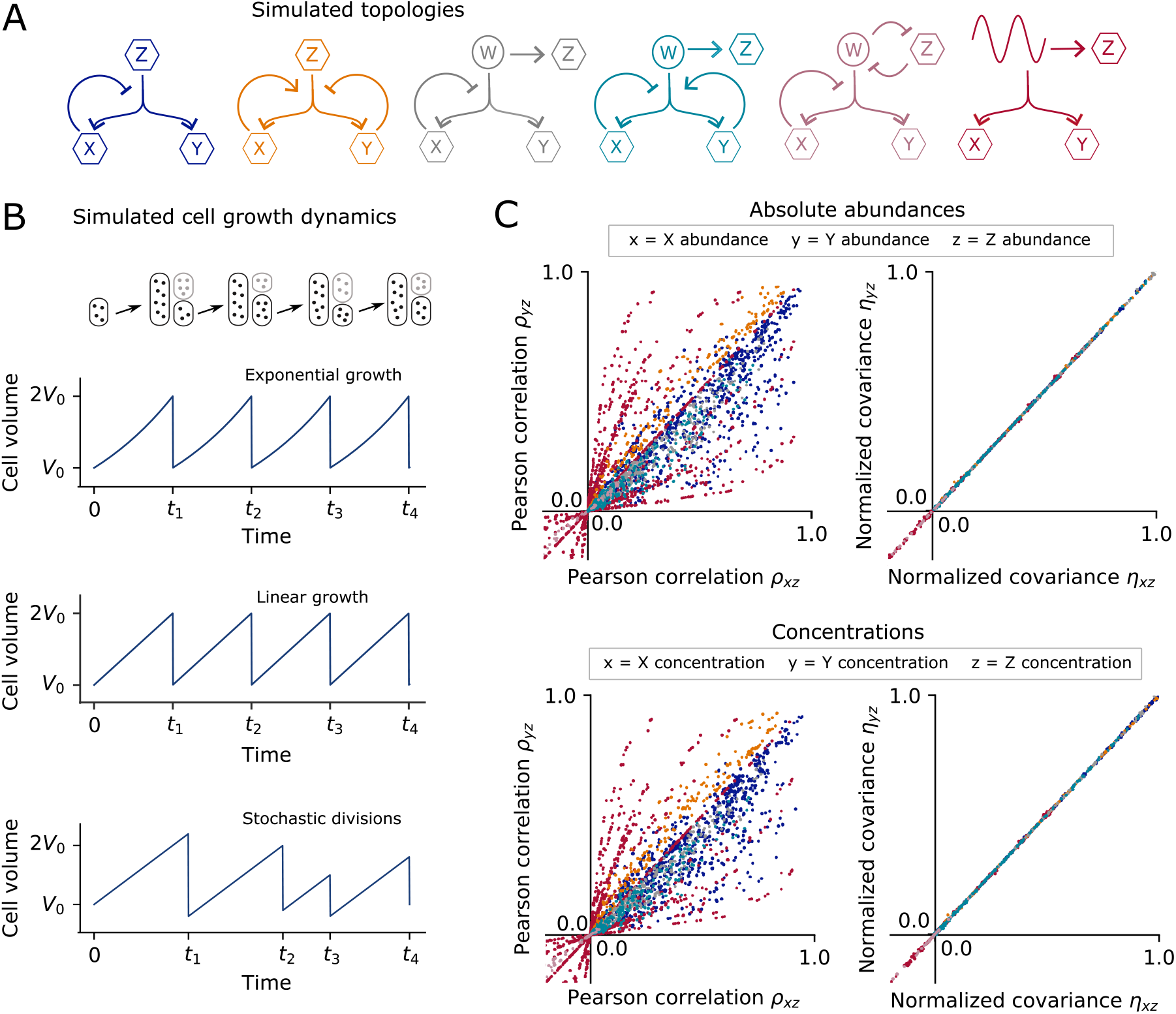
Numerical simulations of example systems confirm computationally that the analytically proven Eq. (2) constrains cellular abundances and concentrations in growing and dividing cells under fairly general assumptions as long as *X* does not affect *Z*. **A)** To numerically demonstrate the validity of Eq. (2) we consider ten example birth-death processes covering six different network topologies with non-linear rates, closed-loop feedback, time-varying upstream signals, and fluctuating degradation rates. In all cases chemical species *X* and *Y* are co-regulated but do not affect a third chemical species *Z* of interest. See supplemental Table S1 for details of the simulated systems. **B)** We consider three different cellular growth dynamics that affect molecular abundances through random partitioning of molecules at cell division and molecular concentrations through dilution. Additionally, system reaction rates depend in varying ways on the cell volume. At cell-division, molecular abundances are assumed to be partitioned on average proportional to cell size. **C)** Simulation results for each of the example topologies from panel A, subject to the different growth dynamics of panel B. Model parameters were varied over several orders of magnitude. The numerically obtained Pearson correlations do not satisfy a general relation whereas normalized covariances satisfy Eq. (2) for absolute abundances as well as concentrations. Colors correspond to the topologies indicated in panel A (results for individual topologies are compared in supplemental Figs. S3 and S4). For each dot, single-cell trajectories of 20,000 cell divisions were simulated at least 40 times, with the center of the dot corresponding to the average of the simulation ensemble (see Materials and Methods).

### Detecting causal interactions between genes through experimentally engineered “dual reporters”

Violation of Eq. (2) can be used to experimentally deduce the existence of a causal effect from a gene of interest *X* onto another gene of interest Z_*k*_ as follows. If we engineer a co-regulated “dual reporter” system with the properties defined in Eq. (1), in which *Y* is a passive reporter (i.e., it does not significantly interact with other cellular components) that responds to the same input as *X*, then a violation of Eq. (2) implies that changes in *X* must causally affect the abundance of component Z_*k*_. This is because intrinsic fluctuations from the expression of *X* propagate through the network and affect Z_*k*_, but those from the passive reporter *Y* do not. As detailed next, this approach can be applied to dual reporters for gene expression both on the transcriptional as well as the translational level, see Fig. 1.

#### Transcriptional dual reporters

Consider two genes of interest, *geneX* with transcript *X* and *geneZ* with transcript *Z*_*k*_. Our approach relies on engineering a passive reporter gene *geneY* with identical transcriptional control to *geneX* and mRNA lifetime, but with transcript that does not affect other cellular components, see Fig. 2A. This could be achieved with a transcriptional reporter that is put under control of the same promoter as *geneX* and placed at a similar gene locus. One way to ensure that *geneY* interacts minimally with other cellular components would be to remove its start codon so that it will not be translated into a protein. Co-regulated mRNA reporters have been engineered to satisfy the assumptions underlying Eq. (1), with transcripts counted with smFISH [15, 25, 26]. Alternatively, future improvements in RNAseq [27–29] accuracy might allow for reliable fluctuation measurements of transcript levels in single-cells through sequencing approaches.

#### Translational dual reporters

The dual reporter assay has been frequently implemented as co-regulated fluorescent proteins [14, 16, 22, 30]. The invariant of Eq. (2) directly applies when *X* and *Y* are co-regulated fluorescent proteins with first-order translation rates and maturation times (see SI Sec. 4 B for details of the proof). Fluorescent proteins can then be used to detect causal interactions in gene regulation as follows. The gene of interest *geneX* is fused to a fluorescent protein to make a functional fusion *geneX-FP*.

A passive reporter protein *Y* is made by introducing a spectrally distinguishable fluorescent protein *Y* under the control of the same (but distinct) promoter as *geneX-FP*. The expression level *z*_*k*_ of *geneZ* can be measured either through a transcriptional reporter or a functional fusion protein with a third fluorescence reporter. Under the assumption that the fluorescent protein *Y* does not directly affect other cellular components, Eq. (2) directly applies to the covariances of fluorescence levels as long as *X* does not causally affect *Z*_*k*_. As the normalized covariances in Eq. (2) are independent of scaling factors, standard fluorescence microscopy methods can be used without the need to determine absolute numbers. Two stable fluorescent proteins with similar maturation times [31] should be chosen to ensure that the assumptions underlying the class of systems is satisfied. Additionally, the translation rates of *X* and *Y* need not be identical, but can be proportional with a fluctuating proportionality factor as defined in the transitions of Eq. (1). As a result, different fluorescent proteins with different ribosome binding sites, mRNA secondary structures, and gene lengths, can be used as long as the translation rates remain proportional.

### Typical experimental single-cell data can be analyzed using the invariant relation of Eq. (2)

In the above sections we introduced the main mathematical result and presented the basic logic of how it can be experimentally exploited using synthetically engineered gene expression reporters. Next, we describe how Eq. (2) generalizes to the broader class of systems necessary to analyze single-cell experimental data.

#### Eq. (2) constrains abundances in growing & dividing cells

The class of stochastic processes presented in Eq. (1) constrains the dynamics of cellular abundances in stationary processes. However, experimental data typically report measurements of growing and dividing cells. Under the assumption that during cell division, molecular abundances are divided on average proportional to cell size, we can show (see SI) that Eq. (2) must be satisfied by the dual reporter abundances when *Z*_*k*_ is a component not affected by *geneX*. In this analysis of cellular growth and division, we allow for partitioning noise, division time fluctuations, asymmetric divisions, and arbitrary growth-rate dynamics. see Fig. 2B.

#### Eq. (2) constrains concentrations in growing & dividing cells

Chemical reaction rates typically depend on molecular concentrations and not absolute abundance numbers. Under general assumptions (see SI Sec. 12), we can show that the covariance constraint of Eq. (2) describes molecular concentrations in growing and dividing cells, see Fig. 2. This result assumes that the abundance of *X* does not affect cell volume or growth rate. This requirement can be intuitively understood, because if *X* affects cell volume then it causally affects the concentration of *Z*_*k*_ which depends on volume (see SI Sec. 12).

#### Eq. (2) applies to reporters with fluctuating degradation rates and fluctuating translation rates

The class of systems defined in Eq. (1) assumes that the dual reporters are degraded in a first-order process with a constant rate parameter *β*, see Eq. (1). This assumption can be relaxed: the invariant of Eq. (2) generalizes to the class of systems in which the degradation rate constant is an arbitrary function of all the cellular components that are not affected by the dual reporters, as long as it does not decay to zero (see SI Sec. 4). As a result, the degradation constant can vary in time with arbitrary extrinsic fluctuations. Additionally, for the class of co-regulated fluorescent proteins, the translation rate parameters can also vary with arbitrary extrinsic fluctuations.

#### A rule for detecting causality in the face of measurement uncertainty

Under the null hypothesis that there is no causal interaction from *X* to *Z*, Eq. (2) is equivalent to *r* = 1, where *r* = *η*_*xz*_*/η*_*yz*_ is the covariability ratio. Given 95% confidence intervals for an experimentally measured covariability ratio, we can conclude at the 2.5% significance level that there is a causal interaction from *X* to *Z* if *r* = 1 falls outside of the 95% confidence interval of the data (see Fig. S2 for an example)

#### In silico validation of analytical results

In order to illustrate the generality of the above results and to perform a numerical sanity check, we simulated several stochastic birth-death systems in growing and dividing cells, see Fig. 2. We modelled 10 systems made up of 6 network topologies in which a component *Z* is correlated with *X* and *Y* but is not affected by *X*, see Fig. 2A. These systems include cellular components with non-linear reaction rates and feedback loops, fluctuating degradation rates, and confounding variables that affect all observed components. We analyzed each system subject to three different cellular growth dynamics: periodic exponential growth, periodic linear growth, and linear growth with fluctuating cell-division times and division sizes, see Fig. 2B.

To numerically test Eq. (2), we generated random sample paths for abundances and concentrations using the Gillespie algorithm [32, 33]. System parameters were varied over several orders of magnitude. The numerical data verify that the normalized covariances of cellular abundances as well as cellular concentrations satisfy Eq. (2) in growing and dividing cells, see Fig. 2C. The same does not hold for the corresponding Pearson correlation coefficients, see Fig. 2C. Note, due to finite sampling, any numerical simulation will necessarily show small deviations from the exact equality of Eq. (2). Through statistical analyses and re-running systems for increased sampling, we confirmed that the (minuscule) deviations observed in numerical simulations were consistent with finite sampling, see Materials and Methods.

In SI Sec. 14 we simulate a larger example network made of 10 components in which *X* regulates a cascade of components. The results suggest that the relative distance down a regulatory cascade can be inferred from the degree of violation of Eq. (2), see Fig. S13.

#### Eq. (2) constrains data with significant measurement noise

Experimental techniques to measure mRNA abundances or fluorescence levels potentially introduce significant measurement noise. For example, when counting mRNA abundances, smFISH can lead to probabilistic undercounting noise and fluorescence microscopy can report fluorescence levels that includes photon noise, read noise, and segmentation errors, while flow cytometry introduces Poisson noise when used with bacteria [34].

The invariant of Eq. (2) holds in the face of measurement noise as long as the noise is symmetric in *X* and *Y* measurements. We can show (SI Sec. 16) that Eq. (2) holds in systems with arbitrary multiplicative noise that is independent of the *X* and *Y* signals, and additive noise that is independent of *Z*_*k*_. Additionally, the invariant holds in the face of systematic undercounting that introduces a binomial readout of the signal of interest, along with Poisson-Gaussian noise. If, however, the experiment introduces a different type of noise in *X* as compared to *Y*, then Eq. (2) is no longer valid. It is thus important to choose similar measurement techniques while measuring the abundances of *X* and its passive reporter *Y*.

### Experimental proof-of-principle

Whether the above theoretical approach works in practice depends on two crucial questions: Can we reliably build dual reporter systems that satisfy the assumptions underlying Eq. (2)? Are experimental violations of Eq. (2) larger than measurement uncertainties for current experimental techniques?

As a first test to address these questions we present an experimental proof-of-principle using synthetically engineered gene regulatory circuits in *E. coli*. These circuits consist of variants of the “repressilator”, a celebrated synthetic control circuit in which three genes respectively repress each other [35, 36].

#### Causally connected genes that break the invariant

Inherent to our fluctuation approach is the direction of inference, i.e., violations of Eq. (2) imply the presence of a causal connection, whereas agreement with Eq. (2) does not imply the absence of causal interactions. The question thus becomes whether a biologically relevant genetic circuit with known causal connections ever breaks our covariance relation.

We engineered four synthetic circuits in which a TetR-YFP fusion protein (*X*) represses an RFP reporter (*Z*), and a passive CFP reporter (*Y*) is under the same transcriptional control as *X*, see Fig. 3A. While the circuits differ in dynamics and connections, *X* affects *Z* in all cases. See SI Table. S2 for details of the synthetic circuits #1–4. Using time-lapse fluorescent microscopy with cells growing in a microfluidic device (see Fig. 3C), we measured the normalized covariances from populations of genetically identical cells with high accuracy while discarding all temporal information.

**Figure 3.**
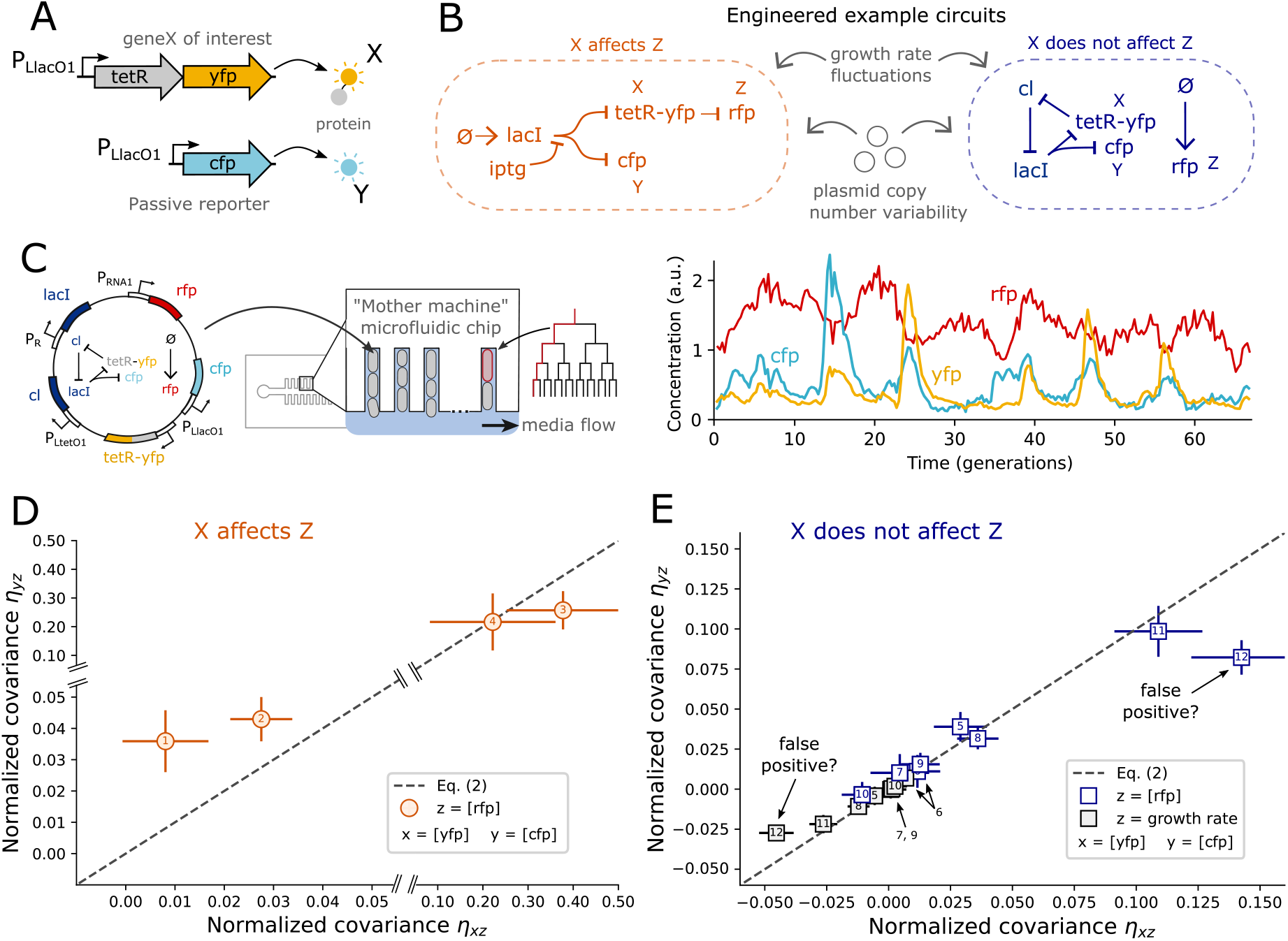
Experimental data from all but one synthetic regulatory circuit are consistent with the theory. **A)** In all synthetic circuits, we considered TetR as our protein of interest (*X*) fused to YFP to allow for quantification through fluorescence microscopy. CFP (*Y*) was used as a passive read-out of the transcriptional control of *tetR* by placing it under the control of a copy of the same *P*_*LlacO*1_ promoter as *tetR*. In all synthetic circuits *X* and *Y* are thus co-regulated by the LacI protein. **B)** We constructed two different types of synthetic circuits using the repressilator motif [35] as a basis. Left: example circuit in which TetR (*X*) causally affects a RFP reporter (*Z*). Right: negative control example circuit in which RFP was expressed constitutively and thus expected to be independent of TetR levels. In this circuit, *X* does not causally affect *Z* but *X* and *Z* are correlated due to plasmid copy number fluctuations. **C)** *E. coli* cells with the synthetic circuit encoded on the pSC101 plasmid were grown in a microfluidic device and observed over hundreds of cell divisions while daughter cells are washed away. Fluorescence levels of YFP, CFP, and RFP were measured simultaneously for hundreds of mother cells, along with cell area, cell length, and the growth rate. For each strain, the time-lapse data of all cells were combined into a population distribution from which the normalized covariances were computed. All temporal information was thus discarded and not used in the analysis. **D)** Two out of four causal interactions were clearly detected through violations of Eq. (2). Of the other two, one was clearly not detected, whereas one was situated right at the limit of experimental detectability but did not violate the null hypothesis test at a 2.5% significance level (see Fig. S2). See Materials and Methods for details of our error analysis. Numbers indicate the synthetic circuits as listed in supplemental Table S2. **E)** All but one negative control circuits led to data consistent with Eq. (2) (dashed line). Numbered synthetic circuits are listed in supplemental Tables S3 and S4. The two inconsistent data points corresponding to the same synthetic circuit (number 12) which is presented in detail in Fig. 4. The false positives either imply that our method works imperfectly or that we detected an unexpected causal interaction from TetR onto growth rate in this circuit. We present evidence for the latter interpretation in the next section.

We found that circuits #1 and #2 clearly broke the constraint, circuit #4 did not violate it, and circuit #3 showed significant deviations right at the limit of what can be reliably detected, see Fig. 3D. Using the estimated error bars and a null hypothesis test at the 2.5% significance level, we find that the data from circuit #3 does not reject the null hypothesis of Eq. (2), see Fig. S2.

The above repressilator test circuits were chosen for their well-characterized genetic interactions and because oscillating abundances make for a convenient test of the constructs. However, large oscillations in *Z* also make it much more challenging to detect violations of Eq. (2) because in those circuits the intrinsic noise we exploit is small compared to the periodically varying dynamics of the circuit. Indeed, the two clear violators correspond to altered versions of the repressilator circuit that do not oscillate (see SI Table. S2). These non-oscillating test cases may in fact be the most relevant assessment of using Eq. (2) in natural gene regulatory circuits. All the above circuits were made of genes encoded on a low copy number plasmid. We have not shown that our method can successfully detect causal interactions in a chromosomally encoded gene regulatory circuit. However, plasmid copy number fluctuations will reduce the degree of violation of Eq. (2) by reducing the fraction of reporter variability that is due to the intrinsic noise we exploit (see Sec. 15). A priori, causal connections between chromosomally encoded genes are thus expected to be easier rather than harder to detect with our method.

Overall the experimental data confirm that Eq. (2) has the fundamental power to experimentally detect causal interactions. However, the precision of current experimental techniques lies at the edge of what is necessary to detect physiologically relevant interactions when genes show large variability relative to the intrinsic noise we exploit. This highlights the importance of correctly estimating measurement uncertainty in our approach. Our error bars estimate sampling error and errors from non-even sample illumination, temporal drift, background fluorescence, and autofluorescence. These were estimated using a bootstrapping approach where the corrections and the normalized covariances were computed recursively to samples of the data, see Materials and Methods, and supplemental Fig. S6.

#### Negative controls

Our crucial theoretical result is that Eq. (2) must be satisfied by any circuit in which *X* does not affect *Z*. To experimentally test this prediction we engineered test cases in which *X* and *Y* are co-regulated, while *Z* is a component that is not affected by *X*. This took one of two forms: First, we constructed eight different circuits in which RFP (*Z*) was expressed constitutively using the pRNA1 promoter as previously used as a segmentation marker in *E. coli* [36, 37]. We used various synthetic circuits and the RFP reporter was either integrated chromosomally or included on the circuit plasmid (the latter leading to correlations between RFP and our synthetic components without introducing a causal interaction). Second, as an additional control, for each circuit we also considered the cellular growth rate as the unaffected cellular property *Z*. While the growth rate affects reporter concentrations through dilution, we did not expect cellular growth to be affected by our synthetic circuits which made growth rates a convenient additional negative control.

We found the data from all but one synthetic circuit consistent with our predicted invariant, see Fig. 3E. Note, the false positives correspond to the same synthetic circuit: one data point is from taking *Z* as the RFP concentration, while the other data point is from taking *Z* as the growth rate. To rule out experimental error we repeated the experiment twice while swapping the YFP and CFP reporters which confirmed the result (see Fig. S7). Additionally, we confirmed the result using a different cell segmentation pipeline (see Fig. S8).

Taken at face-value, our method thus produced two false positives (out of 16 tests), suggesting that our approach could be valuable but somewhat unreliable. However, next we present experimental evidence that indicate the violating data points were not false positives but may have correctly identified an unexpected causal effect from *X* on *Z* in the outlier circuit.

### Evidence that RpoS mediated stress response affected cellular growth in the outlier circuit

We observed that the outlier strain (Fig. 4A) exhibited periods of slow growth (Fig. 4B), suggesting that cells underwent periods of stress. Indeed, although RFP was thought to be constitutively expressed, RFP exhibited clear temporal dynamics that were negatively correlated with growth rate (Fig. 4C, Fig. S9). These fluctuations were larger than other strains expressing RFP from the same chromosomal gene (Fig. S10), suggesting that the outlier circuit caused additional fluctuations in the RFP concentration and growth rate.

**Figure 4.**
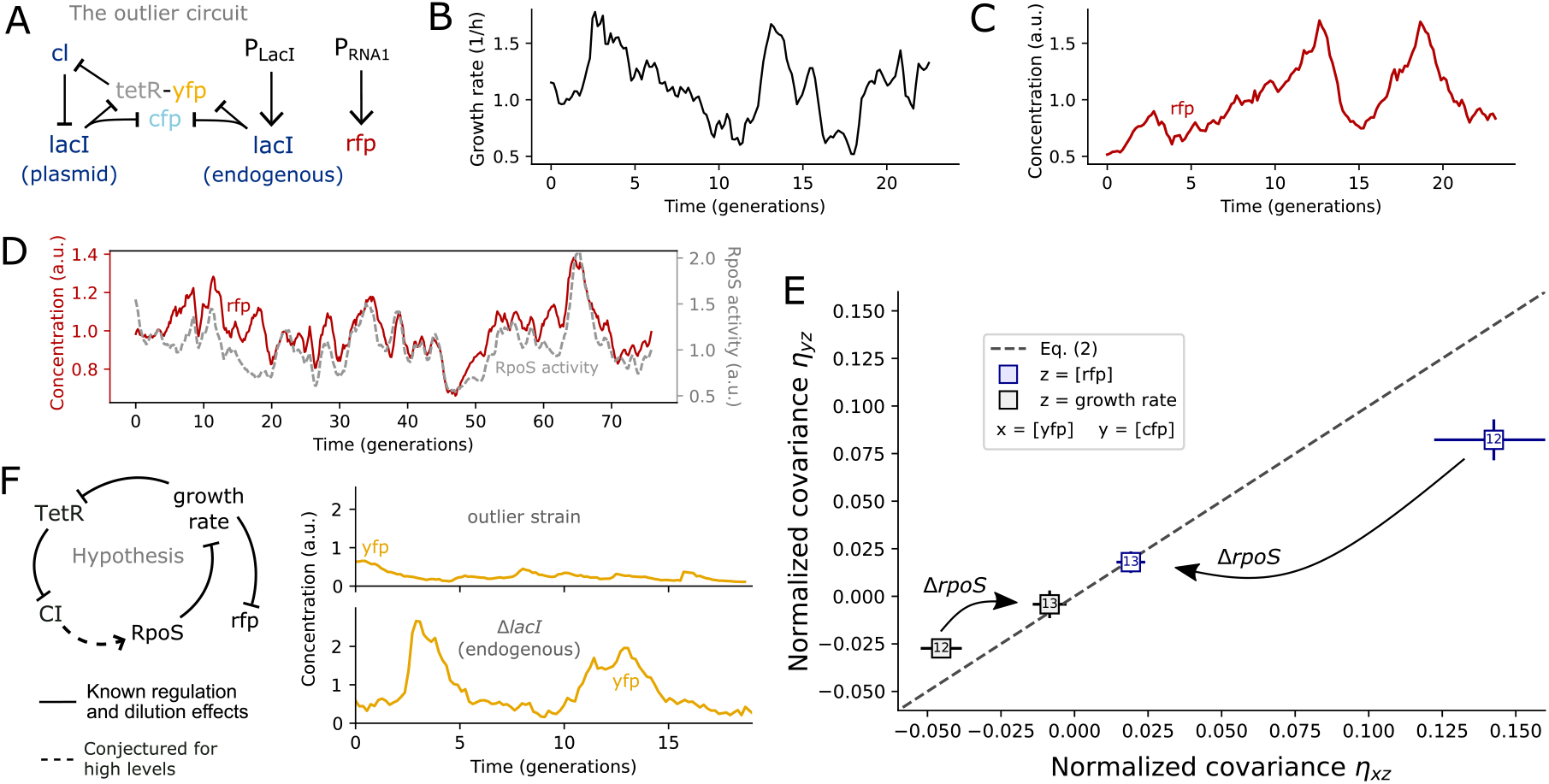
In the outlier strain the bacterial stress response is triggered which in turn regulates the “constitutive” RFP reporter. **A)** The outlier strain consists of the repressilator circuit encoded on the pSC101 plasmid with an endogenous copy of the *lacI* gene encoded on the chromosome. This endogenous source of LacI disrupts the regular oscillations of the repressilator circuit. RFP was chromosomally expressed under the control of the pRNA1 promoter [38]. **B)** The outlier strain exhibited large growth rate variability. **C)** The outlier strain exhibited large variability in RFP levels with the highest peaks occurring after the slowest growth periods. **D)** RFP levels were strongly correlated with RpoS activity as quantified through the known RpoS target gadX [39]. **E)** Deletion of the *rpoS* gene made the outlier strain consistent with Eq. (2). **F)** Expressing LacI endogenously leads to long periods of low TetR-YFP levels rather than regular repressilator oscillations. Because TetR is the only repressor of the *cI* gene in the synthetic circuit, we hypothesize that during periods of low TetR, CI expression is so high that resource competition with the also highly expressed RFP triggers the bacterial stress response. This interaction, hypothesized to be present in cells with high CI and RFP expression levels, is indicated with a dashed arrow. Solid arrows indicate interactions with direct experimental support [40].

We hypothesized that in the outlier strain the “constitutive” RFP reporter was being affected by the bacterial stress response regulator RpoS [41, 42]. By measuring the transcriptional activity of the known RpoS target *gadX* [39] we observed variable RpoS activity with pulses that correlated with periods of slow cellular growth (Fig. S11). Similar pulsing behaviour of RpoS triggering periods of slow growth has been reported previously [39, 40]. RFP levels were in turn strongly correlated with RpoS activity (Fig. 4D), as expected for genes for which transcription rates are constant but cell growth is slowed down by RpoS activity. Indeed, upon deleting RpoS, RFP fluctuations as well as growth rate fluctuations satisfy Eq. (2), see Fig. 4E.

Next, we discuss why the RpoS stress response was triggered in the outlier circuit in a TetR dependent manner. In the outlier strain, LacI was chromosomally expressed, which represses TetR independent of the repressilator circuit. Instead of regular oscillations of TetR, we observed extended periods of very low concentrations, see Fig. 4F. We conjecture that during those extended periods of low TetR levels, expression of CI is exceedingly high, which in combination with the already highly expressed RFP, leads to resource competition in cells that ultimately triggers RpoS mediated stress response. Note, CI is under the control of the very strong pLtetO1 promoter, which can initiate the transcription of up to 0.3 mRNA/sec in bacteria when fully induced [43] and has been shown to burden *E. coli* [44]. RFP is under the control of another strong promoter, pRNA1, previously used as a bright segmentation marker [36, 37], which we find to affect growth rate (Fig. S12). This RpoS mediated causal effect of TetR on RFP would explain the violations of Eq. (2) by the outlier circuit, why deletion of *rpoS* removes the outlier data, and why the same circuit without endogenous *lacI* (strain #5) did not violate the constraint, see Fig. 3E.

The disappearance of the deviation from Eq. (2) in the outlier circuit with RpoS knockout provides evidence that the outlier data may not have been a false positive, but detected an unexpected causal interaction mediated by cellular burden and RpoS.

## Discussion

A ubiquitous problem in understanding gene regulatory networks is identifying which genes regulate the expression levels of which other genes. Stochastic fluctuations of molecular abundances within cells provide a natural source of information about such regulation. Here, we presented a mathematical identity that potentially allows for the translation of such single-cell variability data into causal interactions. Because correlations do not imply causation, this method requires experimental intervention, but crucially it does not require perturbing the physiological state of the cells.

Note that agreement with Eq. (2) does not prove the non-existence of such a causal interaction because the lack of deviation can have two reasons: the causal interaction could be negligible or the effect of the intrinsic noise we exploit is not large enough compared with other sources of variability of the target gene. In other words, while deviations of Eq. (2) rigorously establish causal interactions, “absence of this evidence does not constitute evidence of absence” [45].

### Proposed additional tests

Our data from synthetic circuits in *E. coli* provide a proof-of-principle application in which our theoretical idea survived a first contact with experimental realities in specific synthetic circumstances. Synthetic gene circuits are useful for testing our network inference method because they provide reliably known interactions and can easily be modified to produce an array of circuits to test. However, synthetic circuits are limited in that they may not describe physiologically relevant interactions in naturally occurring systems. In addition, the preceding section suggests that synthetic circuits can put a metabolic burden on cells which can lead to unintended causal interactions. Further experiments are needed to demonstrate that the approach can detect causal interactions in natural gene regulatory circuits.

This requires applying the method on an endogenous network with reliably known natural interactions. For instance, the widely studied lac operon presents an exemplary test candidate: *X* would correspond to LacI, which represses the expression of *lacZ*. Using standard cloning techniques in molecular biology, the endogenous *lacI* and *lacZ* genes in a wild-type strain of *E. coli* can be replaced with *lacI-FP*_1_ *lacZ-FP*_2_ fusion genes, where *FP*_1_ and *FP*_2_ correspond to two fluorescent proteins of choice. A passive reporter *Y* would correspond to a spectrally distinguishable fluorescent reporter gene *FP*_3_ placed on the chromosome at similar gene loci to *geneX*. Violation of Eq. (2) would demonstrate that the method can detect the causal interaction between the LacI repressor and the lac operon expression. Additional negative control genes that are not regulated by LacI can then be studied to test the validity of Eq. (2) under broader conditions.

Note, that to minimize the metabolic burden of fluorescent proteins, the invariant Eq. (2) can be tested sequentially, i.e., doing the experiment with visible *X* and *Z* first, followed by an experiment to measure *Y* and *Z* next. If hybridization or sequencing techniques such as smFISH or scRNAseq are used to measure transcript abundances, then endogenous *geneX* and *geneZ* transcript abundances can be measured directly, and the only metabolic burden comes from the expression of the *geneY* reporter.

### Limitations of this study

Our method relies on a handful of key assumptions about the molecular reporters used. In the absence of standardized synthetic parts we may not know whether a given engineered system satisfies these assumptions. Only if the assumptions are satisfied, can violations of the invariant rigorously identify the existence of a directed causal interaction.

While the presented experimental study works as a proof-of-principle to illustrate the approach and establish its practical feasibility, at face value it also produced a false positive result. Through additional perturbation experiments we argue that the outlier circuit includes an unexpected causal connection. However, the evidence for this conclusion is weaker than the proven theoretical results presented. The fact that our interpretation of the perturbation experiments will be debated may serve as an illustration of how valuable a fluctuation based approach would be that avoids perturbation experiments altogether.

The negative control circuits in this study were chosen because they lacked a causal interaction from *X* to *Z* when considering the known interactions between the used promoters and proteins. However, the false positive result suggests that unexpected causal interactions can still exist in these circuits. Further perturbation experiments and additional controls would thus be beneficial to directly verify that *X* did not affect *Z* in our synthetic negative control circuits.

Additionally, all experimental synthetic circuits were tested in *E. coli* and eukaryotic test cases remain to be investigated. In eukaryotic cells, promoter sequences may not be the sole factor determining transcription rates. For instance, dual reporter genes with identical promoters have been engineered in mammalian cells [25]. The expression of these genes underwent bursts that are not coordinated when the genes are placed at distant gene loci [25]. While the invariant of Eq. (2) holds in the face of such stochastic bursting, it only holds when, on average, the burst frequency is the same for each dual reporter (see SI Sec. 8). Additional consistency checks should also be reported to test whether the passive reporter reads out the same signal as the gene of interest. This could be done, for example, by testing the invariant of Eq. (2) with other cellular components that are known to not be affected by *geneX*, or by direct measurement of the transcription activity using a separate method [46]. However, it is possible that the existence of some of the many other factors of gene regulation in eukaryotes, such as the presence of enhancers or epigenetic modifications, could make the proposed approach difficult to apply. It thus remains to be shown that a passive reporter can be engineered to read-out the same signal as a naturally occurring gene of interest in the face of additional factors of gene regulation in eukaryotes.

## Materials and Methods

### Data and materials availability

Numerical simulation code and data, code for analyzing the single-cell traces to compute the normalized covariances, python code used to run the DeLTA deep learning segmentation pipeline, as well as plasmid sequences, are all available on Github (https://github.com/ejolysmith/Exploitingfluctuations-causal-interactions-manuscript). The segmented and tracked single-cell traces along with our trained U-Net models used for segmentation and chamber identification are available on Zenodo (https://doi.org/10.5281/zenodo.15616830).

### Numerical simulation details

Exact simulated single-cell time trajectories of the abundances were generated using the standard Gillespie algorithm [32], with an additional step to account for time-dependent rates and divisions [33] (see SI Sec. 13 for the exact algorithm used). The time trajectories for the concentrations correspond to the abundance trajectories divided by the cell volume trajectories *V* (*t*), with the latter being simulated independently of the species abundances. Simulations were performed with python.

#### Simulated cell growth and division models

Three cellular growth dynamics were simulated. The first and second are linear and exponential growth respectively, with constant division times and symmetric cell divisions. Here *V* (*t*) trajectories were generated analytically as periodic functions. The volume is reduced by factor 1*/*2 at evenly spaced division times {*τ*_*i*_}. Between division times *τ*_*i*_, *τ*_*i*+1_ the volume is given by 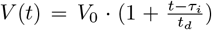 for the linear case and 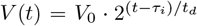 for the exponential case, where *t*_*d*_ is the time between divisions, and *V*_0_ is the volume right after a division. The third simulated volume dynamics is linear growth with stochastic division times and asymmetric divisions. Here a constant linear growth rate 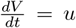 is used. Division times {*τ*_*i*_} and division factors {*a*_*i*_} are picked recursively: The *τ*_*i*+1_ is taken from a normal distribution with mean set to the doubling time:

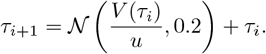

where 𝒩 (*µ, σ*) is the normal distribution with mean *µ* and standard deviation *σ*. Until then the volume grows linearly with constant rate

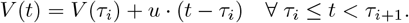

At *τ*_*i*+1_ the volume is reduced by factor *a*_*i*+1_ taken from a normal distribution with mean set to the ratio of cell volumes at the beginning and end of the cycle

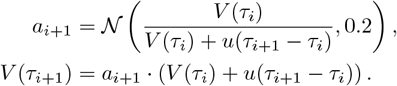

As a result, cells that grow more (less) than double in size during a cycle tend to divide with larger (smaller) division factors, ensuring the volume trajectories do not eventually expand (decay) to infinity (zero).

At division, molecular abundances are reduced according to a binomial splitting with probability given by the division factor. For example, if the volume is reduced by a factor of 0.4 at a cell division, i.e., *V →* 0.4 *V*, then each molecular has probability 0.4 to remain in the followed daughter cell.

#### Computing normalized covariances from trajectories

Normalized covariances were computed by integrating over the trajectories to obtain time averages for first and second moments. This is equivalent to using the distribution given by calculating the fraction of the total system time spent in each sampled state. In the ergodic regime, this distribution converges to the stationary distribution of the ensemble.

#### Simulated systems

We simulated ten groups of systems defined by their rate functions (see SI Table. S1 for details). For each group, each model parameter was picked randomly multiple times from the set {0.1, 1, 10}. This was done for each of the the three volume dynamics we considered. If a simulated system gave an average abundance in one of the components less than 0.01 molecules, then the system was omitted to avoid numerical errors that arise from divisions of small numbers when computing the normalized covariances. In total, there are 6522 resulting simulations which are plotted in Fig. 2C.

#### Confidence intervals for finite sampling error

For each system, simulated trajectories were generated forty times, and then repeated until the percentage errors of the normalized covariances, taken as the standard error of the mean divided by the mean, reached less than 1% or the number of simulations reached 1,000. Each trajectory ran for 20,000 cell divisions and started with a unique random number generator seed. Each component abundance was set to 1 at the beginning of each trajectory, and the cell cycle time set to 0. We let the simulated cells reach 200 cell divisions before we start to compute the time average integrals for the moments, in order for the effect of the initial condition to dissipate and we analyze the systems’ cyclo-stationarity state. The final normalized covariance for each system corresponds to the average taken over the ensemble of simulated trajectories with 95% confidence intervals given by twice the standard error. Distributions for the uncertainties for each class of systems are plotted in supplemental Figs. S3, S4.

For a given system, Eq. (2) was tested by verifying that the ratio *η*_*xz*_*/η*_*yz*_ produced a 95% confidence interval that encompassed the predicted value of 1. The test was satisfied by 6369 out of 6522 systems (97.7%), consistent with the definition of confidence intervals for finite sampling. The percentage of outliers for each simulated process is plotted in supplemental Fig. S5. Re-simulating the outliers for twice the number of simulations led to a reduction in the standard error by a factor of 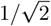, with Eq. (2) being satisfied by 143 of the 153 outliers (93.5%). Re-simulating the remaining 10 violators with four times the number of simulations, reduced the errors by a factor of 1/2, and all of the 10 outliers satisfied Eq. (2) when quantified at such high numerical accuracy.

### Strain and plasmid construction

All strains, plasmids, full construction details are provided in the SI. The base background strain used throughout the manuscript is *E. coli* MG1655.

The base plasmid used throughout the manuscript is the repressilator [35, 36]. It consists of three genes: *tetR* from the Tn10 transposon, *cI* from bacteriophage *λ*, and *lacI* from the lactose operon, which have promoters that are repressed by LacI, TetR, and CI respectively. All genes are placed on the low-copy pSC101 plasmid which is inserted into *E. coli* MG1655.

Using standard molecular biology techniques, the repressilator plasmid sequence was altered in a number of ways to construct the different circuits. The *tetR* repressor was set as the gene of interest *geneX* and TetR levels were measured through fluorescence measurements of a TetR fusion to the yellow fluorescent protein *mVenus NB* (YFP) [31, 47]. We engineered the passive reporter *geneY* by expressing a different fluorescent protein *SCFP3A* (CFP) [31, 48] under the control of the same pLacO1 promoter as *geneX* on the same pSC101 plasmid. Both *X* and *Y* levels are co-regulated by LacI concentrations. To ensure equal degradation times, we used a version of the repressilator in which the three repressing genes lack degradation tags so that circuit proteins are removed predominantly from cell-division and dilution [36]. The fluorescent proteins *mVenus NB* and *SCFP3A* were chosen since they both have a short maturation half-life (4.1 *±* 0.3min and 6.6 *±* 0.5min respectively at 37^*o*^C) compared to the time-scale of the repressilator oscillations (*∼*5 hours).

For the RFP, we used a modified *mkate2* hybrid (with improved translational efficiency) throughout all experiments, which consists of the *mCherry* N-terminal 11 amino acids followed by the *mKate2* sequence. In the experiment shown in Fig. 4D in which GFP was used to measure *gadX* expression, we used *gfpmut2* as a transcriptional reporter for the *gadX* promoter, placed on a low-copy pSC101 plasmid (taken from the *E. coli* promoter library [49]). For Fig. 4E, we used a strain of *E. coli* MG1655 with *rpoS* deletion taken from the *E. coli* Keio Knockouts library [50].

We chose the repressilator as the base circuit because the circuit interactions are known and the resulting dynamics (periodic oscillations) can be used as a consistency check to ensure that the TetR-YFP fusion is functional (i.e., still represses the *λ* gene).

### Mother machine experiment

We used a microfluidic device commonly called “mother machine” to follow growing and dividing cells for hundreds of generations in controlled environments [51, 52]. Single-cells are trapped in micrometer wide trenches. As cells grow and divide, the mother cells remain trapped while newborn daughter cells are washed away by the constant flow of growth media. Automated time-lapse microscopy and cell segmentation software enables us to track hundreds of cells in each experiment, while precisely measuring cell fluorescence, growth rate, and cell size. The time-series data for each mother cell can be pooled into a distribution from which the normalized covariances can be computed.

Though the invariant of Eq. (2) does not require the use of time-series data, we used the additional temporal information as a consistency check for our assumptions. In particular, the highly correlated trajectories of the YFP and CFP fluorescence (see Tables. S3 and S4) are consistent with our assumption that these reporters are co-regulated. Additionally, the observed oscillations in the closed loop circuits are indicative that the TetR-YFP fusion has not lost its function as a result of the added fluorescent protein. Moreover, the time-series data allows us to measure cellular growth rates which can be used to test Eq. (2) as shown in Fig. (3)D.

#### Microfluidic chip preparation

Polydimethylsiloxane (PDMS) (Sylgard 184 Silicon Elastomer, Fisher Scientific) was mixed at a 10:1 (monomer:curing agent) ratio, poured on top of a 1.0 *µ*m tall wafer and degassed for one hour at room temperature before baking for an additional 1.5 hours at 65^°^C. After careful removal of the PDMS from the wafer, individual PDMS chips were cut out with a razor blade. The inlet and outlet holes were punched with a 0.75 mm biopsy puncher (World Precision Instruments). The PDMS chips were sonicated in isopropyl alcohol (Fisher Scientific) for 30 minutes and dried at 65^°^C for 15 minutes. Glass coverslips (Fisher Scientific: 22×40 mm #1.5) were cleaned with 1M potassium hydroxide (KOH, Sigma Aldrich) for 20 minutes. The PDMS chips were bonded to the glass coverslips using a plasma cleaner (Oxygen flow rate at 45 sccm, power at 30W for 15 seconds, Tergeo Plasma Cleaner, PIE Scientific). The completed microfluidic device was heated to reinforce the plasma bonding at 100^°^C for 10 minutes, then 65^°^C for 30 minutes.

#### Cell preparation

*E. coli* strains were grown overnight in LB with appropriate antibiotics to select for cells containing the constructed plasmids. At *∼*3-4 hours prior to the experiment start time, the overnight cultures were diluted 1:100 in imaging media consisting of M9 salts, 10% (v/v) LB, 0.2% (w/v) glucose, 2 mM MgSO4, 0.1 mM CaCl2, 1.5 *µ*M thiamine hydrochloride and 0.85 g/L Pluronic F-108 (Sigma Aldrich, surfactant to prevent cells sticking to the surface of a microfluidic device). At an OD600 of 0.2-0.4, cells were loaded into the main feeding channel of the microfluidic chip and centrifuged at 5000 g for 10 min to push cells into the cell trenches. The feeding channels were then connected to syringes filled with imaging media using Tygon tubing, and the microfluidic chip was placed in a temperature controlled incubation chamber set at 37^°^C. Media was pumped through the feeding channel using syringe pumps (New Era Pump System), first at a rate of 5 *µ*L/min for *∼*0.5-1 hours to get the cells comfortable in the trenches, then at a high rate of 100 *µ*L/min for 1 hour to clear the inlets and outlets. The rate of media flow was then set back to 5 *µ*L/min for the duration of the experiment.

#### Microscopy and image acquisition

Images were acquired using a Zeiss Axio Observer inverted microscope equipped with a 63x Plan-Apochromat M27 oil objective (NA 1.40), an Orca Flash 4.0 LT camera (Hamamatsu), and an LED epifluorescence illuminator (Zeiss Colibri 7). The experiments were performed inside a temperature controlled incubation chamber set at 37^°^C. To reduce photobleaching the exposure time (100 ms) and light intensity (10-20%) were set low, with 16-bit CZI images taken every 5-8 minutes. Focal drift was corrected automatically with the Definite Focus 2 (Zeiss) monitoring and compensation system using an infrared laser (850 nm). In all experiments in which CFP, YFP, and RFP are measured simultaneously (i.e., all experiments except for the one shown in Fig. 4D), the following filter sets were used for acquisition: CFP (Semrock FF02-475/20-25), YFP (Chroma CT560/39bp), RFP (Zeiss Filter Set 91 HE LED), with respective Zeiss Colibri 7 illumination wavelengths set to 430nm (CFP), 511 nm (YFP), and 590 nm (RFP), along with a the Zeiss TBS 450/538/610 beam splitter. For the experiment in Fig. 4D in which a gfp reporter is used to measure gadX expression, the following filter sets were used for acquisition: GFP (Chroma ET525/50m) and cy5 (Zeiss BP 690/50) with Zeiss Colibri 7 illumination wavelengths set to 475 nm (GFP) and 630 nm (cy5), along with Zeiss QBS 405/493/575/653 beam splitter.

### Data analysis

#### Segmentation

Because all three fluorescence channels were used to measure synthetic circuit components throughout, we used the bright-field channel for segmentation. This was achieved with the automated deep learning-based cell segmentation software DeLTA [53, 54]. The U-Net deep learning models used for channel identification and cell segmentation were trained with a large dataset built from 5 mother machine experiments performed in the Potvin Lab that were generated with the same microscopy and image acquisition setup. In these training datasets a bright RFP is expressed and a thresholding segmentation pipeline is performed on the RFP channel images to obtain segmentation masks. The U-Net deep learning models from DeLTA are trained with these segmentation masks in combination with bright-field images from the training dataset as described in [53, 54]. The training dataset was segmented with the same thresholding pipeline used in previously described procedures [36, 55]. We also trained a DelTA model to use the RFP channel for segmentation and obtained similar results on the strains with bright RFP fluorescence. In the SI we include videos of a mother machine with segmentation boundaries obtained from our DeLTA models, along with the single-cell growth trajectories.

We estimated the YFP, CFP, and RFP single-cell concentrations as the average fluorescence intensities of all pixels in the segmentation mask of each cell. Cell area is measured as returned by opencv’s contourArea() over the segmentation mask. Cell length is measured by fitting a rotated bounding box to the segmented cell. We only analyzed data from the mother cells trapped at the top of the growth chambers.

#### Single-cell traces construction

For a given cell chamber with the Region of Interest identified by the U-Net model, a single-cell trace was constructed by selecting the segmented cell in each frame with highest average vertical pixel location. This in effect keeps track of the mother cells trapped at the top of the growth chambers.

Each single-cell trace went through a manual purging process. Traces with cell areas that are not growing and dividing were removed, as they are expected to correspond to dead cells. Traces with growing and dividing cell areas with very low constant fluorescence correspond to cells that lost the inserted plasmid due to random partitioning at cell division. These traces were also purged, but were analyzed separately to measure the cell autofluorescence as described in the next section. Moreover, segmentation and tracking errors were reduced by purging any parts of the traces that exhibited a clear non-biological anomaly.

Cell divisions were identified by a sudden decrease in the cell area. A division is called whenever the cell lengths dropped to less than 80% of its previous value.

#### Temporal drift correction

Focal drift was reduced automatically with a 850 nm infrared laser (Zeiss Definite Focus 2). However, we used an oil objective, which can cause temporal drift from spreading of the oil over time. We thus applied an additional correction to the data as follows.

In a given mother machine experiment, we computed the mean YFP, CFP, and RFP signals of all the cells at each time frame. For example, the mean YFP time-trace is given by

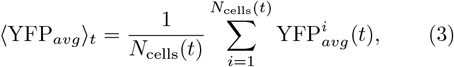

where 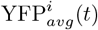 is the average YFP intensity taken over the segmented area of the *i*^*th*^ cell at time frame *t*, and *N*_cells_(*t*) is the number of surviving cells at time frame *t*. The mean time-traces of each fluorophore are fitted with a second degree polynomial according to a least-squared fit. To correct for the drift, we multiply a measured intensity with the reciprocal of the respective fitted polynomial (see Fig. S13). For example, if *f* (*t*) is the obtained fit of YFP_*avg t*_ from Eq. (3), we correct all the YFP_*avg*_ measurements as follows

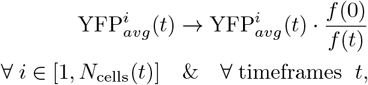

where *f* (0) is the fitted function *f* (*t*) taken over the first time frame.

#### Uneven illumination and background correction

A single image frame from our setup encompasses *∼*30 cell trenches with total spatial extension of *∼*100 *µ*m. This results in uneven illumination onto the cells. We use the linear gain model [56, 57] to correct for uneven illumination and background fluorescence:

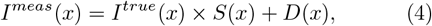

where *I*^*m*^(*x*) and *I*^*true*^(*x*) are the measured and true intensities at horizontal pixel position *x* respectively, the multiplicative term *S*(*x*) models the uneven illumination, and the additive term *D*(*x*) is any background noise that is present when no light is incident on the sensor. For our imaging setup we decompose *I*^*true*^(*x*) as follows

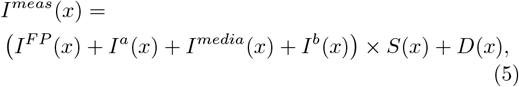

where *I*^*F P*^ (*x*) is the intensity from the fluorescent proteins, *I*^*a*^ is the autofluorescence of the cell, *I*^*media*^ is any fluorescence originating from the media, and *I*^*b*^ is any fluorescence originating from the PDMS background of the microfluidic device. The total background is given by *b*(*x*) = *I*^*b*^(*x*) *× S*(*x*) + *D*(*x*), in which case the imaging model becomes

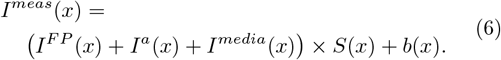

We use a ‘prospective’ approach to correct for *S*(*x*) and *b*(*x*) as follows. For each segmented cell in an image, we compute the average intensity in a box of dimensions 50 by 50 pixels located 15 pixels above the cell trench (see Fig. S15A). This in effect estimates *b*(*x*) at the *x* position of just above each cell trench. In each image, this *b*(*x*) is removed from the *I*^*meas*^(*x*) measurements of each segmented cell. To estimate the uneven illumination gain function *S*(*x*) we pool all of the single cell fluorescence measurements according to their horizontal pixel position *x*. Data is binned into bins of size 50 pixels and averages are taken at each bin (see Fig. S15B). The resulting curve gives an estimate of *I*^*F P*^ + *I*^*af*^ + *I*^*media*^ *× S*(*x*). To estimate *S*(*x*) we fit the curve with a 3rd degree polynomial *p*_*S*_(*x*). To correct for the uneven illumination gain function we multiply the fluorescence measurements by the reciprocal of *p*_*S*_(*x*):

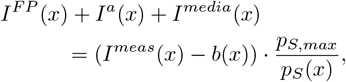

where *p*_*S,max*_ is the max value of the fit *p*_*S*_(*x*) over *x*.

#### Autofluorescence and fluorescing media

The preceding section described how the background and the uneven illumination were corrected using the linear gain imaging model of Eq. (6). Even with the background correction, the fluorescence profile across growth chambers empty of cells is not negligible compared to the profiles of some cell-containing chambers, see Fig. S15. This indicates that media fluorescence is not negligible in experiments with strains that produce low levels of fluorescence. Here we show how we estimated the autofluorescence *I*^*a*^ and the media fluorescence *I*^*media*^ to obtain the sought after fluorescent protein fluorescence *I*^*FP*^.

In each mother machine experiment there was a subset of cells that lost the synthetic circuit plasmid due to random partitioning at cell division. These cells were used to estimate *I*^*a*^ + *I*^*media*^. Cells that lost the plasmid were identified manually by observing single-cell traces: When a cell loses the plasmid, the CFP and YFP fluorescence decay rapidly and reach a constant low signal level for the remainder of the experiment, see Fig. S14. The segments of the time traces at constant low signal level were cut and saved, with the autofluorescence and the media fluorescence taken as the average fluorescence of the saved traces. Around 10–30 cases of plasmid loss occur for each strain in a mother machine experiment.

Note that *I*^*a*^ and *I*^*media*^ do not need to be corrected in the RFP channel measurements. This is because RFP is taken as the component either affected or not affected by *X*. For the systems in which RFP is not affected by *X*, any *function* of RFP (like adding autofluorescence and media fluorescence) will also not be affected by *X* and the invariant of Eq. (2) will be satisfied. Alternatively, if *X* affects RFP, then it will generally also affect any typically expected function of RFP. We did not correct for *I*^*a*^ and *I*^*media*^ in the RFP channel because a fraction of circuits have RFP located on the chromosome and not the plasmid.

#### Estimating confidence intervals

The corrections from the preceding sections rely on the distribution of single-cell measurements. As a result, sampling error affects the accuracy of the corrections, along with the final estimators of the normalized covariances. We applied two methods to estimate normalized covariance confidence intervals by taking into account sampling error and its effect on the corrections.

First we use bootstrapping, where the data corrections and the normalized covariances are computed over many samples of the data, allowing for replacement in sampling. If an experiment produces *N* single-cell traces of cells with plasmid, and *M* single-cell traces of cells that lost the plasmid, random samples of size *N* and *M* of each respective ensemble are taken. The corrections from the previous sections are then applied using the samples, with the corrected data pooled into a distribution from which the normalized covariances are computed. This is repeated until 100 normalized covariances *η*_*sample*_ have been computed for each sample. The final reported normalized covariance corresponds to the average over the *η*_*sample*_, with confidence intervals given by twice the standard deviation. See Fig. S16 and SI Sec. 17 for details.

Second we use sampling without replacement, where the single-cell traces from each experiment are divided into 7 to 10 disjoint sets, and the corrections and the normalized covariances are computed for each set. The reported normalized covariances correspond to the average of those computed from the sets, with error bars being the standard error of the mean (see SI Sec. 17 for details).

A single mother machine experiment typically produced 100–500 single-cell traces of cells with plasmid, and 7–30 single-cell traces of cells that lost the plasmid due to random partitioning at division. The computed normalized covariances from the bootstrapping method are shown in Fig. 3D,E, while those from the alternative splitting method shown in Fig. S6. The two methods give similar results.

### Growth rate estimation

We define growth rate in this work as the rate of change of the cell length normalized by cell length: 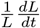. To estimate the growth rate from the single-cell traces we assume exponential growth between division events.

In that case, if there is no division event that occurs at time frames *i*−1, *i*, and *i*+1, then the growth rate at time *t*_*i*_ is computed as 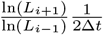, where *L*_*i*_ is the cell length at time *t*_*i*_, *L*_*i*−1_ is the length at the preceding frame, *L*_*i*+1_ is the length at the subsequent frame, and Δ*t* is the time between frames (5–8 min). If there is a division event at the subsequent frame *i* + 1, then the growth rate is computed as 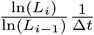. If there is a division event at the current frame *i*, then the growth rate is computed as 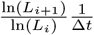.

For illustration we smoothed the growth rate measurements shown in Fig. 4B using a moving average filter with a window of 5 frames. We did not smooth the data when computing the normalized covariances and CVs shown in Fig. 3.

## Acknowledgments

We thank Raymond Fan, B Kell, Seshu Iyengar, Sid Goyal, and Josh Milstein for many helpful discussions. This work was supported by the Natural Sciences and Engineering Research Council of Canada and a New Researcher Award from the University of Toronto Connaught Fund. Simulations were performed on the Niagara, Beluga, and Narval supercomputers at the SciNet HPC Consortium. SciNet is funded by the Canada Foundation for Innovation; the Government of Ontario, Ontario Research Fund - Research Excellence, and the University of Toronto. E.J.S. gratefully acknowledges a doctoral research visit grant to support their experimental work in L.P.T.’s lab at Concordia University.

## Author Contributions

E.J.S. derived the analytical results. M.M.T and E.J.S. performed the numerical simulations. E.J.S. and P.A. carried out the experiments. E.J.S. and F.P. constructed the synthetic circuits. E.J.S. performed data analysis. E.J.S., L.P.T., and A.H. designed the study. E.J.S. and A.H. wrote the manuscript with input from all authors.

