## Supplemental Information for "Exploiting fluctuations in gene expression to detect causal interactions between genes"

Euan Joly-Smith,[1,2] Mir Mikdad Talpur,[1] Paige Allard,[3] Fotini Papazotos,[3] Laurent Potvin-Trottier,[3,4,5] and Andreas Hilfinger[1,2,6,7]

[1]Department of Physics, University of Toronto, Toronto, Canada
[2]Department of Chemical & Physical Sciences, University of Toronto, Mississauga, Canada
[3]Centre for Applied Synthetic Biology, Concordia University, Montréal, Québec, Canada
[4]Department of Biology, Concordia University, Montréal, Québec, Canada
[5]Department of Physics, Concordia University, Montréal, Québec, Canada
[6]Department of Mathematics, University of Toronto, Toronto, Canada
[7]Department of Cell & Systems Biology, University of Toronto, Toronto, Canada

### Contents


1. Supplementary figures 1

2. Supplementary tables 18

3. Cloning details 24
   A. Bacterial strains and growth conditions 24
   B. Plasmid construction 24

4. Class of systems 25
   A. mRNA reporters 26
   B. Fluorescent protein reporters 26

5. The invariant relation — Derivation of Eq. (2) 27

6. Derivation of the invariant relation for fluorescent proteins 30

7. The X and Y production rates can differ by certain types of fluctuations 30

8. Bursty gene expression 33

9. Simulated example systems from Fig. 1B 34

10. Example system with feedback from Fig. S1 34

11. Details on the models simulated in Fig. 2 35

12. Derivation of Eq. (2) in growing and dividing cells 38
    A. Abundances 38
    B. Concentrations 39

13. Simulation algorithm for growing and dividing cells 40

14. Simulation of a 10 component system with a regulatory cascade 41

15. Fluctuations in plasmid copy numbers can reduce the degree of violation of Eq. (2) 43

16. The invariant relation holds in the face of measurement noise 43
    A. Multiplicative noise 43
    B. Additive noise 44
    C. Binomial readout and undercounting noise 45
    D. Poisson-Gaussian noise model 45
    E. Segmentation noise 46

17. Estimating confidence intervals 47
    A. Sampling with replacement (bootstrapping) 47
    B. Sampling without replacement (splitting data) 47

References 48

### 1. Supplementary figures

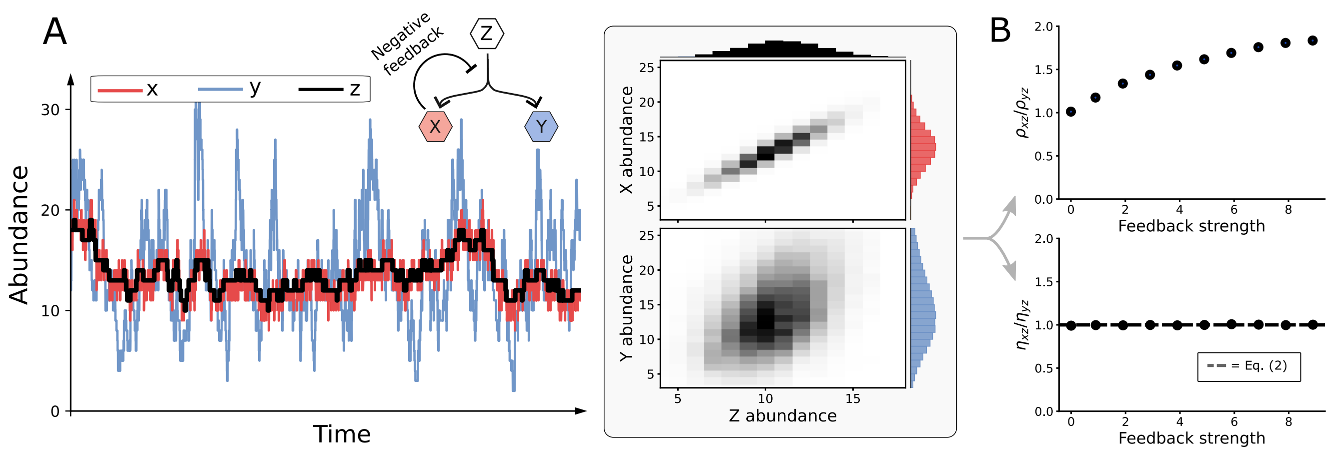


Figure S1. **The dynamics of the "dual reporters" $X$ and $Y$ are not necessarily symmetric despite the fact that $X$ and $Y$ have the same production rate. A)** Feedback in gene regulation can suppress or amplify fluctuations in the abundances of molecular species. Here we show an example system of the class of Eq. (1), where $X$ affects the shared transcription rate of $X$ and $Y$, but does not affect the shared upstream variable $Z$. In this example, the intrinsic fluctuations of $X$ are suppressed through negative feedback, whereas those from $Y$ are not. This leads to unequal dynamics and cell-to-cell variability in $X$ and $Y$ levels. See Sec. 10 for details of the simulated example system. **B)** The invariant Eq. (2) does not hold in terms of Pearson correlation coefficients, i.e., $\rho_{xz} \neq \rho_{yz}$ even when $X$ does not affect $Z$. In this example system, the divergence between $\rho_{xz}$ and $\rho_{yz}$ increases as the feedback strength increases (see Sec. 10). In contrast, Eq. (2) constrains all systems in which $X$ does not affect $Z$, even when the $X$ and $Y$ levels have vastly different dynamics and cell-to-cell variability. Note, the covariances and correlations can be determined from the static snapshots of cell-to-cell variability indicated in panel A. The time-trace to indicate the *dynamics* of the reporters is for illustration purposes only.

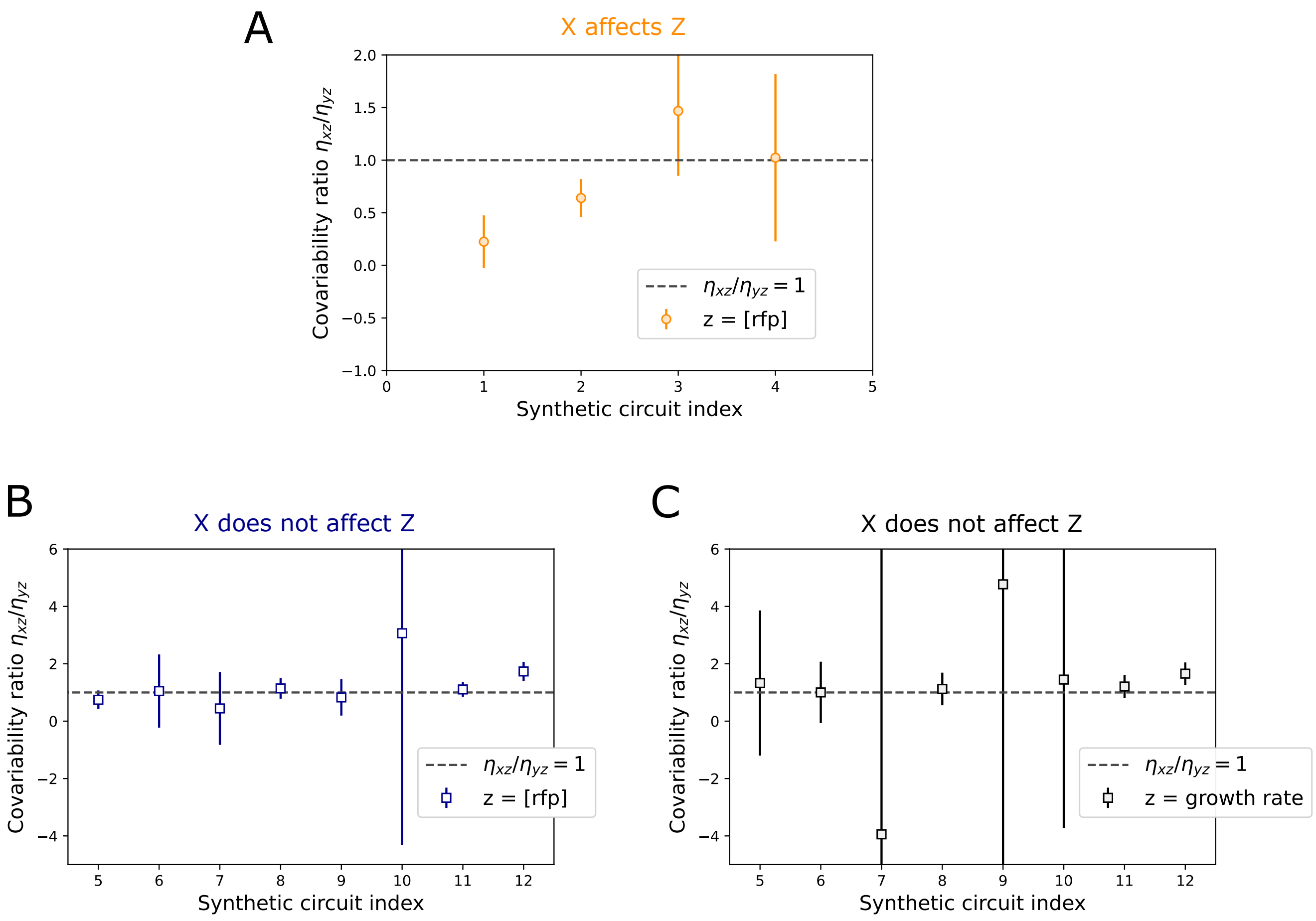


Figure S2. **Using the null hypothesis test on the measured covariability ratios. A)** Plotted are the measured covariability ratios $\eta_{xz}/\eta_{yz}$ for synthetic circuits shown in Fig. 3D and Table. S2 in which $X$ affects $Z$. Error bars correspond to estimated 95% confidence intervals, which were determined with error propagation (using the uncertainties python package) using the 95% confidence intervals for $\eta_{xz}$ and $\eta_{yz}$ that were estimated as detailed in SI Sec. 17 and the Materials and Methods of the main text. Under the null hypothesis that there is no causal interaction from $X$ to $Z$, the expected value of $\eta_{xz}/\eta_{yz} = 1$. If the measured covariability ratio is above (below) the expected value of 1, then there is a 2.5% probability that the expected value of 1 falls below (above) the lower (upper) error bar. As a result, under a significance level of 2.5%, we say there is a causal interaction from $X$ to $Z$ if the value of $\eta_{xz}/\eta_{yz} = 1$ does not fall within the measured 95% confidence intervals. Using this rule, our method detected two causal interactions (synthetic circuits #1 and #2) out of four. **B)** Under the rule for detecting causality outlined in **A**, we detect a causal interaction in synthetic circuit #12 when we set the RFP concentration as the $Z$ component of interest. Plotted are the measured covariability ratios $\eta_{xz}/\eta_{yz}$ for the synthetic circuits shown in Fig. 3E and Tables. S3 and S4. **C)** Similarly to **B**, we detect a causal interaction in synthetic circuit # 12 when we set the growth rate as the $Z$ component of interest. All other negative control circuits satisfy the null hypothesis test.

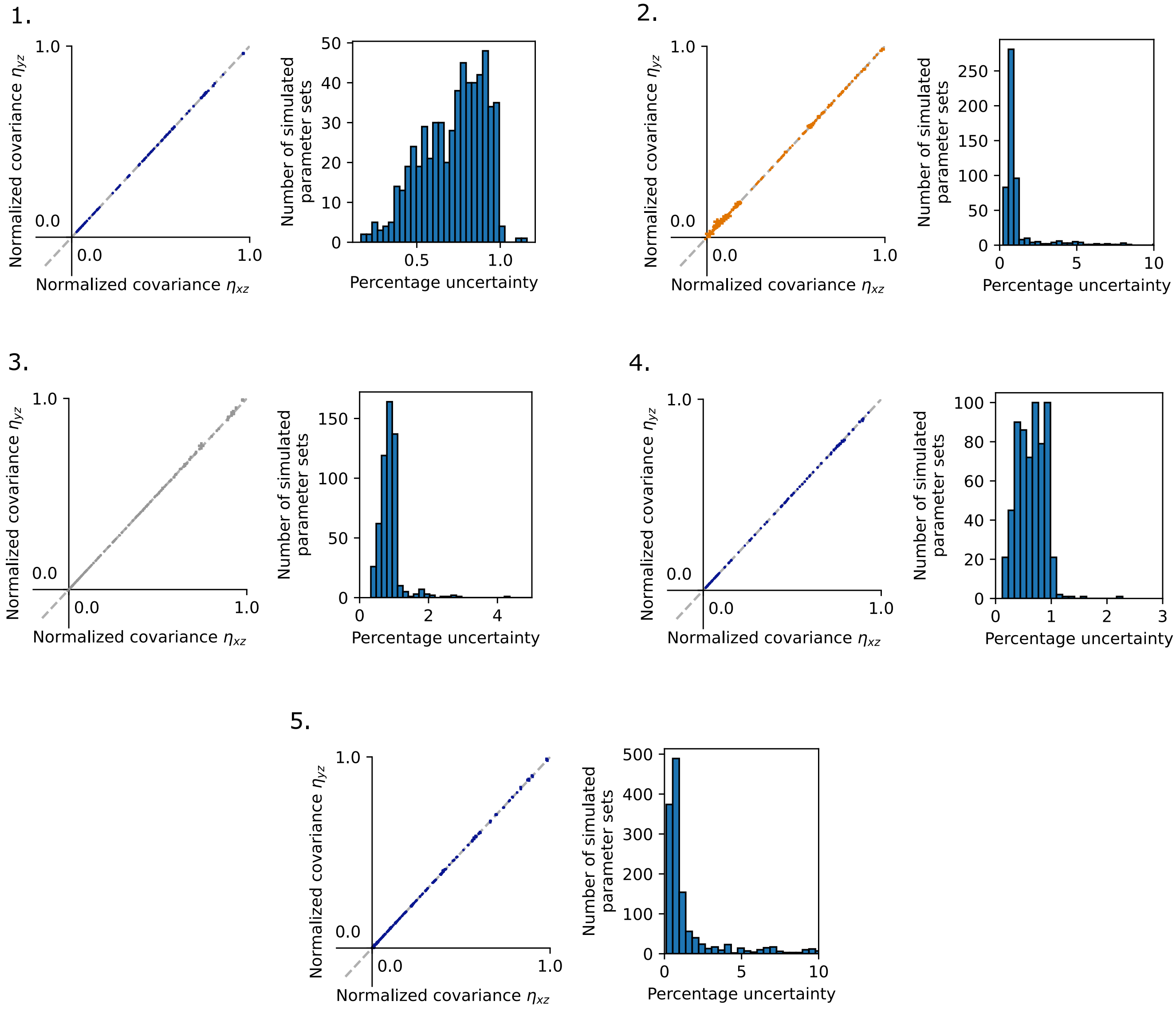


Figure S3. **Percentage uncertainty distributions for individual simulated processes — part 1.** Plotted are the computed normalized covariances from the simulated concentrations of growing and dividing cells in processes 1 to 5 (as listed in Tab. S1). Each number in the top left corners correspond to the process index listed in Tab. S1. For each process, a number of random parameters were chosen and simulated as described in the Materials and Methods section of the main text. On the left of each subplot are the normalized covariances for each of the parameter sets simulated in the respective process. Errorbars are present but are almost always too small to be visible. On the right of each subplot is the distribution of percentage uncertainty, defined as $\sigma_{xz}/|\eta_{xz}| \times 100\%$ and $\sigma_{yz}/|\eta_{yz}| \times 100\%$, where $\sigma_{xz}$ is the estimated 95% confidence interval for $\eta_{xz}$. Most of the percentage errors fall below 5%. However, occasionally a parameter set is chosen that leads to either rare birth and death events in the stochastic process which leads to low sampling, or the particular parameters lead to near zero normalized covariances which can lead to large percentage errors. Colors of dots and errorbars in the left plots correspond to the same colors plotted in the combined figure of Fig. 2.

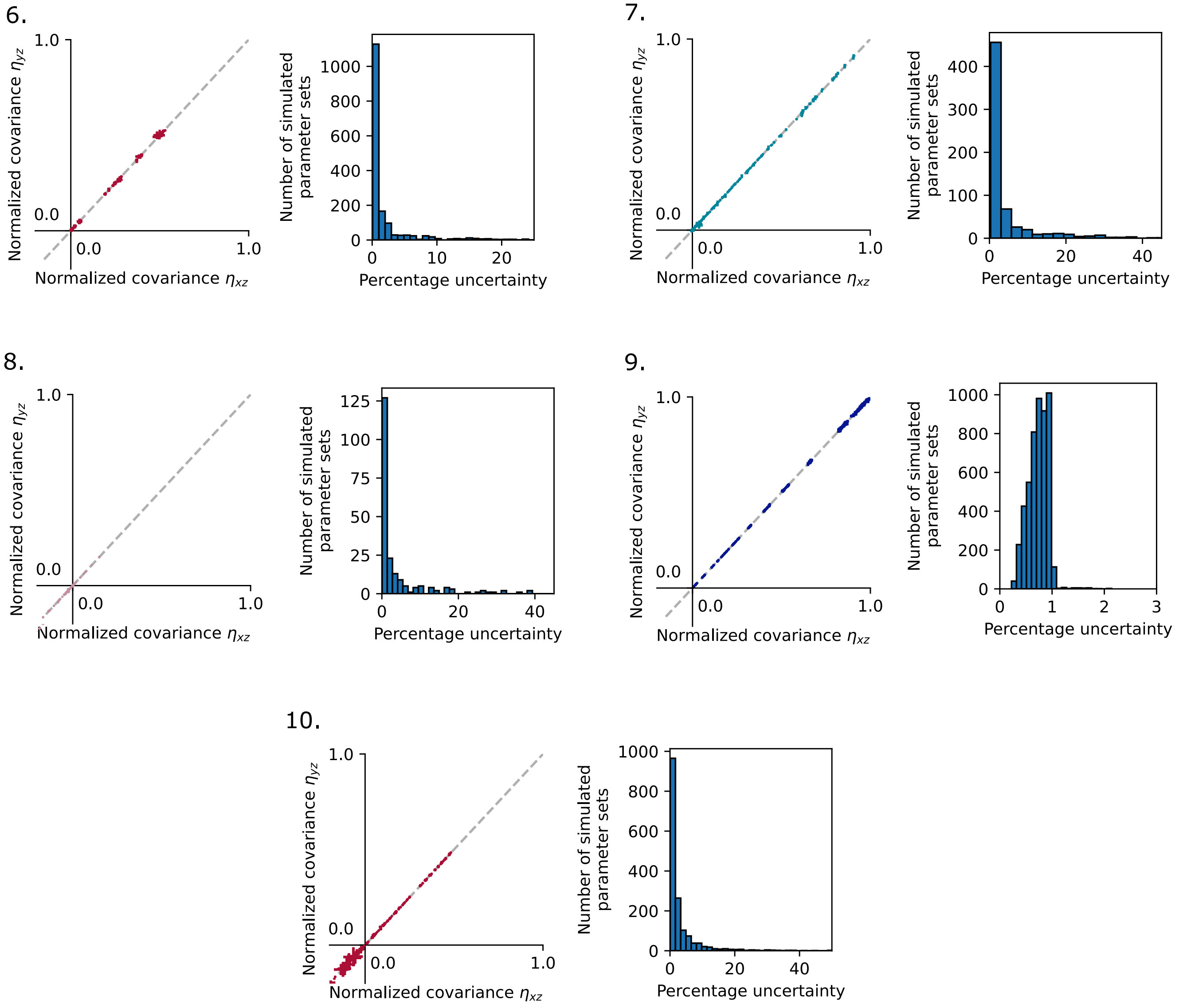


Figure S4. **Percentage uncertainty distributions for individual simulated processes — part 2.** Same as Fig. S3 but for processes 6 to 10 (as listed in Tab. S1).

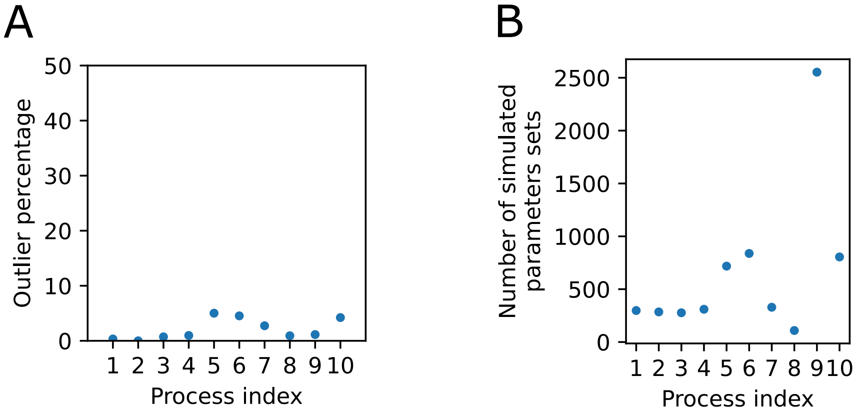


Figure S5. **Percentage of simulations that deviate from the invariant of Eq.** (2)**. A)** Plotted is the percentage of simulated parameter sets for each simulated process (as listed in Tab. S1) that did not produce normalized covariances that satisfied Eq. (2). Specifically, the 95% confidence interval for the ratio $\eta_{xz}/\eta_{yz}$ in these "outlier" simulations did not capture the predicted value of 1. It is expected that not all simulations will agree with Eq. (2) due to sampling errors. The largest outlier percentage is for process 5, which has 5.01% of simulations that did not satisfy Eq. (2). As described in the Materials and Methods section of the main text, all outlier systems were simulated again with additional sampling which resulted in agreement with Eq. (2). **B)** Plotted is the number of parameter sets simulated for each process. Some processes took less time to simulate and as a result resulted in a larger number of parameter sets. All processes were simulated for at least 100 parameter sets.

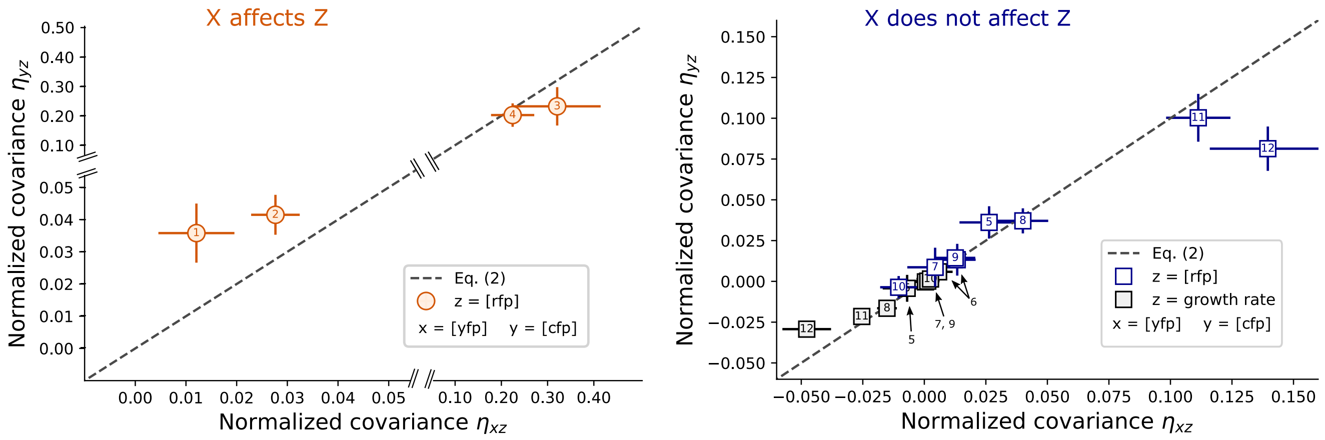


Figure S6. **Estimating uncertainty with a second method does not change the results of Fig. 3D,E.** Shown are the measured normalized covariances for the synthetic positive control circuits (left) and negative control circuits (right), where error bars are computed by splitting the ensemble of single-cell traces into 7-10 disjoint sets. Final normalized covariances are taken as the averages of the sets, with 95% confidence intervals given by two times the standard error of the mean (plotted errorbars). Data corrections such as non-even illumination correction and autofluorescence removal was done separately on each set in order to estimate the error of these corrections. Autofluorescence and imaging media fluorescence were corrected by analyzing cells that lost the synthetic plasmid due to random partitioning at cell division, as describe in Materials and Methods. If 10 or more cells from an experiment lost the plasmid, then the single-cell traces were randomly split into 10 disjoint sets, both with and without the plasmid. Otherwise, the number of disjoint sets was set as the number of cells that lost the plasmid (a minimum of 7), so that each set would contain at least one cell that lost the plasmid to estimate the autofluorescence. This is described in the Materials and methods section of the main text, as well as in Fig. S16 and SI Sec. 17.

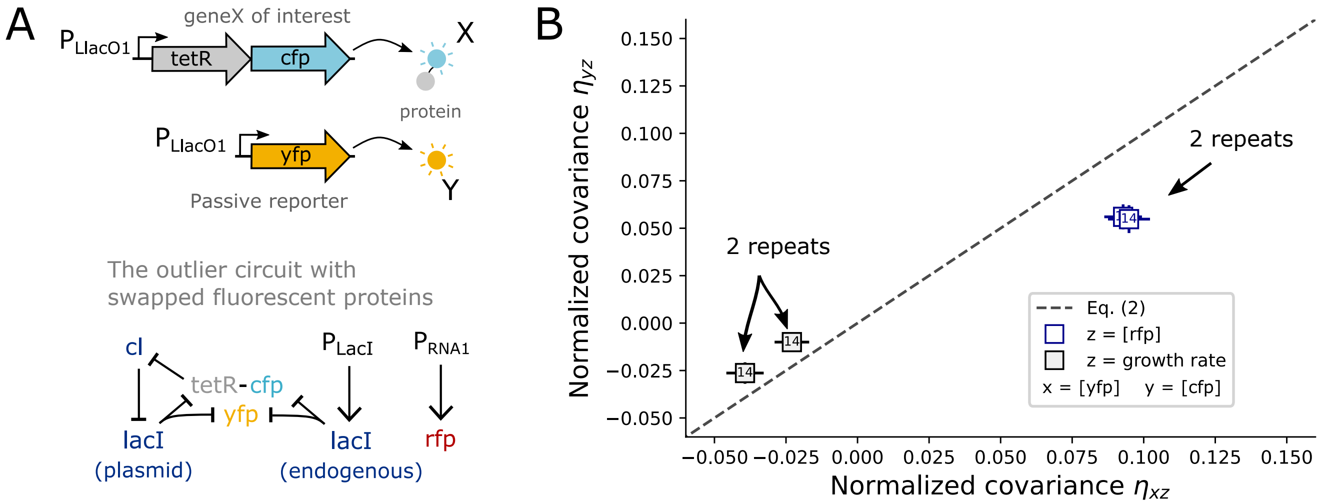


Figure S7. **The outlier circuit does not satisfy Eq. (2) after switching the fluorescent proteins. A)** To rule out the possibility that the outlier in Fig. 3E was caused by a systematic error from an asymmetry between measurements of the CFP and YFP fluorescence signals, we rebuilt the outlier circuit, but with *cfp* fused to *tetR* and *yfp* set as the transcriptional passive reporter. If the outlier was caused by such a systematic error, then the normalized covariance measurements would switch to the other side of the dashed line. **B)** The normalized covariances still do not satisfy Eq. (2), and they do not switch to the other side of the dashed line. Two repeats of the experiment are shown. Note that the numerical values for the normalized covariances are different than the outlier circuit in Fig. 3E. This could be indicative that the CFP protein used changes the TetR function when used as a fusion.

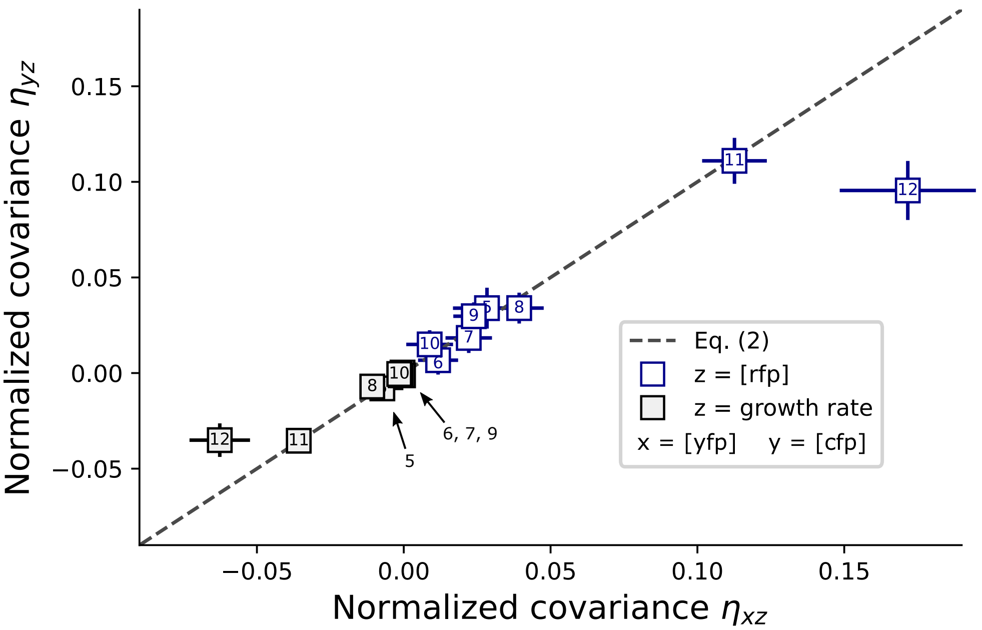


Figure S8. **Using a different cell segmentation pipeline does not alter the main result of Fig. 3E.** For the results shown in Fig. 3 we used the deep-learning cell segmentation pipeline DeLTA [1], see Materials and Methods. We used the DeLTA pipeline to segment cells with the bright field channel because the three fluorescence channels (CFP, YFP, and RFP) were not always bright enough in the positive control strains to use as a segmentation marker. However, in all the negative control strains, RFP is under the control of the strong pRNA1 promoter which has been used as a segmentation marker previously [2, 3]. We thus applied the same segmentation pipeline used in [2, 3] on the RFP channel of the negative control strains. The numerical values of the normalized covariances change slightly, but their relation to Eq. (2) remains intact. This second segmentation pipeline determines the edges of the cells using a thresholding algorithm, and it estimates background fluorescence by taking the median of the median of each image, see [3] for details. The autofluorescence of strains 6, 7, 8, 9, and 10 were not removed because these strains express RFP from the plasmid, so cells that lost the plasmid (which we use to estimate autofluorescence, see Materials and Methods) cannot be tracked with this pipeline.

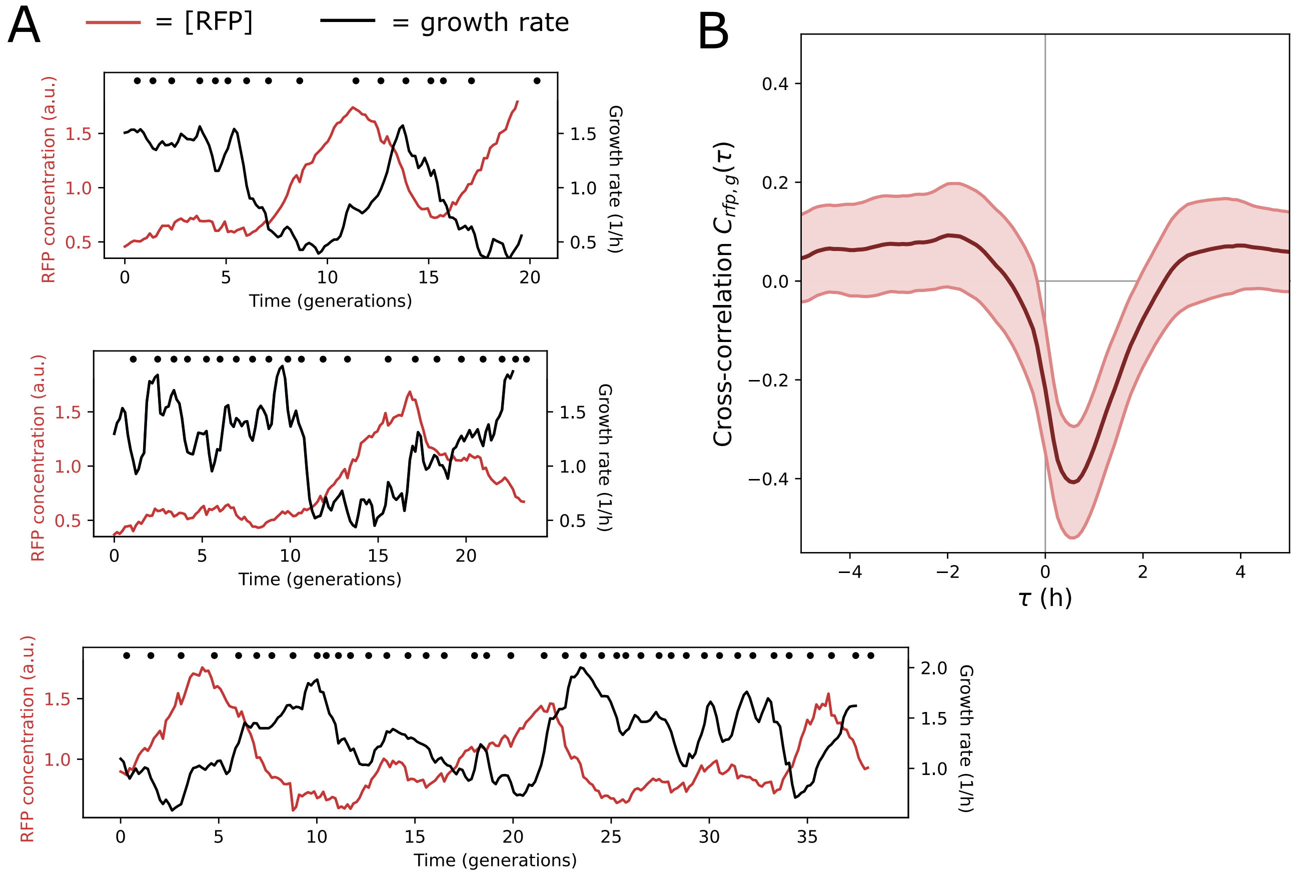


Figure S9. **In the outlier circuit, large RFP fluctuations are often negatively correlated with growth rate fluctuations.** We observed many instances in which RFP concentration was negatively correlated with fluctuations in the cellular growth rate. Shown are three sample single-cell traces. The top two are from the outlier circuit, whereas the bottom trace is from the outlier circuit with swapped fluorescent proteins (see Fig. S7A). Dots correspond to times at which a cell division has been detected. **B)** To quantify this trend over all lineages we used the cross-correlation function $C_{\mathrm{rfp,g}}(\tau)$ between the RFP concentration and growth rate (g) in the outlier circuit. We observe a negative correlation time shifted by approximately 1 hour. $C_{\mathrm{rfp,g}}(\tau)$ quantifies how the RFP concentration is correlated with growth rate when the growth rate signal is shifted by time $\tau$ relative to the RFP signal. $C_{\mathrm{rfp,g}}(\tau)$ was computed using the python scipy function correlate on each single-cell trajectories while normalizing by $\sqrt{C_{\mathrm{rfp,rfp}}(0)C_{\mathrm{g,g}}(0)}$. The dark line corresponds to the average at time $\tau$ over all trajectories, with the width of the shaded region is given by the standard deviation about the mean.

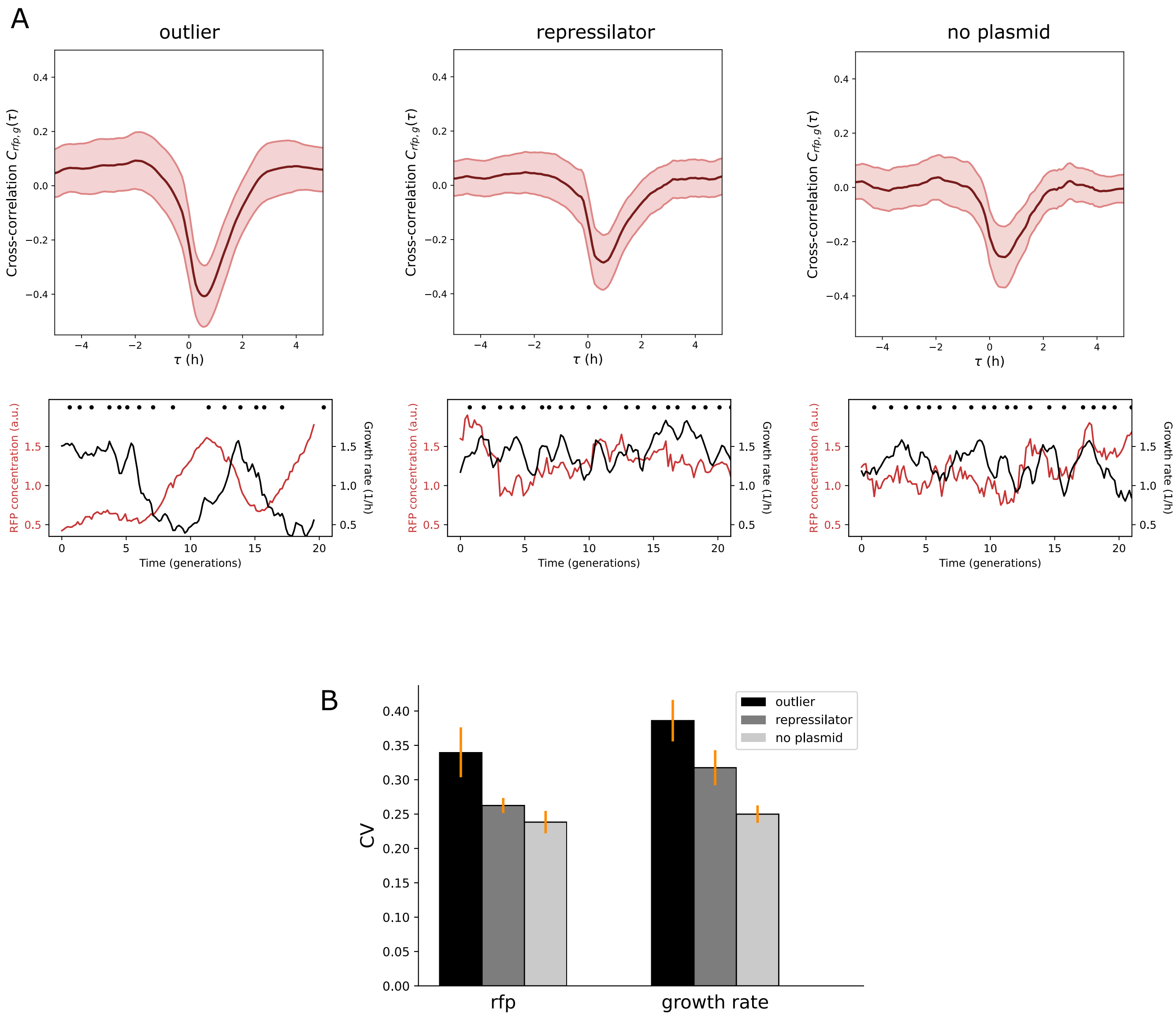


Figure S10. **The outlier circuit shows more variability in RFP concentrations and growth rate measurements as compared to other strains that express RFP from the chromosome with the pRNA1 promoter. A)** The "regular" repressilator circuit without endogenous LacI also exhibits negative correlation between RFP and growth rate, albeit with smaller magnitude. Shown are the cross correlations (defined in Fig. S9B) for the outlier circuit (left), the strain with the regular repressilator (outlier circuit with *lacI* deletion, middle), and the background strain without a plasmid but with RFP still expressed (right). In all cases, RFP is expressed from the chromosome under the control of the same pRNA1 promoter. The outlier circuit exhibits slightly more negative correlation. Below each cross correlation plot is a sample time-trace for a cell with the respective circuit. Dots correspond to times at which a cell division has been detected. **B)** The "regular" repressilator circuit without endogenous LacI does not exhibit the same degree of growth rate fluctuations and does not exhibit as pronounced RFP pulses. The size of the fluctuations of the growth rate and RFP concentration was measured as the coefficient of variability (CV), defined as $\mathrm{CV}_z = \sqrt{\mathrm{Var}(z)}/\langle z\rangle$. The CVs were measured for the outlier circuit, the strain with the regular repressilator (outlier circuit with *lacI* deletion), and the background strain without a plasmid but with RFP still expressed. The outlier circuit exhibits a larger CV in both cases, suggesting that the outlier circuit causes the increased growth rate fluctuations. Orange error bars correspond to 95% confidence intervals estimated using the same bootstrapping approach used for the normalized covariances described in the Materials and Methods of the main text.

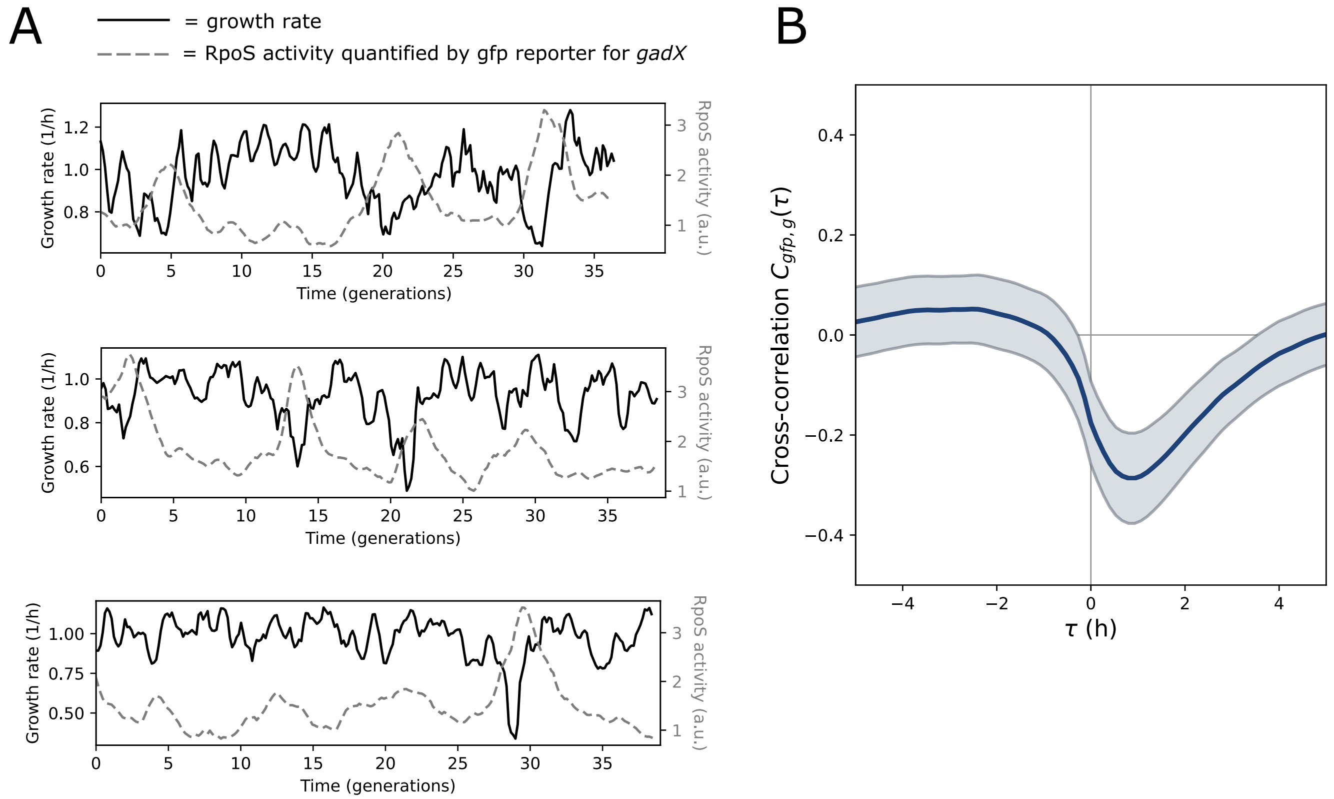


Figure S11. **RpoS activity often displayed pulses that are negatively correlated with growth rate.** We observed many instances in which RpoS activity, quantified by a transcriptional GFP reporter under the control of the *gadX* promoter, was negatively correlated with fluctuations in the cellular growth rate. Shown are three sample single-cell traces. **B)** To quantify this trend over all lineages we used the cross-correlation function $C_{\mathrm{gfp,g}}(\tau)$ between the GFP concentration and growth rate (g). We observe a negative correlation time shifted by approximately 1 hour. $C_{\mathrm{gfp,g}}(\tau)$ quantifies how the GFP concentration is correlated with growth rate when the growth rate signal is shifted by time $\tau$ relative to the GFP signal. It was computed as described in the Fig. S9 caption.

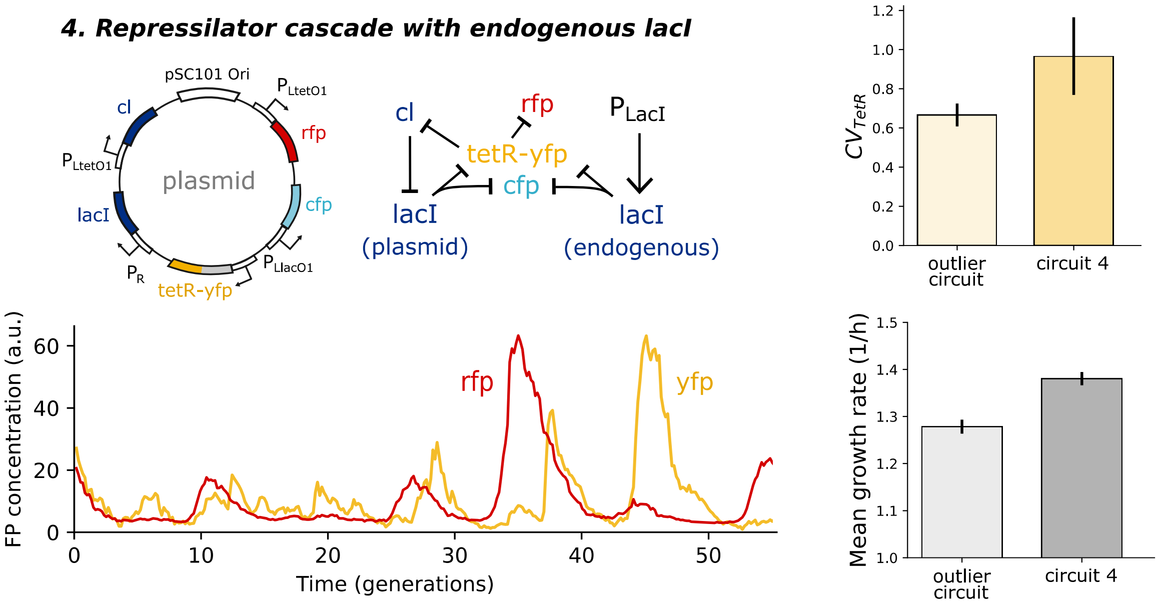


Figure S12. **Removing the pRNA1 controlled *rfp* gene changes the TetR dynamics of the outlier circuit and increases the growth rate.** One of the positive control circuits (4) contains the same synthetic circuit as the outlier circuit, but without the pRNA1 controlled *rfp* on the chromosome, and instead a pLtetO1 controlled *rfp* gene on the plasmid. We observed that the TetR dynamics, quantified by YFP, were different than in the outlier circuit as they displayed random large pulses. The RFP levels in turn also displayed random large pulses when the TetR levels were reduced. To quantify these observations over all lineages, we computed the CV for the TetR levels for both strains (top right). The strain 4 CV is larger. We also computed the average growth rate for both strains and find that the strain 4 growth rate is approximately 9% larger. These results suggest that the pRNA1 controlled *rfp* gene affected the rest of the circuit elements, along with the cellular growth rate, which we hypothesize is from burden caused by resource competition. Error bars correspond to 95% confidence intervals using the boostrapping approach used for the normalized covariances as discussed in the Materials and Methods section.

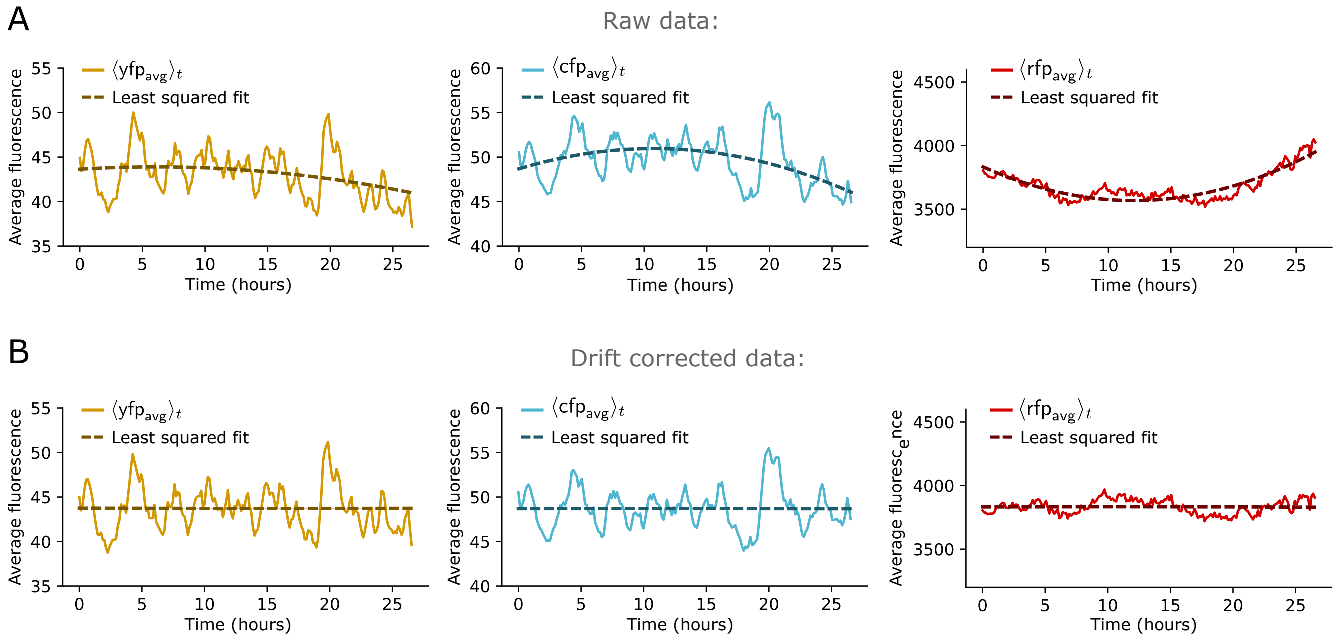


Figure S13. **Correcting for temporal drift of the focal points. A)** Individual average fluorescence measurements (taken over cell area) are binned according to their time-frame. The average of the measurements at each time-frame is plotted. A second degree polynomial is fitted to the resulting time-traces using a least squared fit. Shown is data from the strain 6 in Table. S3. The YFP and CFP time-traces fluctuate more than the RFP time-trace due to the large pulses that they display from the Repressilator. **B)** Temporal drift is corrected by multiplying all of the individual average fluorescence measurements with the reciprocal of the fits. Plotted is the average time-traces taken from the corrected data, along with new fits that display constant trends.

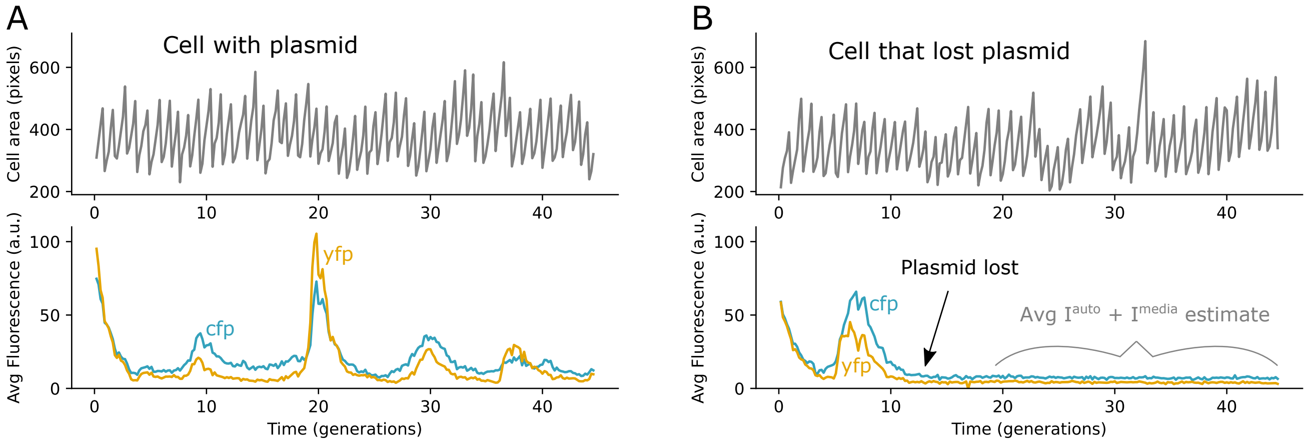


Figure S14. **Estimating autofluorescence and media fluorescence with cells that lost plasmid A)** A single-cell time trace from a mother machine experiment in which the plasmid is kept for the entire duration of the experiment. On the top is the cell area over time displaying cellular growth and division, and on the bottom are the average YFP and CFP signals taken over the cell area. Note that autofluorescence and media fluorescence have not been corrected for here. **B)** A single-cell time trace in which the plasmid containing the synthetic circuit is lost. The signals decay to constant values just above zero. The cell area continues to grow and divide, indicating that the decay is not the result of segmentation error or loss of cell.

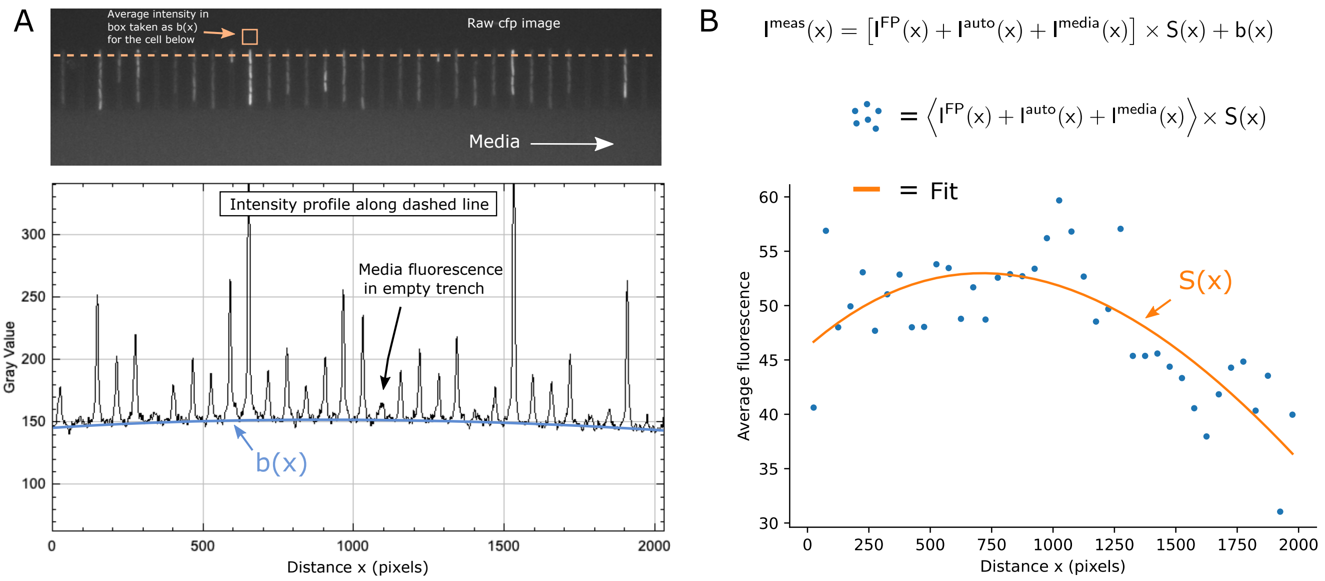


Figure S15. **Correcting for background and uneven illumination. A)** Plotted is the fluorescence intensity profile across the dashed orange line shown in a single image of the CFP channel. The background $b(x)$ is traced by the light blue solid line in the bottom plot, to illustrate the slight $x$ dependence. To estimate and remove the background, in each image the average fluorescence across a 50x50 pixel box located 15 pixels above each mother cell was removed from the fluorescence measurement of each respective cell in that image. Note also that an empty trench produces a significant amount of fluorescence when compared to cell containing trenches. This indicates that fluorescence originating from the media itself needs to be considered. This will be discussed in the next section. The fluorescence profile was created using the plot profile function in Fiji ImageJ. **B)** With the background $b(x)$ removed, it remains to correct for $S(x)$. We bin all of the fluorescence measurement of all cells and over all time according to pixel position $x$ in bins of size 50 pixels and take the average of each bin (blue dots). A single frame has a width of 2048 pixels. This gives an estimate of $\langle I^{FP} + I^{auto} + I^{media} \rangle \times S(x)$, with deviations resulting from finite sampling. The binned averages are fit using the least squared method with a third degree polynomial (solid orange line). The uneven illumination gain function $S(x)$ is corrected by multiplying all the fluorescence measurements with the reciprocal of the fitted polynomial. Data shown is from the strain 6 system from Table. S3.

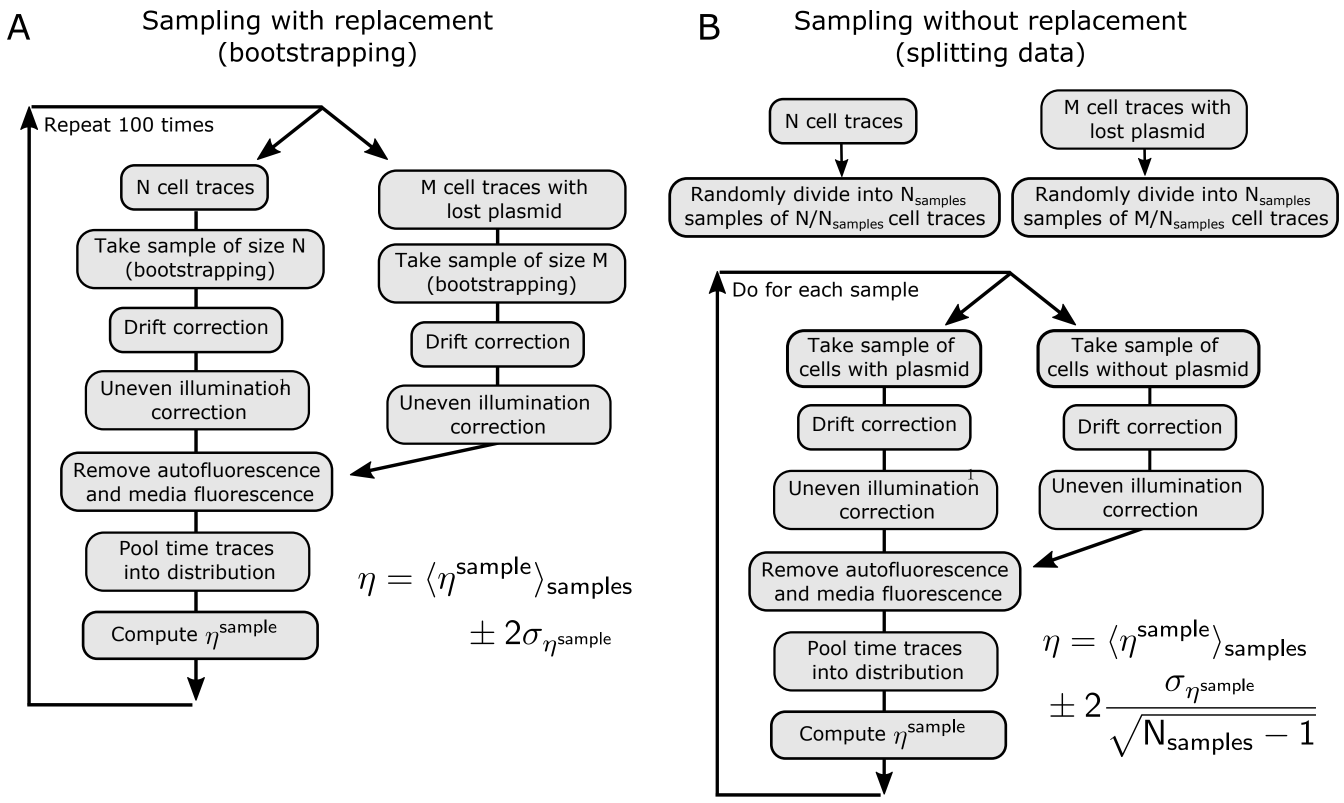


Figure S16. **Estimating confidence intervals for normalized covariance measurements using two methods. A)** Imaging error corrections and the normalized covariance are computed 100 times for 100 bootstrap samples of the experimental data. The final normalized covariance is given as the average over the samples, with confidence intervals given by 2 times the standard deviation. **B)** The data is divided into $N_{samples} = \min(M, 10)$ disjoint sample sets, where $M$ is the number of cells that lost the plasmid. Most experiments produced $M > 10$, but a few produced $M < 10$ (with a minimum of 7). Imaging error corrections and the normalized covariance are computed for each sample. The final normalized covariance is given by the average over the $N_{samples}$ samples, with confidence intervals given by 2 times the standard error of the mean.

**Movie. S1.** **Bright field images of a single imaging position of a mother machine experiment with colored cell boundaries produced from the DelTA segmentation pipeline. Here segmentation was done on bright field images.**

**Movie. S2.** **Fluorescent images of a single imaging position of a mother machine experiment with colored cell boundaries produced from the DelTA segmentation pipeline. Here segmentation was done on the RFP channel, which produced similar results to those in Fig. 3.**

### 2. Supplementary tables

#### General rates for four component system

$$x \xrightarrow{r_x^+} x+1 \quad y \xrightarrow{r_y^+} y+1 \quad z \xrightarrow{r_z^+} z+1 \quad w \xrightarrow{r_w^+} w+1$$
$$x \xrightarrow{r_x^-} x-1 \quad y \xrightarrow{r_y^-} y-1 \quad z \xrightarrow{r_z^-} z-1 \quad w \xrightarrow{r_w^-} w-1$$

| **Process index** | $r_x^+$ | $r_x^-$ | $r_y^+$ | $r_y^-$ | $r_z^+$ | $r_z^-$ | $r_w^+$ | $r_w^-$ |
|---|---|---|---|---|---|---|---|---|
| 1 | $\frac{\lambda z}{K+x}$ | $\beta x$ | $\frac{\lambda z}{K+x}$ | $\beta y$ | $\lambda_z$ | $\beta_z z$ | 0 | 0 |
| 2 | $\frac{\lambda(x+\delta)}{\left(K+x+\left(\frac{y}{z+\delta}\right)^2\right)}$ | $\beta x$ | $\frac{\lambda(x+\delta)}{\left(K+x+\left(\frac{y}{z+\delta}\right)^2\right)}$ | $\beta y$ | $\lambda_z$ | $\beta_z z$ | 0 | 0 |
| 3 | $\frac{\lambda w}{K+x^2}$ | $\beta x$ | $\frac{\lambda w}{K+x^2}$ | $\beta y$ | $\lambda_z w$ | $\beta_z z$ | $\lambda_w$ | $\beta_w w$ |
| 4 | $\frac{\lambda\left(\frac{z}{V(t)}\right)}{1+\left(\frac{x}{V(t)}\right)^2}$ | $\beta x$ | $\frac{\alpha\lambda\left(\frac{z}{V(t)}\right)}{1+\left(\frac{x}{V(t)}\right)^2}$ | $\beta y$ | $\lambda_z$ | $\beta_z z$ | 0 | 0 |
| 5 | $\frac{\alpha\lambda V(t)}{1+\left(\frac{x/V(t)}{\delta+z/V(t)}\right)^3}$ | $\beta x$ | $\frac{\lambda V(t)}{1+\left(\frac{x/V(t)}{\delta+z/V(t)}\right)^3}$ | $\beta y$ | $\lambda_z V(t)$ | $\beta_z z$ | 0 | 0 |
| 6 | $A\left(\sin(\omega t)+1\right)V(t)$ | $\beta x$ | $B\left(\sin(\omega t)+1\right)V(t)$ | $\beta y$ | $C\left(\sin(\omega t)+1\right)V(t)$ | $\beta_z z$ | 0 | 0 |
| 7 | $\frac{\alpha\left(\frac{y}{V(t)}+\delta\right)}{\frac{y}{V(t)}+\delta+\left(\frac{x/V(t)}{\delta+w/V(t)}\right)^2}$ | $\beta x$ | $\frac{\left(\frac{y}{V(t)}+\delta\right)}{\frac{y}{V(t)}+\delta+\left(\frac{x/V(t)}{\delta+w/V(t)}\right)^2}$ | $\beta y$ | $\lambda_z w/V(t)$ | $\beta_z z^2$ | $\lambda_w V(t)$ | $\beta_w w$ |
| 8 | $\frac{\lambda}{1+\left(\frac{x/V(t)}{\delta+w}\right)^3}$ | $\beta x$ | $\frac{\lambda}{1+\left(\frac{x/V(t)}{\delta+w}\right)^3}$ | $\beta y$ | $\frac{\lambda_z}{1+\left(\frac{w/V(t)}{k}\right)^3}$ | $\beta_z z$ | $\frac{\lambda_z}{1+\left(\frac{z/V(t)}{k}\right)^3}$ | $\beta_w w$ |
| 9 | $\frac{\alpha\lambda z}{K+x}$ | $xz$ | $\frac{\lambda z}{K+x}$ | $yz$ | $\lambda_z$ | $\beta_z z$ | 0 | 0 |
| 10 | $A\left(\sin(\omega t)+1\right)V(t)$ | $\beta xz$ | $B\left(\sin(\omega t)+1\right)V(t)$ | $\beta yz$ | $C\left(\sin(\omega t)+1\right)V(t)$ | $\beta_z z$ | 0 | 0 |

Table S1. **Simulated birth-death processes in growing and dividing cells.** Additional details in Sec. 11. Top box shows a general four component birth-death process. Bellow the box shows the respective rates for the 10 processes simulated. For each process, three different single-cell volume trajectories $V(t)$ were simulated according to the Materials and Methods of the main text, and the rate parameters shown above were varied randomly multiple times. In processes 1-3, 8, and 9, $\lambda$ was picked randomly from the set $\{10, 100, 1000\}$, whereas in processes 4, 5, and 7 it was picked randomly from the set $\{1, 10, 100\}$. The parameter $\delta$ is set to 0.01 throughout to avoid divisions by 0. In processes 4, 5, 7, and 9, $\alpha$ is respectively set to $(2, 0.1, 1)$, $(2, 0.5, 0.1)$, $(2, 1, 0.5)$, $(1, 2, 0.5)$ for all simulations with respective volume dynamics $(1, 2, 3)$. For example, in system 4 with volume dynamics 2, $\alpha = 0.1$. All other parameters, namely $\beta, \beta_z, \beta_w, \lambda_z, \lambda_w, K, A, B, C, \omega$, were picked randomly from the set $\{0.1, 1, 10\}$. In all simulations the average division time is set to 1. Simulations were performed using the Gillespie algorithm, with an adjustments to account for time-dependent rates, as described in 13. Simulation data and code are available on Github.

***1. Open-loop cascade I***

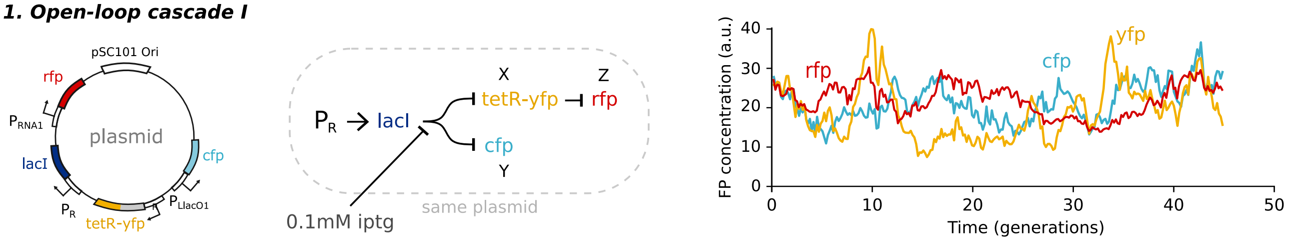


***2. Open-loop cascade II***

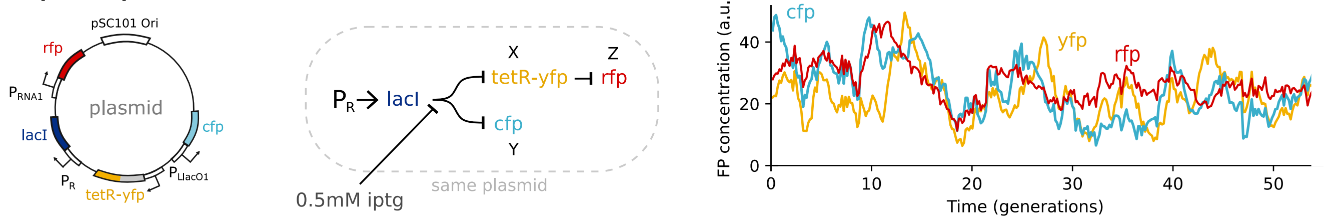


***3. Repressilator cascade***

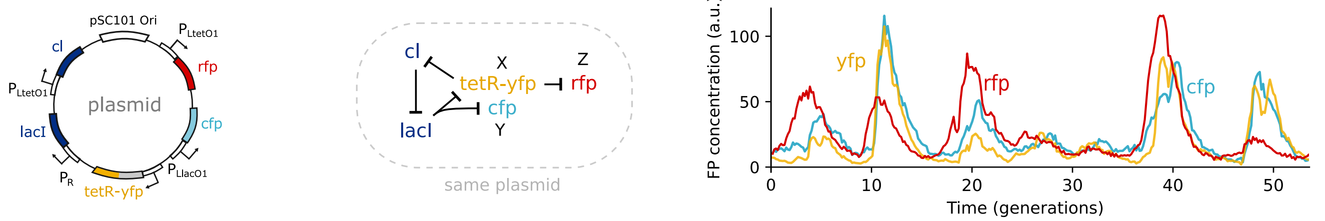


***4. Repressilator cascade with endogenous lacI***

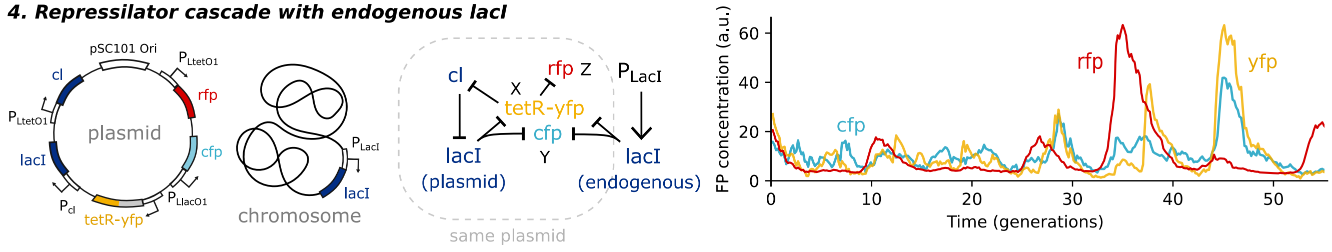


Table S2. **Positive control synthetic circuits with sample single-cell time traces.** Plotted fluorescence units are not consistent from sample to sample.

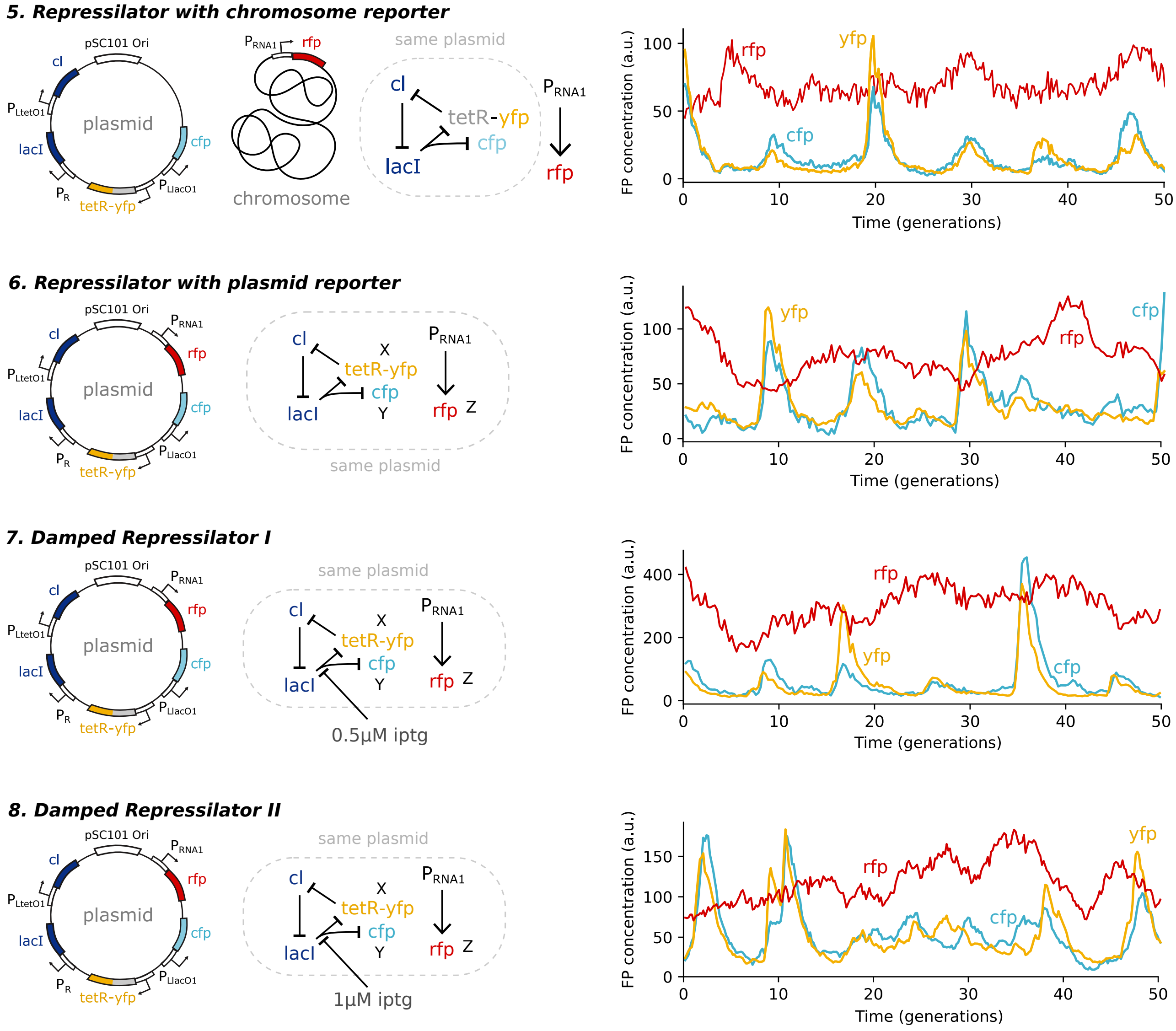


Table S3. **Negative control synthetic circuits with sample single-cell time traces – part 1.** Plotted fluorescence units are not consistent from sample to sample.

***9. Damped Repressilator III***

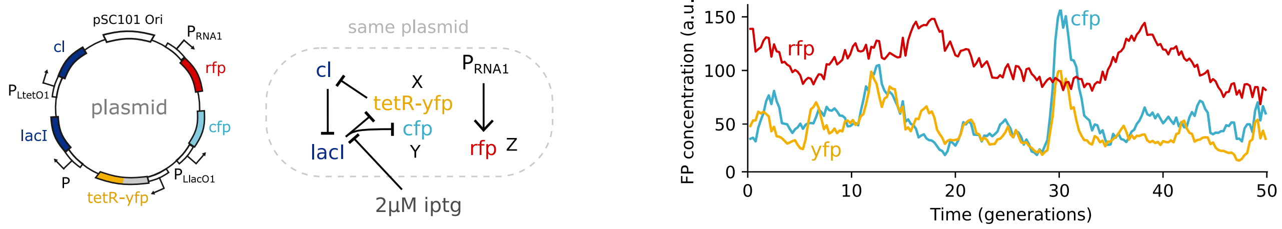


***10. The Repressilator with titration sponge***

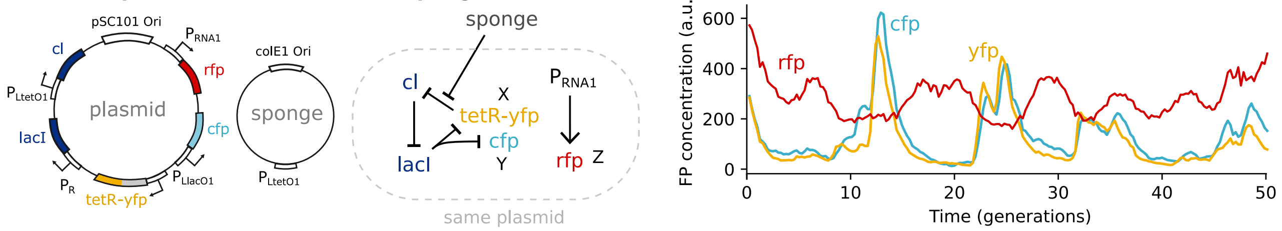


***11. The Repressilator mutant***

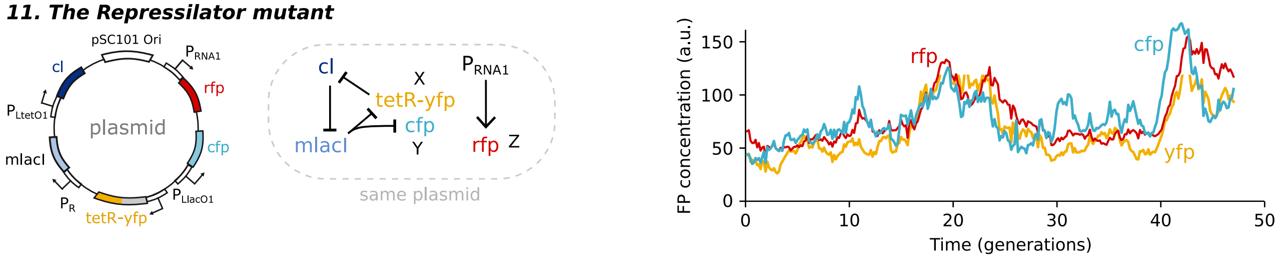


***12. Repressilator with endogenous lacI (outlier circuit)***

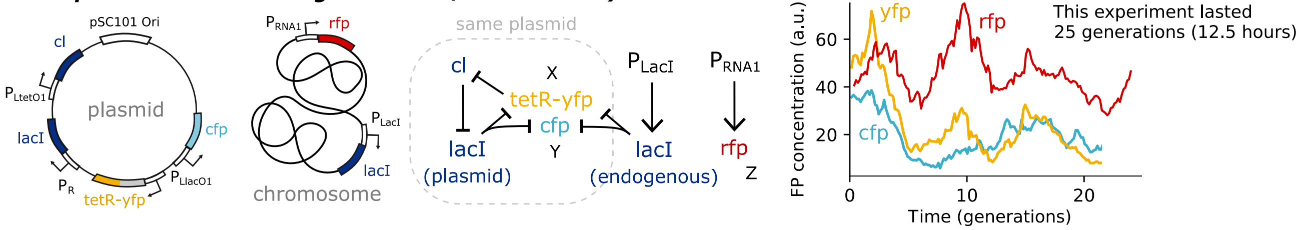


Table S4. **Negative control synthetic circuits with sample single-cell time traces – part 2.** Plotted fluorescence units are not consistent from sample to sample.

***13. Repressilator with endogenous lacI and RpoS deletion***

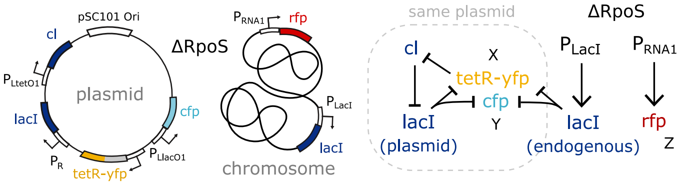


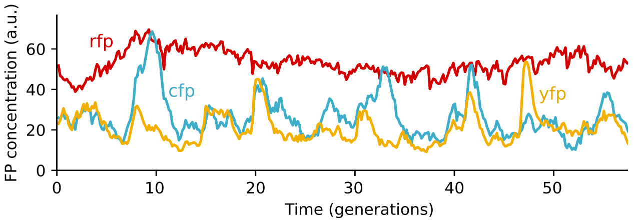


***14. Repressilator with endogenous lacI - swapped FPs***

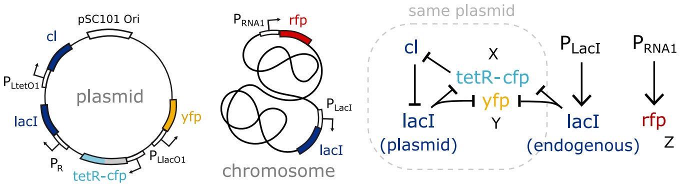


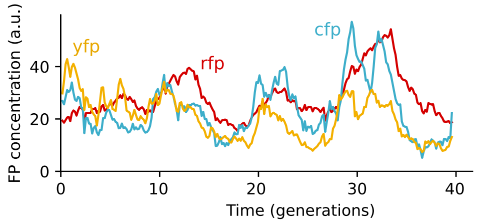


Table S5. **Other synthetic circuits.**

| Strain | Details |
|---|---|
| EJS1 | Synthetic circuit 5. Background strain is MG1655 *E. coli* attB::*lacYA177C*, Δ*araCBAD*, Δ*lacIZYA*, Δ*araE*, Δ*araFGH*, Δ*rhaSRT*, Δ*rhaBADM*, *attn7::mCherry-mKate2-hybrid*. Contains pEJS1. |
| EJS2 | Synthetic circuit 12. Background strain is MG1655 *E. coli* with *glmS::PRNA1-mCherry-mKate2-hybrid* and motility deletion Δ*motA*. Contains pEJS1. |
| EJS3 | Synthetic circuits 1 and 2. Background strain is MG1655 *E. coli* with attB::*lacYA177C*, Δ*araCBAD*, Δ*lacIZYA*, Δ*araE*, Δ*araFGH*, Δ*rhaSRT*, and Δ*rhaBADM*. Contains plasmid pEJS3. |
| EJS4 | Synthetic circuit 3. Background strain is MG1655 *E. coli* with attB::*lacYA177C*, Δ*araCBAD*, Δ*lacIZYA*, Δ*araE*, Δ*araFGH*, Δ*rhaSRT*, and Δ*rhaBADM*. Contains plasmid pEJS4. |
| EJS6 | Synthetic circuit 4. Background strain is MG1655 *E. coli* with motility deletion Δ*motA*. Contains plasmid pEJS4. |
| EJS7 | Synthetic circuit 11. Background strain is MG1655 *E. coli* with attB::*lacYA177C*, Δ*araCBAD*, Δ*lacIZYA*, Δ*araE*, Δ*araFGH*, Δ*rhaSRT*, and Δ*rhaBADM*. Contains plasmid pEJS5m. |
| EJS9 | Synthetic circuit 14. Background strain is MG1655 *E. coli* with *glmS::PRNA1-mCherry-mKate2-hybrid* and motility deletion Δ*motA*. Contains plasmid pEJS10. |
| PA12 | Strain with no plasmid in Fig. S10. Background strain is MG1655 *E. coli* attB::*lacYA177C*, Δ*araCBAD*, Δ*lacIZYA*, Δ*araE*, Δ*araFGH*, Δ*rhaSRT*, Δ*rhaBADM*, *attn7::mCherry-mKate2-hybrid*. |
| PA52 | Synthetic circuits 6, 7, 8, and 9. Background strain is MG1655 *E. coli* with attB::*lacYA177C*, Δ*araCBAD*, Δ*lacIZYA*, Δ*araE*, Δ*araFGH*, Δ*rhaSRT*, and Δ*rhaBADM*. Contains plasmid pEJS5. |
| PA53 | Synthetic circuit 10. Background strain is MG1655 *E. coli* with attB::*lacYA177C*, Δ*araCBAD*, Δ*lacIZYA*, Δ*araE*, Δ*araFGH*, Δ*rhaSRT*, and Δ*rhaBADM*. Contains plasmids pEJS5 and pLPT41. |
| PA64 | Used in Fig. 4D. Background strain is MG1655 *E. coli* with *glmS::PRNA1-mCherry-mKate2-hybrid* and motility deletion Δ*motA*. Contains pSC101 plasmid with Kanamycin resistance and *gfpmut2* reporter gene with *gadX* promoter taken from [4]. |
| PA66 | Synthetic circuit 13. Background strain is MG1655 *E. coli* with *glmS::PRNA1-mCherry-mKate2-hybrid*, motility deletion Δ*motA*, and *DRpos::FRT Kan frt*. Contains plasmid pEJS1. |

| Plasmid | Details |
|---|---|
| pEJS1 | Used in circuits 5, 12, and 13. Repressilator with no degradation tags, with *tetR* replaced with *tetR-mVenusNB fusion*, along with an integrated *SCFP3A* reporter. pSC101 origin. Ampicillin resistance. |
| pEJS3 | Used in circuit 1 and 2. Repressilator with no degradation tags with *cI* replaced with *mCherry-mKate2-hybrid*, *tetR* replaced with *tetR-mVenusNB fusion*, and an integrated *SCFP3A* reporter. pSC101 origin. Ampicillin resistance. |
| pEJS4 | Used in circuit 3 and 4. Repressilator with no degradation tags, with *tetR* replaced with *tetR-mVenusNB fusion*, and integrated *SCFP3A* and *mCherry-mKate2-hybrid* reporters. pSC101 origin. Ampicillin resistance. |
| pEJS5 | Used in circuits 6, 7, 8, and 9. Repressilator with no degradation tags, with *tetR* replaced with *tetR-mVenusNB fusion*, and integrated *SCFP3A* and *mCherry-mKate2-hybrid* reporters. pSC101 origin. Ampicillin resistance. |
| pEJS5m | Used in circuit 11. Same as pEJS5, with a single missense point mutation on *lacI* that results in no oscillations forming (see 11 in Tab. S4). LacI function must be affected by the mutation. pSC101 origin. Ampicillin resistance. |
| pEJS10 | Used in circuit 14. Same as pEJS1 but *SCFP3A* forms the fusion with *tetR* while *mVenusNB* is integrated as a transcriptional reporter. pSC101 origin. Ampicillin resistance. |
| pLPT41 | Used in circuit 10. ColE1 origin plasmid with $P_{LtetO1}$ promoter, taken from [2]. Acts as a "titration sponge" for small numbers of TetR levels using repressor-binding sites, see [2]. Kanamycin resistance. |

Table S6. **Strains and plasmids. See Tables S2, S3, S4, and S5 for circuits.**

### 3. Cloning details

#### A. Bacterial strains and growth conditions

A complete list of the synthetic circuits can be found in Tables S2, S3, S4, and S5. A complete list of strains can be found in Table. S6.

The MG1655 *E. coli* strain used (SKA360 from [5]) for the microfluidic experiments with circuits 1, 2, 3, 6, 7, 8, 9, 10, and 11 had gene attB::*lacYA177C* and the following deletions: Δ*araCBAD*, Δ*lacIZYA*, Δ*araE*, Δ*araFGH*, Δ*rhaSRT*, and Δ*rhaBADM*. The strain used for circuit 5 was identical with an added *attn7::PRNA1-mCherry-mKate2-hybrid* chromosomal gene. This last strain was also used in Fig. S10 for the strain with no plasmid.

The MG1655 *E. coli* strain used (NDL162 from [6]) for the microfluidic experiments with circuits 12, 14, and 15 had chromosomal gene *glmS::PRNA1-mCherry-mKate2-hybrid* and a motility deletion Δ*motA*. The strain used for circuit 13 was identical with the additional Δ*rpoS::FRT-Kan-frt*. The strain used for circuit 4 was identical to NDL162 but without the chromosomal gene *glmS::PRNA1-mCherry-mKate2-hybrid*.

The standard heat shock procedure was used to transform plasmid DNA into chemically competent *E. coli*. During cloning procedures, *E. coli* strains were grown in LB medium (1% tryptone, 0.5% yeast extract, 1%NaCl) with aeration at 250 rpm or LB agar plates at 37$^o$C with appropriate antibiotics at standard concentrations: Ampicillin 100 $\mu$g/ml (Amp) and Kanamycin 50 $\mu$g/ml (Kan) (antibiotics and LB medium sourced from Fisher Scientific).

Prior to a mother machine experiment, cells were grown overnight in LB medium with appropriate antibiotics from glycerol stocks. At ~3-4 hours prior to the experiment start time, the overnight cultures were diluted 1:100 in imaging media consisting of M9 salts, 10% (v/v) LB, 0.2% (w/v) glucose, 2 mM MgSO4, 0.1 mM CaCl2, 1.5 $\mu$M thiamine hydrochloride and 0.85 g/L Pluronic F-108 (Sigma Aldrich, surfactant to prevent cells sticking to the surface of a microfluidic device). For experiments with circuits 2, 7, 8, and 9 in which IPTG is used, the respective concentrations of IPTG shown in Tables. S2, S3, and S4 are added to the imaging media given to the cells at this point and throughout the remainder of the experiment.

#### B. Plasmid construction

A complete list of plasmids can be found in Table. S6. Snapgene files are available on GitHub. Primers and PCR fragments are labeled in the Snapgene files.

Plasmids on the left side of Tables S2, S3, S4, and S5 were constructed using Gibson assembly (NEBuilder® HiFi DNA Assembly Master Mix, NEB, MA, USA). Reactions were performed according to the manufacturers protocols, with Gibson assembly reactions incubated for 15 min at 50°C.

Primers were ordered from Thermo Fisher Scientific and PCR were performed with Accuprime Pfx (Thermo Fisher) or Phusion polymerase (New England BioLabs) according to manufacturers protocols. All plasmids were verified by DNA sequencing, either Sanger sequencing by Quintara Biosciences (MA, USA) or whole plasmid nanopore sequencing with Plasmidsaurus certified by Oxford Nanopore Technologies.

Various fragments from plasmids built in [2] were amplified by PCR with the ordered primers. Most notably is pLPT119, which formed the backbone for all the plasmid constructs. It consists of the repressilator with no degradation tags on the *lacI*, *cI*, and *tetR* genes [2].

To assemble pEJS1 and pEJS10 by phusion reaction, two fragments were amplified from pLPT119 and make up the three repressors from the repressilator, SCFP3A with PLlacO-1 promoter was amplified from pLPT107, and mVenus NB was amplified from pTP85.

To assemble pEJS3 and pEJS4, SCFP3A with PLlacO-1 promoter was amplified from pLPT107, mCherry-mKate2-hybrid with PLtetO-1 promoter was amplified from pLPT27, mVenus fusion from pEJ1, and repressilator fragments were amplified from pLPT119.

To assemble pEJS5, SCFP3A with PLlacO-1 promoter was amplified from pLPT107, mCherry-mKate2-hybrid with PRNA1 promoter was amplified from pLPT26, mVenus fusion from pEJS1, and repressilator fragments were amplified from pLPT119.

For the experiment shown in Fig. 4D, we used a pSC101 plasmid with Kanamycin resistance and a *gfpmut2* reporter gene with *gadX* promoter taken from [4] (ordered from the E. coli Promoter Collection from Horizon Discovery).

The *tetR-yfp* fusion was made by removing the stop codon from the *tetR* sequence and replacing it with a short linker sequence followed by the mVenus NB sequence. The first nucleotide of from the mVenus NB start codon is not added. The linker sequence is GGGTCTGACATCCTCGAGT.

All repressilator gene sequences and fluorescent protein sequences are followed by transcription terminator T1 from the *E. coli rrnB* gene.

### 4. Class of systems

We consider an arbitrarily complex time-continuous Markov chain of birth-death processes with unspecified reaction rates, corresponding to a biochemical reaction network in a cell. The state of the system at a given moment in time is given by the numbers $\mathbf{z} = \{z_k\}$, consisting of the abundances of all the chemical species in the cell, and all stochastic processes that influence their chemical reaction rates. The dynamics of the system varies stochastically according to unspecified reaction rates:

$$\mathbf{z} \xrightarrow{W_i(\mathbf{z})} \mathbf{z} + \mathbf{d}_i \quad \text{for} \quad i = 1, 2, 3 \ldots, \tag{4.1}$$

where $W_i(\mathbf{z})$ is the probability per unit time that the i-th reaction occurs, and $\mathbf{d}_i$ is the step size of the i-th reaction.

The reaction rate functions $W_i(\mathbf{z})$ can depend on all the system variables or on a subset of them. The Markov chain can be drawn as a network, where nodes correspond to the components $\{z_k\}$, and directional links corresponding to explicit parameter dependencies in the rates $W_i(\mathbf{z})$. For instance, if the component $z_n$ ever changes with a reaction rate that depends explicitly on $z_m$, then an arrow is drawn from the $z_m$ node to the $z_n$ node, see Fig. S17A.

Note that because molecular abundances cannot be negative, any biochemical system must have the property that the propensity of a reaction must be equal to zero when the system is in a state in which an occurrence of that reaction would take one of the abundances to negative numbers. Stochastic networks that do not have this property cannot correspond to biochemical reaction networks. As a result, the abundances $\{z_k\}$ are bounded to be non-negative through reaction rate dependencies. For example, if $z_j$ and $z_k$ correspond to the abundance of molecular species $Z_j$ and $Z_k$ respectively which undergo the following conversion reaction $(z_j, z_k) \xrightarrow{W_i(\mathbf{z})} (z_j - 1, z_k + 1)$, then the transition probability function $W_i$ must have an explicit $z_i$ dependence in that the following must be satisfied: $W_i(\mathbf{z}) = 0$ when $z_j = 0$. Even in the particular case where the conversion rate is approximately independent of the educt concentration when the latter is non-zero, the reaction cannot physically occur when $z_j = 0$, and so $W_i$ must have an explicit $z_j$ dependence.

The system probability distribution $P(\mathbf{z}, t)$ for this network evolves according to the chemical master equation [7, 8]:

$$\frac{d}{dt} P(\mathbf{z}, t) = \sum_k \left[ W_k(\mathbf{z} - \mathbf{d}_k) P(\mathbf{z} - \mathbf{d}_k, t) - W_k(\mathbf{z}) P(\mathbf{z}, t) \right]. \tag{4.2}$$

Time evolution equations for the moments $\langle u_k^n \rangle$ of each component follow directly from the master equation .

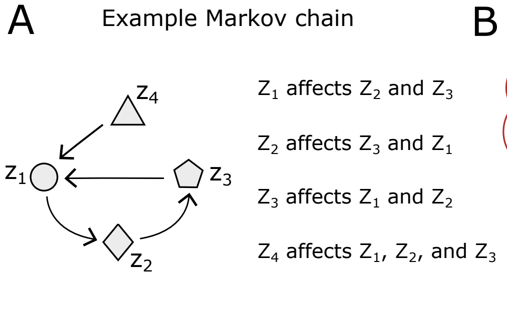


Figure S17. **Definition of the class of systems modelling a gene and its passive reporter. A)** An example Markov chain with the state of the system given by $\mathbf{z} = (z_1, z_2, z_3, z_4)$. Nodes correspond to system variables, whereas arrows correspond to reaction rate dependencies. For instance, if $z_2$ changes under any transition with a reaction rate that depends explicitly on $z_1$, then we draw an arrow from $z_1$ to $z_2$. We say that a variable affects another variable if a path can be drawn in the network topology from the first variable to the second. **B)** We consider all stochastic processes in which two cellular components, $X$ and $Y$, are made with an identical, but unspecified, rate. This rate can depend in any way on a cloud of unknown time-varying components $\mathbf{z}$, which in turn can depend in arbitrary ways on the number of $X$ and $Y$ molecules, denoted by $x$ and $y$. The birth and death reactions of $X$ and $Y$ in Eqs. (4.3) and (4.4) are the only specified parts within this arbitrarily large network. We define two disjoint sets that make up all the unspecified components in the networks. The variables affected by $X$ or $Y$, denoted by $\mathbf{z}_{\text{aff}}$ are all the components in the network that $X$ or $Y$ affects. All other components in the system are denoted by $\mathbf{z}^c_{\text{aff}}$, which can affect the components affected by $X$ in an arbitrary way.

#### A. mRNA reporters

We let $X$ correspond to the transcript of a gene of interest, with $x$ corresponding to the number of $X$ molecules. Functional interactions in this network are specified by the dependencies of the reaction rates in the network. We say that $X$ affects another component $Z$ if a path can be drawn from $X$ to $Z$ in the topology of the network, see Fig. S17A. The set of all components in the network that are affected by $X$ or $Y$ is denoted by $\mathbf{z}_{\text{aff}}$. All other components in the cell are denoted as $\mathbf{z}^c_{\text{aff}}$, see Fig. S17B. We let the $X$ number dynamics to be described by a stochastic birth-death process, with production and degradation reactions of $X$ to be given by

$$x \xrightarrow{R(\mathbf{z}_{\text{aff}},\, \mathbf{z}^c_{\text{aff}})} x+1 \qquad\qquad x \xrightarrow{x\beta(\mathbf{z}^c_{\text{aff}})} x-1, \tag{4.3}$$

where the production rate $R$ can depend in any way on the sets of unspecified components $\mathbf{z}_{\text{aff}}$ and $\mathbf{z}^c_{\text{aff}}$, and the degradation rate $\beta$ can depend in any way on all components not affected by $X$ or $Y$, as long as any possible realization of $\beta_t$ does not decay to zero over time (i.e., $\langle\beta\rangle_t = \lim_{T\to\infty}\frac{1}{T}\int_0^T \beta_t dt > 0$). If $X$ is the transcript of *geneX*, then the rate $R$ corresponds production rate corresponds to the transcription rate of *geneX*.

We next introduce into the system a passive reporter $Y$, engineered to readout the same signal as $X$ that degrades with the same rate

$$y \xrightarrow{\alpha R(\mathbf{z}_{\text{aff}},\, \mathbf{z}^c_{\text{aff}})} y+1 \qquad\qquad y \xrightarrow{y\beta(\mathbf{z}^c_{\text{aff}})} y-1, \tag{4.4}$$

where $\alpha$ is an arbitrary proportionality constant. We extend the definition of $\mathbf{z}_{\text{aff}}$ to include all components affected by $X$ or $Y$. These components can in turn depend in arbitrary ways on the number of $X$ and $Y$ molecules. The dynamics of all other cellular components in the cell thus remains unspecified, and the resulting dynamics need not be Markovian or ergodic in $X$ and $Y$. Note that the sets of unknown components can include an unbounded number of nonphysical mock variables, which can model the different spatial compartments that $X$ can find itself in as well as any complex history dependent dynamics. We only specify that the total $X$ abundance undergoes first-order degradation, where the degradation rate can depend in an arbitrary way on the any of the variables not affected by $X$ or $Y$.

#### B. Fluorescent protein reporters

We can also consider a different class of systems in which $X$ denotes the protein of a gene of interest that is fused to a fluorescent protein. This fluorescent protein must undergo a maturation step before it can fluoresce. We denote the immature fluorescent protein as $X_d$, and the transcript for this protein as $M_x$. The mRNA abundances are described by the following stochastic birth-death processes modelling transcription and mRNA degradation

$$m_x \xrightarrow{R(\mathbf{z}_{\text{aff}},\, \mathbf{z}^c_{\text{aff}})} m_x+1 \qquad m_x \xrightarrow{m_x\beta(\mathbf{z}^c_{\text{aff}})} m_x-1 \tag{4.5}$$

The "dark" fluorescent proteins $X_d$ are translated from the mRNA and then mature into "bright" fluorescent proteins $X$, which are then degraded:

$$x_d \xrightarrow{\lambda(\mathbf{z}^c_{\text{aff}})m_x} x_d+1 \qquad (x_d, x) \xrightarrow{x_d/\tau_{mat}} (x_d-1, x+1) \qquad x \xrightarrow{x\beta_p(\mathbf{z}^c_{\text{aff}})} x-1, \tag{4.6}$$

where $\lambda(\mathbf{z}^c_{\text{aff}})$ is an unspecified translation rate, $\tau_{mat}$ is the average maturation time, and $\beta_p(\mathbf{z}^c_{\text{aff}})$ is an unspecified protein degradation factor. The translation rate $\lambda$ and degradation rate $\beta_p$ can fluctuate and can depend in arbitrary ways on all the components not affected by $X$ or $M_x$ or $Y$ or $M_y$. We assume that any possible realization of $\beta_p$ does not decay to zero over time (i.e., $\langle\beta_p\rangle_t = \lim_{T\to\infty}\frac{1}{T}\int_0^T \beta_{p,t} dt > 0$). The fluorescent proteins are assumed to undergo first-order maturation. In [9] the authors show that approximately a third of a large set of common fluorescent proteins undergo first-order maturation kinetics in bacteria.

We next introduce into the system a passive reporter $Y$ corresponding to a second fluorescent protein reading out the same transcription rate:

$$\begin{gathered} m_y \xrightarrow{\alpha R(\mathbf{z}_{\text{aff}},\, \mathbf{z}^c_{\text{aff}})} m_y+1 \qquad m_y \xrightarrow{m_y\beta(\mathbf{z}^c_{\text{aff}})} m_y-1 \\ y_d \xrightarrow{\alpha_p\lambda(\mathbf{z}^c_{\text{aff}})m_y} y_d+1 \qquad (y_d, y) \xrightarrow{y_d/\tau_{mat}} (y_d-1, y+1) \qquad y \xrightarrow{y\beta_p(\mathbf{z}^c_{\text{aff}})} y-1, \end{gathered} \tag{4.7}$$

where the transcription and translation rates are proportional to those of *geneX* with unknown proportionality constants $\alpha$ and $\alpha_p$. The average maturation and degradation times are equal to those of $X$.

#### 5. The invariant relation — Derivation of Eq. (2)

**Definition 1.** *A co-transition event is a transition event in the system where two or more components change simultaneously. That is, a reaction where the step size* $\mathbf{d}$ *has more than one vector component.*

An example would be a conversion event, where a chemical species $z_m$ converts to another chemical species $z_n$

$$(z_m, z_n) \xrightarrow{W(z_m)} (z_m - 1, z_n + 1).$$

According to our framework, an arrow would be drawn from $z_m$ to $z_n$ in the network if the above reaction was part of the system. Another example would be two molecules $z_m$, $z_n$ that bind to form a complex $z_l$

$$(z_m, z_n, z_l) \xrightarrow{W(z_m,z_n)} (z_m - 1, z_n - 1, z_l + 1).$$

If the above reaction was part of the system, an arrow would be drawn from $z_m$ to $z_n$, from $z_n$ to $z_m$, and from both $z_m$, $z_n$ to $z_l$.

In order to derive Eq. (2), we need to assume that no components in $\mathbf{z}^c_{\text{aff}}$ are part of a co-transition event with any variable in $\mathbf{z}_{\text{aff}}$. We thus begin by proving the following lemma.

**Lemma 1.** *Let $Z_k$ be a component that is not affect by X or Y. For any network of the class in Fig. S17B (Eqs. (4.3) and (4.4)), there exists another network with the exact same dynamics in X, Y, and $Z_k$, but where no components in $\mathbf{z}^c_{aff}$ are part of a co-transition event with any component in $\mathbf{z}_{aff}$.*

*Proof of Lemma 1.* Let the following co-transition reaction be part of the system, where $\{a_k\}$ are variables in $\mathbf{z}_{\text{aff}}$ and $\{b_k\}$ are variables in $\mathbf{z}^c_{\text{aff}}$

$$(\mathbf{a},\ \mathbf{b}) \xrightarrow{W(\mathbf{b})} (\mathbf{a} + \mathbf{d}_a,\ \mathbf{b} + \mathbf{d}_b)\,. \tag{5.1}$$

By definition of $\mathbf{z}_{\text{aff}}$ and $\mathbf{z}^c_{\text{aff}}$, the reaction rate $W$ in the above reaction cannot depend on the variables $\{a_k\}$. However, because $\mathbf{a}$ is part of $\mathbf{z}_{\text{aff}}$, the $\{a_k\}$ variables must be affected by $X$ or $Y$. Therefore, there must exist one or more reactions, labeled with $i \in \{1, 2, \dots\}$, in the system, that change $\mathbf{a}$ with reaction rates that depend on the variables affected by $X$ or $Y$

$$\mathbf{a} \xrightarrow{W_i(\mathbf{z}_{\text{aff}},\ \mathbf{z}^c_{\text{aff}})} \mathbf{a} + \mathbf{d}_i \quad \text{for} \quad i = 1, 2, \dots \tag{5.2}$$

Note that the above reactions cannot make changes to $\mathbf{b}$ otherwise the $\{b_k\}$ variables would not be in the variables not affected by $X$ or $Y$. We now consider an exact copy of the whole system $\mathbf{z}$ and the system reactions, with the change that the $\mathbf{a}$ variables are decomposed into two sets of mock variables. Specifically, we define the variables $\{a^{int}_k\}$ and $\{a^b_k\}$ that undergo the following reactions

$$\begin{gathered}(\mathbf{a}^b,\ \mathbf{b}) \xrightarrow{\quad W(\mathbf{b}) \quad} \left(\mathbf{a}^b + \mathbf{d}_a,\ \mathbf{b} + \mathbf{d}_b\right) \\ \mathbf{a}^{int} \xrightarrow{W_i(\mathbf{z}_{\text{aff}},\ \mathbf{z}^c_{\text{aff}})} \mathbf{a}^{int} + \mathbf{d}_i \quad \text{for} \quad i = 1, 2, \dots,\end{gathered}$$

which correspond to the reactions in Eqs. (5.1) and (5.2). We replace all the explicit $\mathbf{a}$ dependencies in the reaction rates of the system with $\mathbf{a}^{int} + \mathbf{a}^b$. As a result, all the reactions that depended on $\mathbf{a}$ are unchanged, but now we can put the $\mathbf{a}^b$ variables in $\mathbf{z}^c_{\text{aff}}$, and we can put the $\mathbf{a}^{int}$ variables in $\mathbf{z}_{\text{aff}}$. We are left with a system where the $\mathbf{a}^{int}$ are not part of a co-transition event with variables in $\mathbf{z}^c_{\text{aff}}$, and the dynamics of $X$ and $Y$ remain unchanged. Moreover, none of the reaction rates that govern the dynamics of any component in the original $\mathbf{z}^c_{\text{aff}}$ have been altered, meaning the dynamics of $\text{Z}_k$ remain unchanged. Such a decomposition can be done for any co-transition reaction that involve components from $\mathbf{z}_{\text{aff}}$ and $\mathbf{z}^c_{\text{aff}}$. □

We now let $Z_k$ correspond to another component of interest in the system, as a continuous-time Markov process. It can be any stochastic process that can be measured, like the abundance of a molecular species in the network, the size of the cell, or any parameter that can influence the reaction rates of the system.

**Theorem 1.** *Let $Z_k$ be a component in $\mathbf{z}^c_{aff}$. If the averages $\langle x_t \rangle$, $\langle y_t \rangle$ and the covariances $Cov(x_t, z_{k,t})$, $Cov(x_t, z_{k,t})$ over the ensemble have reached a statiorary state (i.e., they have become constant over time), then $\eta_{xz_k} = \eta_{yz_k}$.*

*Proof of Theorem 1.* We condition on the components not affected by $X$ or $Y$, $\mathbf{z}_{\text{aff}}^c[-\infty, t]$. This corresponds to a hypothetical system where all the variables in $\mathbf{z}_{\text{aff}}^c$ become deterministic time-varying signals $\{z_k(t)\}$. The reactions governing $X$ and $Y$ in the conditional system become

$$\begin{aligned} &x \xrightarrow{R(\mathbf{z}_{\text{aff}},\, t)} x+1 && y \xrightarrow{\alpha R(\mathbf{z}_{\text{aff}},\, t)} y+1 \\ &x \xrightarrow{x\beta(t)} x-1 && y \xrightarrow{y\beta(t)} y-1 \end{aligned} \tag{5.3}$$

where $R(\mathbf{z}_{\text{aff}}, t) = R(\mathbf{z}_{\text{aff}},\ \mathbf{z}_{\text{aff}}^c(t))$ and $\beta(t) = \beta(\mathbf{z}_{\text{aff}}^c(t))$ now have an explicit time dependence from the conditioned history $\mathbf{z}_{\text{aff}}^c[-\infty, t]$. We let $A$ be the set of all integers $k$ such that the $k$-th reaction in Eq. (4.1) leads to a change in at least one of the components in $\mathbf{z}_{\text{aff}}$. In the conditional probability space, the components affected in $\mathbf{z}_{\text{aff}}$ follow the following reactions

$$\mathbf{z}_{\text{aff}} \xrightarrow{W_k(\mathbf{z}_{\text{aff}},\, t)} \mathbf{z}_{\text{aff}} + \mathbf{d}_k \quad \forall k \in A, \tag{5.4}$$

where $W_k(\mathbf{z}_{\text{aff}},\ t) = W_k(\mathbf{z}_{\text{aff}},\ \mathbf{z}_{\text{aff}}^c(t))$ now have an explicit time dependence from the conditioned history of the variables not affected by $X$ or $Y$. Note that if some components in $\mathbf{z}_{\text{aff}}^c$ were part of a co-transition event with components in $\mathbf{z}_{\text{aff}}$, then Eqs. (5.3) and (5.4) would not hold. This is because conditioning on the history of those extrinsic variables effectively conditions on those birth events in $\mathbf{z}_{\text{aff}}$ that are caused by those co-transitions. However, from Lemma 1 we can always work with another network in which there are no such co-transition events, and where the dynamics of $X$ and $Y$ remain unchanged.

This conditional system follows the following master equation

$$\begin{aligned} &\frac{d}{dt}P\left(x, y, \mathbf{z}_{\text{aff}}, t | \mathbf{z}_{\text{aff}}^c[-\infty, t]\right) = \\ &\quad \sum_{k\in A}\left[W_k(\mathbf{z}_{\text{aff}} - \mathbf{d}_k, t)P\left(x, y, \mathbf{z}_{\text{aff}} - \mathbf{d}_k, t\middle|\mathbf{z}_{\text{aff}}^c[-\infty, t]\right) - W_k(\mathbf{z}_{\text{aff}}, t)P\left(x, y, \mathbf{z}_{\text{aff}}, t\middle|\mathbf{z}_{\text{aff}}^c[-\infty, t]\right)\right] \\ &\quad + R\left(x-1, y, \mathbf{z}_{\text{aff}}, t\right)P\left(x-1, y, \mathbf{z}_{\text{aff}}, t\middle|\mathbf{z}_{\text{aff}}^c[-\infty, t]\right) - R\left(x, y, \mathbf{z}_{\text{aff}}, t\right)P\left(x, y, \mathbf{z}_{\text{aff}}, t\middle|\mathbf{z}_{\text{aff}}^c[-\infty, t]\right) \\ &\quad + \alpha R\left(x, y-1, \mathbf{z}_{\text{aff}}, t\right)P\left(x, y-1, \mathbf{z}_{\text{aff}}, t\middle|\mathbf{z}_{\text{aff}}^c[-\infty, t]\right) - \alpha R\left(x, y, \mathbf{z}_{\text{aff}}, t\right)P\left(x, y, \mathbf{z}_{\text{aff}}; t\middle|\mathbf{z}_{\text{aff}}^c[-\infty, t]\right) \\ &\quad + (x+1)\beta(t)P\left(x+1, y, \mathbf{z}_{\text{aff}}, t\middle|\mathbf{z}_{\text{aff}}^c[-\infty, t]\right) - x\beta(t)P\left(x, y, \mathbf{z}_{\text{aff}}, t\middle|\mathbf{z}_{\text{aff}}^c[-\infty, t]\right) \\ &\quad + (y+1)\beta(t)P\left(x, y+1, \mathbf{z}_{\text{aff}}, t\middle|\mathbf{z}_{\text{aff}}^c[-\infty, t]\right) - y\beta(t)P\left(x, y, \mathbf{z}_{\text{aff}}, t\middle|\mathbf{z}_{\text{aff}}^c[-\infty, t]\right). \end{aligned}$$

We consider the averages of $X$ and $Y$ conditioned on the upstream history, $\bar{x}(t) = E\left[x_t|\mathbf{z}_{\text{aff}}^c[-\infty, t]\right]$ and $\bar{y}(t) = E\left[y_t|\mathbf{z}_{\text{aff}}^c[-\infty, t]\right]$, where $x_t$ and $y_t$ are the $X$ and $Y$ abundances at time $t$. From the above master equation, the time-evolution for these first moments can be derived [10, 11]

$$\frac{d\bar{x}}{dt} = \bar{R}(t) - \bar{x}\bar{\beta}(t) \quad \& \quad \frac{d\bar{y}}{dt} = \alpha\bar{R}(t) - \bar{y}\bar{\beta}(t), \tag{5.5}$$

where $\bar{R}(t) = E\left[R(x_t, y_t, \mathbf{z}_{\text{aff}}, t)|\mathbf{z}_{\text{aff}}^c[-\infty, t]\right]$ and $\bar{\beta}(t) = E\left[\beta(\mathbf{z}_{\text{aff}}^c)|\mathbf{z}_{\text{aff}}^c[-\infty, t]\right]$ are the average production and degradation rates conditioned on the history of the variables not affected by $X$ or $Y$. Note that $\bar{\beta}(t) = E\left[\beta(\mathbf{z}_{\text{aff}}^c)|\mathbf{z}_{\text{aff}}^c[-\infty, t]\right] = \beta(\mathbf{z}_{\text{aff}}^c(t)) = \beta(t)$, because the time trajectory of $\mathbf{z}_{\text{aff}}^c$ is set through the conditioning on the upstream history. We can then take the expectation of $\bar{x}$ over all possible histories of $\mathbf{z}_{\text{aff}}^c$ to get

$$E[\bar{x}(t)]_{\text{histories}} = E\Big[E\left[x_t|\mathbf{z}_{\text{aff}}^c[-\infty, t]\right]\Big]_{\text{histories}} = \langle x_t\rangle, \tag{5.6}$$

which follows from the law of total expectation. We now let $Z_k$ correspond to any component in the network that is not affected by $X$ or $Y$. It can be a molecular abundance, concentration, or another stochastic cellular variable like the growth rate of the cell. It follows that

$$E[\bar{x}(t)z_k(t)]_{\text{histories}} = E\Big[E\left[x_t|\mathbf{z}_{\text{aff}}^c[-\infty, t]\right]\cdot E\left[z_{k,t}|\mathbf{z}_{\text{aff}}^c[-\infty, t]\right]\Big]_{\text{histories}} = E\Big[E\left[x_t z_{k,t}|\mathbf{z}_{\text{aff}}^c[-\infty, t]\right]\Big]_{\text{histories}} = \langle x_t z_{k,t}\rangle, \tag{5.7}$$

where the second step comes from the fact that conditioning on the history of $\mathbf{z}_{\text{aff}}^c$ effectively also conditions on the history of $Z_k$ with $z_k(t) = E\left[z_{k,t}|\mathbf{z}_{\text{aff}}^c[-\infty, t]\right]$ (so $x$ and $z_k$ are independent when conditioning on the $\mathbf{z}_{\text{aff}}^c$ history), the last step follows from the law of total expectation, and $z_{k,t}$ is the measured amount of $Z_k$ at time $t$. From Eqs. (5.6) and (5.7), it follows that

$$\text{Cov}\left(x_t, z_{k,t}\right) = \text{Cov}\left(\bar{x}(t), \bar{z}_k(t)\right), \tag{5.8}$$

where $\bar{z}_k(t) = E\left[z_{k,t}|\mathbf{z}^c_{\text{aff}}[-\infty, t]\right] = z_k(t)$, where the last step comes from the fact that the time trajectory of $z_k$ is set through the conditioning of the upstream histories. Intuitively, Eq. (5.8) says that when $X$ does not affect $Z_k$, the stochastic fluctuations of $X$ average out when taking the covariance between $X$ and $Z_k$. Strikingly, this is independent of any type of feedback that $X$ may impose through interactions in the cloud of components $\mathbf{z}(t)$. As a result, the same should hold for $Y$, and since $\bar{y}(t)$ is governed by the same differential equation as $\bar{x}$, the intrinsic fluctuations that differentiate $X$ and $Y$ will average out when taking the covariances with $Z_k$.

That is, dividing the right equation in Eq. (5.5) with $\alpha$, we write the general solution for $\bar{x}(t)$ and $\bar{y}(t)/\alpha$, and find

$$\bar{x}(t) = \bar{x}(0)e^{-\int_0^t \beta(u)du} + \int_0^t e^{-\left(\int_0^t \beta(u)du - \int_0^{t'} \beta(v)dv\right)} R(t')dt' \tag{5.9}$$

$$\bar{y}(t)/\alpha = \bar{y}(0)e^{-\int_0^t \beta(u)du}/\alpha + \int_0^t e^{-\left(\int_0^t \beta(u)du - \int_0^{t'} \beta(v)dv\right)} R(t')dt'. \tag{5.10}$$

Taking the average over all histories and subtracting the equations we have

$$\begin{aligned} E[\bar{x}(t)]_{\text{histories}} - E[\bar{y}(t)]_{\text{histories}} &= E\left[\left(\bar{x}(0) - \bar{y}(0)/\alpha\right) e^{-\int_0^t \beta(u)du}\right]_{\text{histories}} \\ \Rightarrow \langle x_t\rangle - \langle y_t\rangle/\alpha &= E\left[\left(\bar{x}(0) - \bar{y}(0)/\alpha\right) e^{-\int_0^t \beta(u)du}\right]_{\text{histories}}, \end{aligned} \tag{5.11}$$

where in the second step we used Eq. (5.6) which also holds for $y$ by symmetry. We now invoke the requirement that the averages $\langle x_t\rangle$ and $\langle y_t\rangle$ are stationary, meaning they are constant over time. In that case, $\langle x_t\rangle - \langle y_t\rangle/\alpha$ is constant over time, and so

$$\langle x_t\rangle - \langle y_t\rangle/\alpha = \lim_{t\to\infty} \left(\langle x_t\rangle - \langle y_t\rangle/\alpha\right). \tag{5.12}$$

Substituting Eq. (5.11), we thus have

$$\langle x_t\rangle - \langle y_t\rangle/\alpha = E\left[\left(\bar{x}(0) - \bar{y}(0)/\alpha\right) e^{-\int_0^\infty \beta(u)du}\right]_{\text{histories}}. \tag{5.13}$$

Now, we must have $\int_0^\infty \beta(u)du \to \infty$ when $\langle\beta\rangle_t = \lim_{T\to\infty} \frac{1}{T}\int_0^T \beta(u)du > 0$. Therefor, the right hand side of Eq. (5.13) becomes 0:

$$\langle x_t\rangle - \langle y_t\rangle/\alpha = E\left[\left(\bar{x}(0) - \bar{y}(0)/\alpha\right) e^{-\int_0^\infty \beta(u)du}\right]_{\text{histories}} = 0, \tag{5.14}$$

Similarly, we now multiply Eqs. (5.9) and (5.10) with $z_k(t)$, average over all histories, and subtract to obtain

$$\begin{aligned} E[\bar{x}(t)z_k(t)]_{\text{histories}} - E[\bar{y}(t)z_k(t)]_{\text{histories}} &= E\left[\left(\bar{x}(0) - \bar{y}(0)/\alpha\right) z_k(t)e^{-\int_0^t \beta(u)du}\right]_{\text{histories}} \\ \Rightarrow \langle x_t z_{k,t}\rangle - \langle y_t z_{k,t}\rangle/\alpha &= E\left[\left(\bar{x}(0) - \bar{y}(0)/\alpha\right) z_k(t)e^{-\int_0^t \beta(u)du}\right]_{\text{histories}} = 0, \end{aligned} \tag{5.15}$$

where in the second step we used Eq. (5.7) which also holds for $y$ by symmetry. The last step follows when $\langle x_t z_{k,t}\rangle$ and $\langle y_t z_{k,t}\rangle$ have reached stationarity and are constant over time, along with $\langle\beta\rangle_t = \lim_{T\to\infty} \frac{1}{T}\int_0^T \beta(u)du > 0$.

It then follows from Eqs. (5.14) and (5.15) that

$$\langle x\rangle = \langle y\rangle/\alpha \quad \& \quad \mathrm{Cov}(x, z_k) = \mathrm{Cov}(y, z_k)/\alpha. \tag{5.16}$$

Dividing $\mathrm{Cov}(x, z_k)$ by $\langle x\rangle\langle z\rangle$, and $\mathrm{Cov}(y, z_k)/\alpha$ by $\langle y\rangle\langle z\rangle/\alpha$, we find $\eta_{xz_k} = \eta_{yz_k}$. □

If the reporter $Y$ is engineered to be passive (i.e., it does not affect components in the network), then a violation of Eq. (2) would imply that $X$ affects $Z_k$. Otherwise, such a violation would imply that $X$ or $Y$ affect $Z_k$.

Thus far we assumed that all the components not affected by $X$ or $Y$ are part of a continuous-time Markov chain. This was in order to make a rigorous definition of causal interaction in our framework as a path in the topology of the transition rates. Alternatively, if we relax the requirement that the components not affected by $X$ or $Y$ be Markov chains (i.e., they can be a set of arbitrary stochastic processes), we can operationally define "no causal interaction from $X$ or $Y$" to mean that we can condition on the history of those stochastic processes and write down Eqs. (5.3) and (5.4). We can then operationally define any violation of Eq. (2) as a "causal interaction from $X$ or $Y$" to a stochastic process $Z_k$.

### 6. Derivation of the invariant relation for fluorescent proteins

The derivation follows analogously to the proof of Theorem 1. We condition on the history of the components not affected by $X$ or $Y$, from which the following differential equations follow like those in Eq. (5.5):

$$\begin{aligned} \frac{d\bar{m}_x}{dt} &= \bar{R}(t) - \bar{m}_x\beta(t) \quad \& \quad \frac{d\bar{m}_y}{dt} = \alpha\bar{R}(t) - \bar{m}_y\beta(t) \\ \frac{d\bar{x}_d}{dt} &= \lambda(t)\bar{m}_x - \bar{x}_d/\tau_{mat} \quad \& \quad \frac{d\bar{y}_d}{dt} = \alpha_p\lambda(t)\bar{m}_y - \bar{y}_d/\tau_{mat} \\ \frac{d\bar{x}}{dt} &= \bar{x}_d/\tau_{mat} - \bar{x}\beta_p(t) \quad \& \quad \frac{d\bar{y}}{dt} = \bar{y}_d/\tau_{mat} - \bar{y}\beta_p(t), \end{aligned} \tag{6.1}$$

where $\lambda(t) = \lambda(\mathbf{z}^c_{\text{aff}}(t))$ and $\beta_p(t) = \beta_p(\mathbf{z}^c_{\text{aff}}(t))$ . When the degradation rates $\beta$, $\beta_p$ do not decay to zero for any possible realization, and when the dual reporter ensemble averages and their covariances with $Z_k$ have reached a time-independent state, it follows from Eq. (6.1) analogously to the derivation from Eq. (5.5) to Eq. (5.16) that $\eta_{xz_k} = \eta_{yz_k}$.

### 7. The X and Y production rates can differ by certain types of fluctuations

In Sec. 5 we assumed that the production rates of $X$ and $Y$ were proportional with an arbitrary proportionality constant. This requirement can be relaxed. Here, we show that the $X$ and $Y$ production rates can differ by certain types of fluctuations. We do this by introducing a cloud of components in the network that model fluctuations in the $X$ or $Y$ production rates that cause them to not be exactly the same.

In particular, consider the following class of systems

$$\begin{aligned} &x \xrightarrow{R_x(\mathbf{z}_{\text{aff}},\mathbf{z}^c_{\text{aff}})} x+1 \qquad && y \xrightarrow{R_y(\mathbf{z}_{\text{aff}},\mathbf{z}^c_{\text{aff}})} y+1 \\ &x \xrightarrow{x\beta(\mathbf{z}^c_{\text{aff}})} x-1 \qquad && y \xrightarrow{y\beta(\mathbf{z}^c_{\text{aff}})} y-1 \end{aligned} \tag{7.1}$$

where $R_x$ and $R_y$ are now different production rates for $X$ and $Y$. Given a component $Z_k$ that is not affected by $X$ or $Y$, we define the following two disjoint sets: $\mathbf{A}$ and $\mathbf{B}$, see Fig. S18. The set of cellular components $\mathbf{A}$ corresponds to all components that affect $Z_k$, along with all components that affect the degradation rate of $X$ and $Y$, along with the component of interest $Z_k$. The set $\mathbf{B}$ corresponds to all components that are not affected by $X$ or $Y$ but that also do not affect $Z_k$. Note that both sets together, $\mathbf{A} \cup \mathbf{B}$, form the set of components not affected by $X$ or $Y$ (which we refer to as $\mathbf{z}^c_{\text{aff}}$ in previous sections). The cloud of components $\mathbf{B}$ can model fluctuations in the $X$ or $Y$ production rates.

Let $R_x(x, y, \mathbf{z}_{\text{aff}}, \mathbf{A}, \mathbf{B})$ and $R_y(x, y, \mathbf{z}_{\text{aff}}, \mathbf{A}, \mathbf{B})$ be the production rates of $X$ and $Y$ respectively. We can condition on the history of $\mathbf{A}$, in which case the following differential equations follow similarly to Eq. (5.5):

$$\frac{d\bar{\bar{x}}}{dt} = \bar{\bar{R}}_x(t) - \bar{\bar{x}}\bar{\bar{\beta}}(t) \quad \& \quad \frac{d\bar{\bar{y}}}{dt} = \bar{\bar{R}}_y(t) - \bar{\bar{y}}\bar{\bar{\beta}}(t), \tag{7.2}$$

where $\bar{\bar{x}}(t) = E\left[x_t|\mathbf{A}[-\infty, t]\right]$, $\bar{\bar{y}}(t) = E\left[y_t|\mathbf{A}[-\infty, t]\right]$, and $\bar{\bar{\beta}}(t) = E\left[\beta(\mathbf{A}(t))|\mathbf{A}[-\infty, t]\right]$, and where

$$\begin{aligned} \bar{\bar{R}}_x(t) &= E\left[R_x(x_t, y_t, \mathbf{z}_{\text{aff}}, \mathbf{A}(t), \mathbf{B}(t))|\mathbf{A}[-\infty, t]\right], \\ \bar{\bar{R}}_y(t) &= E\left[R_y(x_t, y_t, \mathbf{z}_{\text{aff}}, \mathbf{A}(t), \mathbf{B}(t))|\mathbf{A}[-\infty, t]\right]. \end{aligned}$$

These are the average production rates conditioned on the history of the components $\mathbf{A}$. Note that Eq. (7.2) is identical to Eq. (5.5) if the average conditional rates are proportional to one another:

$$\bar{\bar{R}}_x(t) \propto \bar{\bar{R}}_y(t). \tag{7.3}$$

As long as Eq. (7.3) holds, the derivation that follows Eq. (5.5) applies here and it follows that $\eta_{xz_k} = \eta_{yz_k}$.

As a result, $R_x$ and $R_y$ can differ by any fluctuations that come out of the components in $\mathbf{B}$ or those in $\mathbf{z}^c_{\text{aff}}$ that average out when taking the average conditioned on the history of the components in $\mathbf{A}$.

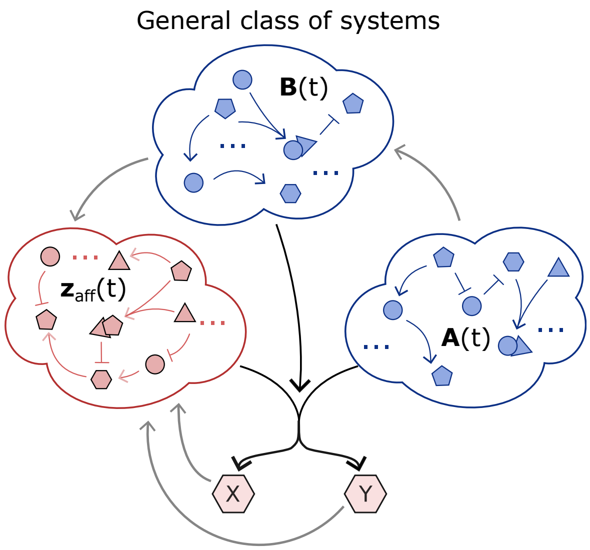


Figure S18. **Definition of the class of systems modelling a gene and its passive reporter.** Given a component $Z_k$ that is not affected by $X$ or $Y$, we define $\mathbf{A}$ to be all the components in the network that affect $Z_k$, along with all the components that affect the degradation rates of $X$ and $Y$, along with the component $Z_k$ itself. We define $\mathbf{B}$ to be all the components that are not affected by $X$ or $Y$ and that are not in the set $\mathbf{A}$. The union of $\mathbf{A}$ and $\mathbf{B}$ make up the cloud of components $\mathbf{z}^c_{\text{aff}}$ in Fig. S17B that are not affected by $X$ or $Y$. We show in this section that the proof of Eq. (2) still holds when the production rates of $X$ and $Y$ differ by fluctuations that arise from the cloud of components $\mathbf{B}$.

**Additive fluctuations:** The covariability relation of Eq. (2) still holds if the production rates differ by some unspecified additive fluctuations that average out to zero. That is, if we have an unspecified additive noise term $\delta$:

$$R_x(x_t, y_t, \mathbf{z}_{\text{aff}}(t), \mathbf{A}(t), \mathbf{B}(t)) = R_y(x_t, y_t, \mathbf{z}_{\text{aff}}(t), \mathbf{A}(t), \mathbf{B}(t)) + \delta(x_t, y_t, \mathbf{z}_{\text{aff}}, \mathbf{A}, \mathbf{B}) \tag{7.4}$$

such that $E\left[\delta(x_t, y_t, \mathbf{z}_{\text{aff}}, \mathbf{A}, \mathbf{B}) | \mathbf{A}[-\infty, t]\right] = 0$. This noise term $\delta$ can depend in an arbitrary way on components in the system, as long as the fluctuations average out to zero when conditioned on the upstream history of $\mathbf{A}$.

For example, we can consider the following birth-death process:

$$\begin{array}{llllll} w \xrightarrow{\quad\lambda\quad} w+1 & p \xrightarrow{\quad\lambda_p\quad} p+1 & x \xrightarrow{w+K(p-\delta\langle p\rangle)} x+1 & y \xrightarrow{\quad w\quad} y+1 & z \xrightarrow{\quad w\quad} z+1 \\ w \xrightarrow{\quad w/\tau\quad} w-1 & p \xrightarrow{\quad p/\tau_p\quad} p-1 & x \xrightarrow{\quad x/\tau\quad} x-1 & y \xrightarrow{\quad y/\tau\quad} y-1 & z \xrightarrow{\quad z/\tau\quad} z-1 \end{array} \tag{7.5}$$

Here $X$ and $Y$ have production rates given by the abundance of $W$, with $X$ having an additional additive term $K(p - \delta\langle p\rangle)$ which depends on the fluctuations in $p$ abundances. Both $W$ and $p$ fluctuate according to a poisson process. Crucially, when $\delta = 1$ the additive noise term will average to zero, in which case the invariant of Eq. (2) must still be satisfied because $X$ and $Y$ do not affect $Z$, see Fig. 7A. However, when $\delta = 0$ the additive noise term does not average to zero, which can lead to a deviation of Eq. (2) even when $X$ and $Y$ do not affect $Z$, see Fig. 7B. This indicates that deviations from Eq. (2) can occur when the $X$ and $Y$ production rates differ by certain types of fluctuations even when $X$ and $Y$ do not affect $Z$.

As an example, an additive noise term of this type can be the result of a linear cascade of an unspecified number

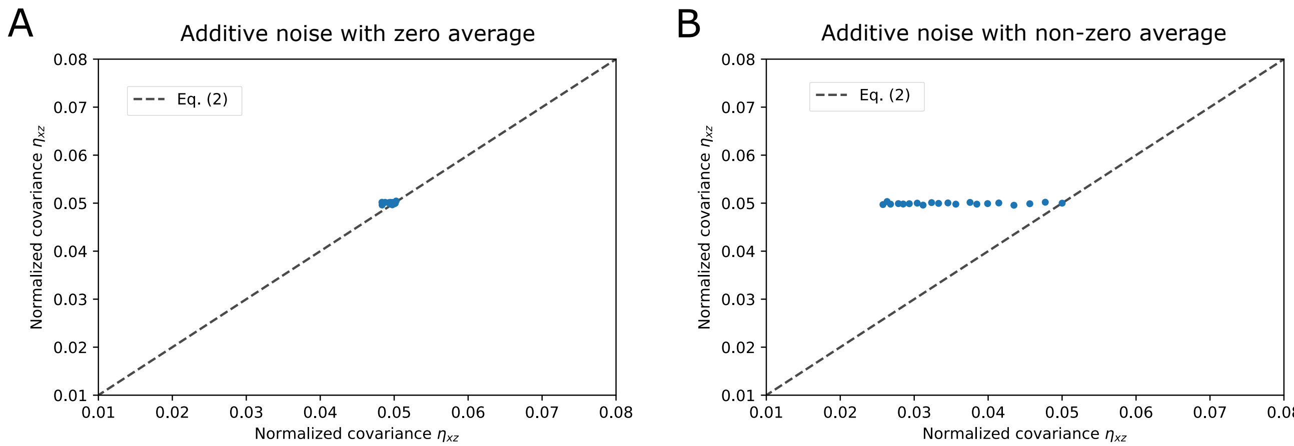


Figure S19. **An example system shows that the invariant holds when the rates differ by additive fluctuations that average to zero. A)** Simulations of the birth-death process of Eq. (7.5) with $\delta = 1$, which ensures that the additive noise term averages to zero. Each dot corresponds to a single iteration of the Gillespie algorithm that was run until each reaction occurred at least $10^7$ times. In each dot the parameters are $\tau = 1$, $\tau_p = 0.1$, $\lambda_p = 5/\tau_p$, $\lambda = 10$. The $K$ parameter quantifies the strength of the additive noise and was varied from 0 to 40 in increments of 2, with each dot having a particular $K$ value. As can be seen, the dots all lie on or very near the line given by Eq. (2). Small deviations from the line are inevitable and are due to numerical errors brought on by finite sampling of the stochastic process. **B)** Same as in **A** except now $\delta = 0$, which means the additive noise term does not average to zero. Here as the $K$ parameter is increased and more noise is added to the $X$ production rate, there is further deviation from Eq. (2) even as $X$ and $Y$ do not affect $Z$.

of variables upstream of $X$ and $Y$. That is, consider the following system

$$
\begin{array}{llll}
x_1 \xrightarrow{R(\mathbf{z}_{\text{aff}},\mathbf{z}^c_{\text{aff}})} x_1 + 1 & x_2 \xrightarrow{\lambda_2 x_1} x_2 + 1 & \ldots & x_n \xrightarrow{\lambda_n x_{n-1}} x_n + 1 \\
x_1 \xrightarrow{\beta_1(\mathbf{z}^c_{\text{aff}})} x_1 - 1 & x_2 \xrightarrow{\beta_2(\mathbf{z}^c_{\text{aff}})} x_2 - 1 & \ldots & x_n \xrightarrow{\beta_n(\mathbf{z}^c_{\text{aff}})} x_n - 1
\end{array}
$$

$$
\begin{array}{llll}
y_1 \xrightarrow{\alpha_1 R(\mathbf{z}_{\text{aff}},\mathbf{z}^c_{\text{aff}})} y_1 + 1 & y_2 \xrightarrow{\alpha_2 \lambda_2 y_1} y_2 + 1 & \ldots & y_n \xrightarrow{\alpha_n \lambda_n y_{n-1}} y_n + 1 \\
y_1 \xrightarrow{\beta_1(\mathbf{z}^c_{\text{aff}})} y_1 - 1 & y_2 \xrightarrow{\beta_2(\mathbf{z}^c_{\text{aff}})} y_2 - 1 & \ldots & y_n \xrightarrow{\beta_n(\mathbf{z}^c_{\text{aff}})} y_n - 1
\end{array}
$$

$$
\begin{array}{ll}
x \xrightarrow{\lambda x_n} x + 1 & y \xrightarrow{\alpha \lambda y_n} y + 1 \\
x \xrightarrow{x\beta(\mathbf{z}^c_{\text{aff}})} x - 1 & y \xrightarrow{y\beta(\mathbf{z}^c_{\text{aff}})} y - 1
\end{array}
$$

where the $\{\lambda_k\}$, $\{\alpha_k\}$, $\lambda$, and $\alpha$ parameters are unspecified constants. The result of this system is that the production rates of $X$ and $Y$ will differ by additive fluctuations, yet Theorem 1 still holds.

**Multiplicative noise that is independent of the production rates:** The covariability relation of Eq. (2) still holds if the production rates differ by some unspecified multiplicative fluctuations that are independent of the production rates:

$$R_x = n_x R(x_t, y_t, \mathbf{z}_{\text{aff}}(t), \mathbf{A}(t), \mathbf{B}(t)) \quad \text{and} \quad R_y = n_y R(x_t, y_t, \mathbf{z}_{\text{aff}}(t), \mathbf{A}(t), \mathbf{B}(t)) \tag{7.6}$$

where $n_x$ and $n_y$ are multiplicative random variables such that

$$
\begin{aligned}
E\left[n_x R(x_t, y_t, \mathbf{z}_{\text{aff}}, \mathbf{A}, \mathbf{B}) | \mathbf{A}[-\infty, t]\right] &= E\left[n_x\right] \cdot E\left[R(x_t, y_t, \mathbf{z}_{\text{aff}}, \mathbf{A}, \mathbf{B}) | \mathbf{A}[-\infty, t]\right] \\
E\left[n_y R(x_t, y_t, \mathbf{z}_{\text{aff}}, \mathbf{A}, \mathbf{B}) | \mathbf{A}[-\infty, t]\right] &= E\left[n_y\right] \cdot E\left[R(x_t, y_t, \mathbf{z}_{\text{aff}}, \mathbf{A}, \mathbf{B}) | \mathbf{A}[-\infty, t]\right]
\end{aligned}
$$

and where $E\left[n_x | \mathbf{A}[-\infty, t]\right] = E\left[n_x | \mathbf{A}[-\infty, t]\right]$. These noise terms $n_x$ and $n_y$ can depend in an arbitrary way on components in the system, as long as they remain independent of the production rate $R$ when conditioned on the upstream history of $\mathbf{A}$.

### 8. Bursty gene expression

In Eukaryotes, a popular model of gene expression assumes that transcription occurs in bursts [12–14]. The dynamics of such bursts are then described by the burst frequency and the burst size. Here we show that the invariant of Eq. (2) holds in the face of transcription bursting, as long as the burst frequency is identical for both *geneX* and the passive reporter *geneY*.

**Coordinated bursting:** Dual reporter genes with identical promoters have been engineered in mammalian cells [13], where it was observed that they underwent bursts in gene expression. Of note, the bursts between these dual reporters were highly correlated when the genes were placed at similar gene loci. On the other hand, the bursts were not correlated when placed at distant gene loci. We first consider the case where the dual reporters undergo perfectly coordinated bursting which would model the former case.

In fact, the class of systems described in Sec. 4 A encompasses such coordinated bursting transcription rates. The rate $R$ is left unspecified, and can vary by switching stochastically between different states, which could model sudden bursts in the production of mRNA. This class of systems can include transcription rates with arbitrary bursting dynamics, frequencies, and burst sizes. The key assumption, however, is that a burst in *geneX* will occur at the same time as a burst in *geneY*.

**Un-coordinated bursting with constant burst sizes:** Bursts in mRNA abundances due to bursty transcription rates can be modelled by adding a burst size $b$ (a positive integer) to the class of systems:

$$\begin{aligned} x &\xrightarrow{R(\mathbf{z}_{\text{aff}},\,\mathbf{z}^c_{\text{aff}})} x + b_x \qquad & x &\xrightarrow{x\beta(\mathbf{z}^c_{\text{aff}})} x - 1, \\ y &\xrightarrow{\alpha R(\mathbf{z}_{\text{aff}},\,\mathbf{z}^c_{\text{aff}})} y + b_y \qquad & y &\xrightarrow{y\beta(\mathbf{z}^c_{\text{aff}})} y - 1. \end{aligned} \tag{8.1}$$

Here, $R$ corresponds to the burst frequency, while $b_x$ and $b_y$ are the sizes of the bursts in the $X$ and $Y$ abundances respectively. Of note, a burst in the $X$ abundances can occur at a different time as a burst in $Y$ abundances (and have a different size). The key assumption here is that the probability that a burst occurs at any given time in the $Y$ abundance is proportional to the respective probability at the same given time for $X$.

The derivation described in Sec. 5 follows for this class of systems, with the exception that a factor of $b_x$ and $b_y$ will go on front of the $\bar{R}(t)$ terms in Eq. (5.5). Nevertheless, the normalized covariances are not affected by these burst sizes, and so the final result remains identical: the invariant of Eq. (2) holds for this class of transcriptional bursting.

It has been hypothesised that chromatin remodeling is a cause of transcriptional bursting in Eukaryotes [14]. Under this model, transcription of a gene is free to occur when the surrounding chromatin is in an open state. When the chromatin is in a condensed state, however, transcription of the gene cannot occur. Under this hypothesis, the class of systems of Eq. (8.1) could correspond to two genes located at different loci that each have the same probability at a given moment in time to suddenly be exposed due to stochastic chromatin remodeling.

**Un-coordinated bursting with distributed burst sizes:** In the previous section and in the class of systems described by Eq. (8.1), the burst sizes $b_x$ and $b_y$ are constant. Here we show that the invariant of Eq. (2) still holds when the burst sizes are distributed. Such distributed burst sizes can be modelled through a series of stochastic reactions that can occur for each possible burst size:

$$x \xrightarrow{R_k(\mathbf{z}_{\text{aff}},\,\mathbf{z}^c_{\text{aff}})} x + k \qquad \text{for } k = 1, 2, 3, \ldots \tag{8.2}$$

Here, a burst of $k$ $X$ molecules can be produced with probabilistic rate $R_k$. The production dynamics of the $Y$ molecules are described by

$$y \xrightarrow{\alpha R_k(\mathbf{z}_{\text{aff}},\,\mathbf{z}^c_{\text{aff}})} y + b_y k \qquad \text{for } k = 1, 2, 3, \ldots \tag{8.3}$$

Here the burst frequencies for each possible burst size can be multiplied by an arbitrary factor $\alpha$, and the burst sizes can also be multiplied by an arbitrary integer $b_y$. Following the same derivation as described in Sec. 5, it follows that the invariant of Eq. (2) holds for this class of systems that models un-coordinated bursting with distributed burst sizes. Note that since the individual rates $R_k$ for each burst size are not specified, any distribution of burst sizes is included.

In this class of systems, a burst is modelled as occurring instantaneously. It remains to be shown that the invariant of Eq. (2) holds in the face of slow bursts in transcription that are un-coordinated between the dual reporters.

#### 9. Simulated example systems from Fig. 1B

To illustrate Eq. (2) in Fig. 1B of the main text, we simulated a number of four-component stochastic birth-death processes with the following general rates

$$x \xrightarrow{r_x^+} x+1 \quad y \xrightarrow{r_y^+} y+1 \quad z_k \xrightarrow{r_z^+} z_k+1 \quad w \xrightarrow{r_w^+} w+1$$
$$x \xrightarrow{r_x^-} x-1 \quad y \xrightarrow{r_y^-} y-1 \quad z_k \xrightarrow{r_z^-} z_k-1 \quad w \xrightarrow{r_w^-} w-1$$

For cases in which $X$ affects $Z$ (circles in Fig. 1B), we simulated the following systems.

1. $r_x^+ = \lambda w, \quad r_y^+ = \alpha\lambda w, \quad r_z^+ = \lambda_z x, \quad r_w^+ = \lambda_w, \quad r_x^- = x/\tau, \quad r_y^- = y/\tau, \quad r_z^- = x/\tau_z, \quad r_w^- = w/\tau_w.$
2. $r_x^+ = \lambda w, \quad r_y^+ = \alpha\lambda w, \quad r_z^+ = \lambda_z \frac{w^2}{0.01+0.1x}, \quad r_w^+ = \lambda_w, \quad r_x^- = x/\tau, \quad r_y^- = y/\tau, \quad r_z^- = x/\tau_z, \quad r_w^- = w/\tau_w.$
3. $r_x^+ = \lambda w, \quad r_y^+ = \alpha\lambda w, \quad r_z^+ = \lambda_z \frac{w^2}{0.01+0.001x}, \quad r_w^+ = \lambda_w, \quad r_x^- = x/\tau, \quad r_y^- = y/\tau, \quad r_z^- = x/\tau_z, \quad r_w^- = w/\tau_w.$
4. $r_x^+ = \lambda w, \quad r_y^+ = \alpha\lambda w, \quad r_z^+ = \lambda_z \frac{x}{0.01+w}, \quad r_w^+ = \lambda_w, \quad r_x^- = x/\tau, \quad r_y^- = y/\tau, \quad r_z^- = x/\tau_z, \quad r_w^- = w/\tau_w.$
5. $r_x^+ = \lambda w, \quad r_y^+ = \alpha\lambda w, \quad r_z^+ = \lambda_z \frac{w^2}{0.01+0.5x+w}, \quad r_w^+ = \lambda_w, \quad r_x^- = x/\tau, \quad r_y^- = y/\tau, \quad r_z^- = x/\tau_z, \quad r_w^- = w/\tau_w.$

For cases in which $X$ does not affect $Z$ (squares in Fig. 1B), we simulated the following systems.

1. $r_x^+ = \lambda w, \quad r_y^+ = \alpha\lambda w, \quad r_z^+ = \lambda_z w, \quad r_w^+ = \lambda_w, \quad r_x^- = x/\tau, \quad r_y^- = y/\tau, \quad r_z^- = x/\tau_z, \quad r_w^- = w/\tau_w.$
2. $r_x^+ = \lambda w, \quad r_y^+ = \alpha\lambda w, \quad r_z^+ = \lambda_z \frac{1}{1+w}, \quad r_w^+ = \lambda_w, \quad r_x^- = x/\tau, \quad r_y^- = y/\tau, \quad r_z^- = x/\tau_z, \quad r_w^- = w/\tau_w.$

For each system, 1000 simulations were performed, each time randomly picking the following parameters $\alpha$, $\tau_z$, $\tau_w$, $\lambda$, $\lambda_z$, and $\lambda_w$ from the respective sets $[0,2]$, $[0,2]$, $[0,2]$, $[0,10]$, $[0,10]$, and $[0,10]$, where $[a,b]$ is defined as the set of all real numbers $n$ that satisfy $a \leq n \leq b$. The parameter $\tau$ was set to 1 throughout. Simulations were performed using the Gillespie algorithm [15] using C. Trajectories were simulated for $10^7$ reaction events. Normalized covariances were computed by integrating over the trajectories to obtain time averages for first and second moments. This is equivalent to using the distribution given by calculating the fraction of the total system time spent in each sampled state. In the ergodic regime, this distribution converges to the stationary distribution of the ensemble.

No violations of Eq. (2) were found, with plotted circles and squares corresponding to a representative sub-sample, with arbitrarily chosen sampling density, of our numerical simulations to indicate the accessible space

#### 10. Example system with feedback from Fig. S1

For Fig. S1 we simulated the following stochastic birth-death process

$$\begin{array}{lll} z \xrightarrow{\lambda_z} z+1 & x \xrightarrow{R(x,z)} x+1 & y \xrightarrow{R(x,z)} y+1 \\ z \xrightarrow{z\beta_z} z-1 & x \xrightarrow{x\beta} x-1 & y \xrightarrow{y\beta} y-1 \end{array} \quad \text{with} \quad R(x,z) = \lambda\left(1+\left(\frac{x}{z+b}\right)^n\right)^{-1} \tag{10.1}$$

where $\lambda_z = 0.02$, $\beta_z = 0.001$, $\lambda = 10$, $b = 0.01$, $\beta = 0.01$, and $n$ was varied to tune the feedback strength. In this system $Z$ affects both $X$ and $Y$ because the rate $R$ depends on $z$. On the other hand, $X$ is not affected by $X$ and $Y$. The rate $R$ also depends on the $X$ numbers, leading to negative feedback in $X$ but not in $Y$. This system was simulated using the Gillespie algorithm [15].

For Fig. S1a we set $n = 20$, and simulated the system for 100,000 reaction events. The initial conditions at $t = 0$ were $z = x = y = 0$. The time-trace shown in Fig. S1a is a small segment of the total simulated time-trace, whereas the shown distributions are made by pooling the entire time-trace into distributions (the system is ergodic). The $Z$ abundance was given a constant offset of 3 in Fig. S1a for illustrative purposes to align the time-traces.

As a measure of feedback strength in this system we use the absolute value of the logarithmic gain taken at the averages

$$\text{Feedback strength} = \left|\left(\frac{\partial \ln(R(x,z))}{\partial \ln(x)}\Big|_{\langle x\rangle,\langle z\rangle}\right)\right| = n\left(\frac{\langle x\rangle^n}{(\langle z\rangle+b)^n + \langle x\rangle^n}\right).$$

This quantifies how the rate $R$ changes with $x$ [16], and for this system is approximately equal to $n$. For Fig. S1b we varied the feedback strength by simulating systems with different $n$ from 0 to 9. Each simulation ran for $10^7$ reaction events, and the Pearson correlation coefficients and the normalized covariances were computed from the distributions pooled from the sample paths as described above.

#### 11. Details on the models simulated in Fig. 2

Here we provide details of the stochastic birth-death processes that were simulated to generate the normalized covariances plotted in Fig. 2. The full model rates with selected parameters are described in Tab. S1. Additional details on the computation of the normalized covariances, as well as sampling error estimations, are discussed in the Materials and Methods of the main text. The algorithm used to simulate these systems is discussed in Sec. 13.

In each of the processes descriptions below, we first show the network topology, followed by the stochastic birth-death reactions. Note that some processes share the same topology, but differ in the rate functions. Moreover, we denote $[x]$ as the concentration of $X$, defined as $x/V(t)$, where $V(t)$ is the cell volume at time $t$ (similarly for $[y]$, $[z]$, and $[w]$) . Some of the rates depend directly on the cell volume $V(t)$. In these reactions we write $V(t)$ as $V$ in the rate functions to shorten the notation, but it should be noted that when $V$ is written in a rate function it still corresponds to the cell volume at time $t$. The cell volume varies over time along with the molecular abundances, and three different volume dynamics are simulated as described in the Materials and Methods section of the main text.

Note that in most of the systems below we included a feedback mechanism. This is because feedback can make the dynamics of $X$ and $Y$ vastly different even though they are under the same control (see Fig. S1. We thus added feedback in most of these systems to demonstrate that the invariant of Eq. (2) holds even when $X$ and $Y$ have different dynamics.

**Process 1 — Negative feedback in X but not Y:**

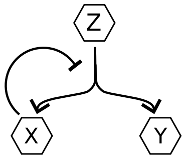


$$x \xrightarrow{\frac{\lambda z}{K+x}} x+1 \quad y \xrightarrow{\frac{\lambda z}{K+x}} y+1 \quad z \xrightarrow{\lambda_z} z+1$$
$$x \xrightarrow{\beta x} x-1 \quad y \xrightarrow{\beta y} y-1 \quad z \xrightarrow{\beta_z z} z-1$$

Here the $X$ and $Y$ production rate depends linearly on the $Z$ abundances, but is also suppressed by $X$ abundances through negative feedback. The $Z$ component is produced according to a poisson process. This could model, for example, a gene regulated by an upstream transcription factor that also has an auto repression mechanism. All components in this model are degraded with a first-order reaction with constant degradation constants.

**Process 2 — Asymmetric feedback:**

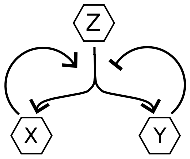


$$x \xrightarrow{\frac{\lambda(x+\delta)}{\left(K+x+\left(\frac{y}{z+\delta}\right)^2\right)}} x+1 \quad y \xrightarrow{\frac{\lambda(x+\delta)}{\left(K+x+\left(\frac{y}{z+\delta}\right)^2\right)}} y+1 \quad z \xrightarrow{\lambda_z} z+1$$
$$x \xrightarrow{\beta x} x-1 \quad y \xrightarrow{\beta y} y-1 \quad z \xrightarrow{\beta_z z} z-1$$

Though we state in the main text that $Y$ be a passive reporter that does not affect other components in the network, in the proof of the invariant of Eq. (2) we only assume that $Y$ does not affect the $Z$ component of interest. Therefor, as long as $Y$ does not affect $Z$, the invariant of Eq. (2) should hold even if $X$ and $Y$ are involved in a feedback mechanism that depends on different ways on the $X$ and $Y$ abundances. To test this, we simulated the process above. Here, the $X$ and $Y$ production rate is a hill function that increases with $X$ and $Z$ abundances, but has noise suppression in $Y$ abundances. The $\delta$ parameter is a small number that is added to stop the rate from ever reaching zero when $x = 0$ (otherwise the rate would stay at zero at the first moment that the $X$ abundances hit zero). There is also a $\delta$ in the denominator to stop the denominator from ever becoming zero which would lead to numerical error.

**Process 3 — Negative feedback with confounding variable:**

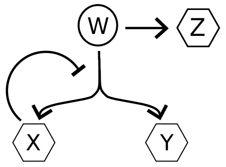


$$x \xrightarrow{\frac{\lambda w}{K+x^2}} x+1 \quad y \xrightarrow{\frac{\lambda w}{K+x^2}} y+1 \quad z \xrightarrow{\lambda_z w} z+1 \quad w \xrightarrow{\lambda_w} w+1$$
$$x \xrightarrow{\beta x} x-1 \quad y \xrightarrow{\beta y} y-1 \quad z \xrightarrow{\beta_z z} z-1 \quad w \xrightarrow{\beta_w w} w-1$$

Next, we test the invariant on a process in which $Z$ is correlated with $X$ and $Y$ through a confounding variable $W$. Here, $X$, $Y$, and $Z$ are produced with rates that are proportional to the $W$ abundances. However, the $X$ and $Y$ production rate is also suppressed by the $X$ abundances through negative feedback. This could model, for example, a transcription factor that regulates $X$, $Y$, and $Z$, in which $X$ is involved in an auto repression mechanism.

**Process 4 — Negative feedback with concentration dependence 1:**

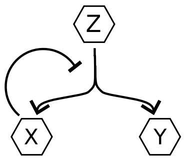


$$x \xrightarrow{\frac{\lambda[z]}{1+[x]^2}} x+1 \quad y \xrightarrow{\alpha\frac{\lambda[z]}{1+[x]^2}} y+1 \quad z \xrightarrow{\lambda_z} z+1$$
$$x \xrightarrow{\beta x} x-1 \quad y \xrightarrow{\beta y} y-1 \quad z \xrightarrow{\beta_z z} z-1$$

In the previous systems, the rates depend on the abundances of the components. However, reaction rates can depend on the concentration of components instead of the abundances, which will exhibit different dynamics in growing and dividing cells. In order to demonstrate that the invariant of Eq. (2) holds when the reaction rates depend on concentrations, we simulated the above process. Here, the production rate of $X$ and $Y$ increases linearly with the $Z$ concentration, but is suppressed by the $X$ concentration. The $Z$ production rate is a poisson process, and all components undergo first order degradation with constant degradation constant.

**Process 5 — Negative feedback with concentration dependence 2:**

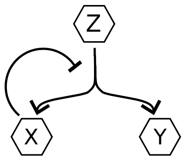


$$x \xrightarrow{\alpha\frac{\lambda V}{1+\left(\frac{[x]}{\delta+[z]}\right)^3}} x+1 \quad y \xrightarrow{\frac{\lambda V}{1+\left(\frac{[x]}{\delta+[z]}\right)^3}} y+1 \quad z \xrightarrow{\lambda_z V} z+1$$
$$x \xrightarrow{\beta x} x-1 \quad y \xrightarrow{\beta y} y-1 \quad z \xrightarrow{\beta_z z} z-1$$

Here we simulated another process with rates that depend on the component concentrations. However, here the rates scale with the cell volume $V$. This is to model a transcription rate that scales with the cell volume. In growing and dividing cells, this scaling in the transcription rate can allow for the mRNA concentration to be on average constant throughout the cell-cycle, which has been observed in many genes. The $\delta$ parameter is added so that the denominator does not ever go to zero which would cause numerical error.

**Process 6 — Shared time-varying upstream signal:**

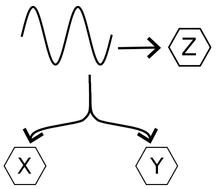


$$x \xrightarrow{A(\sin(\omega t)+1)V} x+1 \quad y \xrightarrow{B(\sin(\omega t)+1)V} y+1 \quad z \xrightarrow{C(\sin(\omega t)+1)V} z+1$$
$$x \xrightarrow{\beta x} x-1 \quad y \xrightarrow{\beta y} y-1 \quad z \xrightarrow{\beta_z z} z-1$$

Here we consider a process where the $X$, $Y$, and $Z$ production rates are driven by a common upstream signal. This signal is a deterministic sine function. The rates scale with the cell volume $V$ so that the rate of production of the concentrations, given by the abundance production rates divided by the cell volume, are themselves sine functions. This could model, for example, three genes that are co-regulated by an oscillating signal.

**Process 7 — Asymmetric feedback with confounding variable and concentration dependencies 1:**

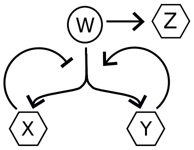


$$x \xrightarrow{\alpha \frac{([y]+\delta)}{[y]+\delta+\left(\frac{[x]}{\delta+[w]}\right)^2}} x+1 \quad y \xrightarrow{\frac{([y]+\delta)}{[y]+\delta+\left(\frac{[x]}{\delta+[w]}\right)^2}} y+1 \quad z \xrightarrow{\lambda_z [w]} z+1 \quad w \xrightarrow{\lambda_w V} w+1$$
$$x \xrightarrow{\beta x} x-1 \quad y \xrightarrow{\beta y} y-1 \quad z \xrightarrow{\beta_z z^2} z-1 \quad w \xrightarrow{\beta_w w} w-1$$

This process is similar to process 2 in that the production rate of $X$ and $Y$ depends on both components in different ways. In particular, the shared production rate of $X$ and $Y$ is a hill function that increases with $Y$ concentration but is suppressed by $X$ concentrations. There is also a confounding variable $W$ that regulates the dual reporters by acting as the hill coefficient in the shared production rate of $X$ and $Y$. The $Z$ production is also regulated by the $W$ component, through a linear dependency on the $W$ concentration.

**Process 8: Interaction mediated by a confounding variable**

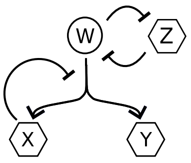


$$x \xrightarrow{\frac{\lambda}{1+\left(\frac{[x]}{\delta+w}\right)^3}} x+1 \quad y \xrightarrow{\frac{\lambda}{1+\left(\frac{[x]}{\delta+w}\right)^3}} y+1 \quad z \xrightarrow{\frac{\lambda_z}{1+\left(\frac{[w]}{k}\right)^3}} z+1 \quad w \xrightarrow{\frac{\lambda_z}{1+\left(\frac{[z]}{k}\right)^3}} w+1$$
$$x \xrightarrow{\beta x} x-1 \quad y \xrightarrow{\beta y} y-1 \quad z \xrightarrow{\beta_z z} z-1 \quad w \xrightarrow{\beta_w w} w-1$$

Here we have a process where the $X$ and $Y$ production rate is a hill function that depends non-linearly on the abundance of component $W$. In addition, the production rate of the $W$ component is repressed by the $Z$ concentration, which itself is repressed by the $W$ concentration. Therefor, $X$ and $Y$ are correlated with $Z$ because they are affected by $Z$ (through $W$), but also because $W$ affects $X$, $Y$, and $Z$.

**System 9 — Fluctuating degradation rates:**

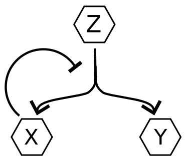


$$x \xrightarrow{\alpha\frac{\lambda z}{K+x}} x+1 \quad y \xrightarrow{\frac{\lambda z}{K+x}} y+1 \quad z \xrightarrow{\lambda_z} z+1$$

$$x \xrightarrow{xz} x-1 \quad y \xrightarrow{yz} y-1 \quad z \xrightarrow{\beta_z z} z-1$$

In the previous process the $X$ and $Y$ components underwent first-order degradation with constant degradation time. The invariant of Eq. (2) holds when the degradation constants fluctuate as long as they are not affected by $X$ or $Y$ (see Sec. 4 A). To demonstrate that the invariant holds with fluctuations in the degradation constants, we simulated the above process where the degradation constant is given by the $Z$ abundance, where the production of $Z$ follows a poisson process. This could model, for example, stochastic fluctuations in the abundances of an enzyme that degrades the $X$ and $Y$ components.

**Process 10 — Oscillating degradation rates**

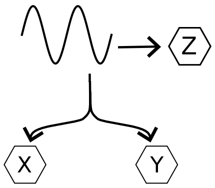


$$x \xrightarrow{A(\sin(\omega t)+1)V} x+1 \quad y \xrightarrow{B(\sin(\omega t)+1)V} y+1 \quad z \xrightarrow{C(\sin(\omega t)+1)V} z+1$$

$$x \xrightarrow{\beta xz} x-1 \quad y \xrightarrow{\beta yz} y-1 \quad z \xrightarrow{\beta_z z} z-1$$

This process is similar to the previous process, but here all the components are driven by a shared upstream oscillating signal. This could model, for example, a system where a degradation enzyme is regulated by an oscillating signal.

### 12. Derivation of Eq. (2) in growing and dividing cells

#### A. Abundances

The class of systems of Eq. (1) does not consider the effect on the $X$ and $Y$ abundances caused by random partitioning of molecules at cell division. Here we show that Eq. (2) still holds when cell division is added to the class of systems. In particular, the $X$ and $Y$ production and degradation reactions are governed by the same Eqs. (4.3) and (4.4), with the addition that the shared unspecified production rate $R$ can now depend on the cell volume $V$ along with the other cellular components: $R(\mathbf{z}_{\text{aff}}, \mathbf{z}^c_{\text{aff}}, V)$.

Additionally, the cell volume divides at unspecified times $\{\tau_k\}$ which can vary over the cell ensemble. At these moments the volume is reduced by multiplicative factors that can lie between 0 and 1. That is, $V \to a_k V$ at $\tau_k$, as we follow one of the daughter cells, where $\{a_k\}$ is a set of factors, with each $a_k$ satisfying $0 < a_k < 1$. As an example, in the case where cells divide perfectly symmetrically at every division, the factors would all be given by $a_k = 1/2$. In general, we allow for stochastic divisions in cell volume, i.e., we let the $a_k$ factors be randomly distributed along with the division times through unspecified ensemble distributions.

At division times, the molecular content of the cell is divided between the two daughter cells. To allow for random partitioning of molecules at cell division, we let the $X$ and $Y$ abundances of a single lineage be reduced by unspecified distributions $\mathcal{A}_k(x)$ and $\mathcal{B}_k(y)$ that can vary over the divisions. We only specify that the averages satisfy $\langle \mathcal{A}_k(x)\rangle = a_k x$ and $\langle \mathcal{B}_k(y)\rangle = a_k y$. That is, on average the abundances are reduced in proportion to the cell division factors. As an example, the $X$ and $Y$ abundances could be divided according to a binomial distribution with probability $a_k$ to remain in the tracked daughter cell.

In total, this amounts to adding the following "reaction" to the class of systems

$$\big(\, V,\ x,\ y\,\big) \xrightarrow{\text{At time } \tau_k} \big(\, a_k V,\ \mathcal{A}_k(x), \mathcal{B}_k(y)\,\big) \quad \text{for } \tau_1,\ \tau_2,\ \tau_3 \ldots$$

To derive Eq. (2), we define the two disjoint sets $\mathbf{z}_{\text{aff}}$ and $\mathbf{z}^c_{\text{aff}}$ similarly to Sec. 4 A. In addition, we include the volume variable $V$ into the set of components not affected by $X$ or $Y$. Similarly to Sec. 5 we consider the average stochastic dual reporter dynamics conditioned on the history of their upstream influences: $\bar{x}(t) = \mathrm{E}\big[x_t|\mathbf{z}^c_{\text{aff}}[-\infty,t], V[-\infty,t]\big]$ and $\bar{y}(t) = \mathrm{E}\big[y_t|\mathbf{z}^c_{\text{aff}}[-\infty,t], V[-\infty,t]\big]$. All cell lineages in this conditional probability space have the same volume history, so they all undergo divisions at the same times $\tau_1, \tau_2, \ldots$, with the same division factors $a_1, a_2, \ldots$ (i.e., these are no longer random variables as they are conditioned on from the conditioning of the volume history). Between any two adjacent division times, $\tau_i$ and $\tau_{i+1}$, the time evolution is specified completely by the reactions in Eqs. (4.3) and (4.4). That is, for $\tau_i < t < \tau_{i+1}$, the time evolution of the conditional averages is given by

$$\frac{d\bar{x}}{dt} = \bar{R}(t) - \bar{x}\beta(t) \quad \& \quad \frac{d\bar{y}}{dt} = \alpha\bar{R}(t) - \bar{y}\beta(t) \quad \text{when} \quad \tau_i < t < \tau_{i+1}, \tag{12.1}$$

where $\bar{R}(t) = E[R(\mathbf{z}_{\text{aff}}, \mathbf{z}^c_{\text{aff}}, V|\mathbf{z}^c_{\text{aff}}[-\infty,t], V[-\infty,t]]$. Division occurs at $t = \tau_{i+1}$, at which point we have $V(\tau_{i+1}) \to a_{i+1}V(\tau_{i+1})$ and

$$\bar{x}(\tau_{i+1}) = \mathrm{E}\big[x_{\tau_{i+1}}|\mathbf{z}^c_{\text{aff}}[-\infty,\tau_{i+1}], V[-\infty,\tau_{i+1}]\big] \to \mathrm{E}\big[\mathcal{A}\left(x_{\tau_{i+1}}|\mathbf{z}^c_{\text{aff}}[-\infty,\tau_{i+1}], V[-\infty,\tau_{i+1}]\right)\big] = a_{i+1}\bar{x}(\tau_{i+1}), \tag{12.2}$$

and similarly $\bar{y}(\tau_{i+1}) \to a_{i+1}\bar{y}(\tau_{i+1})$. This gives us the boundary conditions for the above differential equation. Dividing the right equation in Eq. (12.1) by $\alpha$, we find that the two differential equations are identical in $\bar{x}$ and $\bar{y}/\alpha$, and they have the same boundary conditions. Once the transience of the system has vanished, we are left with $\bar{x}(t) = \bar{y}(t)/\alpha$. It then follows analogously to the proof of Theorem 1 that $\eta_{xz_k} = \eta_{yz_k}$ when $Z_k$ is any component not affected by $X$ or $Y$.

#### B. Concentrations

We now show that Eq. (2) holds in terms of the molecular concentrations of $X$ and $Y$. In particular, we let $x_c := x/V$ and $y := y/V$, where $x$ and $y$ denote the abundances of $X$ and $Y$ respectively. We condition on the history of the upstream influences and the volume:

$$\bar{x}_c(t) = \mathrm{E}\Big[\frac{x_t}{V_t}\Big|\mathbf{z}^c_{\text{aff}}[-\infty,t], V[-\infty,t]\Big] = \frac{1}{V(t)}\mathrm{E}\big[x_t|\mathbf{z}^c_{\text{aff}}[-\infty,t], V[-\infty,t]\big] = \frac{\bar{x}}{V(t)},$$

where $V(t)$ can be pulled out from the expectation brackets since it is specified by the conditioning and becomes equivalent to a constant at time $t$. When $\tau_i < t < \tau_{i+1}$, Eq. (12.1) governs the dynamics of $\bar{x}$ and $\bar{y}$. We use the product rule to find

$$\frac{d\bar{x}_c}{dt} = \frac{d}{dt}\left(\frac{\bar{x}}{V(t)}\right) = \bar{R}_c(t) - \bar{x}_c\beta(t) - \bar{x}_c\frac{V'(t)}{V(t)}, \tag{12.3}$$

where $R_c := R/V$ is interpreted as the production rate of the concentration $X_c$. At division time $\tau_{i+1}$, the cell volume is reduced $V \to a_{i+1}V$ and so is $\bar{x}$ (see Eq. 12.2). We thus have $\bar{x}_c \to \frac{a_{i+1}\bar{x}}{a_{i+1}V} = \bar{x}_c$, i.e., $\bar{x}_c$ is unchanged at the division times. As a result, $\bar{x}_c(t)$ is continuous, meaning Eq. (12.3) holds for all $t$. Note that Eq. (12.3) holds for any volume dynamics between the division times, with the only requirement that it is a differential function between the division times.

We can then take the expectation of $\bar{x}_c$ over all possible histories of $\mathbf{z}^c_{\text{aff}}$ and all possible histories of the volume dynamics $V(t)$

$$\begin{aligned} E[\bar{x}_c(t)]_{\text{histories}} &= E\Big[E\left[x_t/V(t)|\mathbf{z}^c_{\text{aff}}[-\infty,t], V[-\infty,t]\right]\Big]_{\text{histories}} \\ &= E\Big[E\left[x_t/V_t|\mathbf{z}^c_{\text{aff}}[-\infty,t], V[-\infty,t]\right]\Big]_{\text{histories}} = \langle x_t/V\rangle = \langle x_c\rangle, \end{aligned} \tag{12.4}$$

which follows from the law of total expectation. We now consider any component $Z_k$ not affected by $X$ or $Y$. It can be a molecular abundance, concentration, or another stochastic cellular variable like the growth rate of the cell. We then take the following expectation over all possible histories

$$\begin{aligned} E[\bar{x}_c(t) z_k(t)]_{\text{histories}} &= E\Big[E\left[x_{c,t}|\mathbf{z}^c_{\text{aff}}[-\infty,t],V[-\infty,t]\right]\cdot E\left[z_{k,t}|\mathbf{z}^c_{\text{aff}}[-\infty,t],V[-\infty,t]\right]\Big]_{\text{histories}} \\ &= E\Big[E\left[x_{c,t} z_{k,t}|\mathbf{z}^c_{\text{aff}}[-\infty,t],V[-\infty,t]\right]\Big]_{\text{histories}} = \langle x_{c,t} z_{k,t}\rangle, \end{aligned} \tag{12.5}$$

where the second step comes from the fact that conditioning on the history of $\mathbf{z}^c_{\text{aff}}$ effectively conditions on the history of $z_k$ (so $x_c$ and $z_k$ are independent when conditioning on the $\mathbf{z}^c_{\text{aff}}$ history), the last step follows from the law of total expectation, $x_{c,t}$ is the concentration of $X$ at time $t$, $z_{k,t}$ is the measured quantity of $Z_k$ at time $t$, and $z_k(t) = E\left[z_{k,t}|\mathbf{z}^c_{\text{aff}}[-\infty,t],V[-\infty,t]\right]$ which is fully determined through the conditioning of the history of $\mathbf{z}^c_{\text{aff}}$ of which $Z_k$ is an element. From Eqs. (12.4) and (12.5), it follows that

$$\text{Cov}\left(x_{c,t}, z_{k,t}\right) = \text{Cov}\left(\bar{x}_c(t), z_k(t)\right). \tag{12.6}$$

Similarly, we can derive the equivalent differential equation for the Y concentration

$$\frac{d\bar{y}_c}{dt} = \alpha \bar{R}_c(t) - \bar{y}_c\beta(t) - \bar{y}_c\frac{V'(t)}{V(t)}, \tag{12.7}$$

which holds for all $t$. Dividing Eq. (12.7) by $\alpha$, we find that the differential equations governing $\bar{x}_c(t)$ and $\bar{y}_c(t)/\alpha$ are identical. As a result, after the initial transience has decayed, we have $\bar{x}_c(t) = \bar{y}_c(t)/\alpha$. We can thus derive the analogues of Eqs. (12.4) and (12.5) for $\bar{y}/\alpha$. It then follows that

$$\langle x_c\rangle = \langle y_c\rangle/\alpha \quad \& \quad \text{Cov}(x_c, z_k) = \text{Cov}(y_c, z_k)/\alpha. \tag{12.8}$$

Dividing $\text{Cov}(x_c, z_k)$ by $\langle x_c\rangle\langle z_k\rangle$, and $\text{Cov}(y_c, z_k)/\alpha$ by $\langle y_c\rangle\langle z_k\rangle/\alpha$, we find

$$\eta_{x_c z_k} = \eta_{y_c z_k}.$$

If we then re-label the variables and let $x$ and $y$ denote concentrations of $X$ and $Z$ respectively, we obtain Eq. (2).

### 13. Simulation algorithm for growing and dividing cells

Many of the processes in Tab. S1 have time dependent rates through the time-varying cell volume $V(t)$. As a result, we employ the Gillespie algorithm [15] in combination with a "trick" that allows us to simulate exact time trajectories of the abundances and concentrations with time-varying rates [17].

First we simulate the division times and division factors for a single growing and dividing cell. There are three volume dynamics that we consider, see Materials and Methods of the main text. In the first two, division times and division factors are constant. We set the constant division times to 1 and division factors to 1/2. The third is simulated with a simple python script which produces an array of division times $\{\tau_i\}$ and division factors $\{a_i\}$ as described in the Materials and Methods of the main text.

The algorithm is then as follows:

1. The state of the system at time $t_0$ is $\mathbf{s} = (x,\ y,\ z,\ w)$. The waiting time $t$ for the next reaction event to occur is chosen by picking a random number from an exponential distribution with cumulative distribution function $F(t) = 1 - e^{-\int_t^{t_0+t} r_T(\mathbf{s},t)dt}$, where $r_T(\mathbf{s},t)$ is the total rate of reaction events at time $t$:

$$r_T(\mathbf{s},t) = r_x^+(\mathbf{s},t) + r_x^-(\mathbf{s},t) + r_y^+(\mathbf{s},t) + r_y^-(\mathbf{s},t) + r_z^+(\mathbf{s},t) + r_z^-(\mathbf{s},t) + r_w^+(\mathbf{s},t) + r_w^-(\mathbf{s},t),$$

with the rates given in Tab. SS1 and the $t$ dependence comes through the time dependent functions $V(t)$ and $\sin(\omega t)$ in the rates. Determining the waiting time from this distribution requires numerically integrating $r_T(\mathbf{s},t)$ which is computationally time consuming while introducing numerical error. Instead, we use the following trick. We introduce a virtual component $\phi$ that follows the following null rate

$$\phi \xrightarrow{r_\phi(\mathbf{s},t)} \phi, \tag{13.1}$$

where $r_\phi(\mathbf{s},t)$ is a function that satisfies $r_\phi(\mathbf{s},t) + r_T(\mathbf{s},t) = f(\mathbf{s}) > 0$. That is, adding $r_\phi(\mathbf{s},t)$ to $r_T(\mathbf{s},t)$ eliminates the explicit time dependence and leads to a positive function $f$ that only depends on the state $\mathbf{s}$. For example, in process 6 from Tab. S1, we can take $r_\phi(\mathbf{s},t) = \max\{r_T(\mathbf{s},t)\}_t - r_T(\mathbf{s},t)$, where

$$\max\{r_T(\mathbf{s},t)\}_t = 2AV_{max} + 2BV_{max} + 2CV_{max} + x\beta + y\beta + z\beta_z,$$

where $V_{max}$ is the max volume $V(t)$ takes throughout the simulation. For instance, if $V(t)$ exponentially grows from $V_0$ to $2V_0$ and divides back to $V_0$ with constant division times, then $V_{max} = 2V_0$ (the first volume dynamics we consider). In another example, process 3, we could have

$$r_\phi(\mathbf{s}, t) = (1+\alpha)\lambda z/V_{min} + \lambda_z + x\beta + y\beta + z\beta_z - r_T(\mathbf{s}, t).$$

By introducing the null variable $\phi$ into the system the total reaction rate becomes $\tilde{r}_T(\mathbf{s}, t) = r_\phi(\mathbf{s}, t) + r_T(\mathbf{s}, t) = f(\mathbf{s})$, a function that does not explicitly depend on time. As a result, since the abundances $\mathbf{s}$ do not change by definition of the waiting time in $(t_0, t_0+t)$, the waiting time is picked from the following cumulative distribution function

$$F(t) = 1 - e^{-\int_{t_0}^{t_0+t} \tilde{r}_T(\mathbf{s},t)dt} = 1 - e^{-f(\mathbf{s})t}. \tag{13.2}$$

The waiting time $t$ for the next event to occur is thus generated through the cumulative distribution function given by Eq. (13.2), which does not require numerical integration.

2. Check if a cell division $\{\tau_i\}$ occurred between time $t_0$ and $t_0 + t$. If so, the system time is updated to when the division occurs: $t_0 \to \tau_k$ where $\tau_k$ the particular division time that occurred. We then update the cell volume that divides with division factor $a_k$: $V(t_0) \to a_k V(t_0)$. The abundances are then reduced due to cell division and random partitioning

$$(x,\ y,\ z,\ w) \longrightarrow (\ \mathrm{Bin}(x, a_k),\ \mathrm{Bin}(y, a_k),\ \mathrm{Bin}(z, a_k),\ \mathrm{Bin}(w, a_k)\ ),$$

where $\mathrm{Bin}(x, a_k)$ is the binomial distribution on $x$ with probability $a_k$. We then go back to step 1.

3. Else, if no division time occurred between $t_0$ and $t_0 + t$, then update the cell volume $V \to V(t_0 + t)$. Then determine which of the reactions occurs at the time $t_0 + t$, where the i-th reaction occurs with probability $\frac{r_i}{\tilde{r}_T}$. For instance, an $X$ molecule produced with probability $r_x^+(\mathbf{s}, t_0 + t)/\tilde{r}_T(\mathbf{s})$.

4. Update the system according to the reaction that was determined in the previous step. For instance, if the event turns out to be that an $X$ molecule is produced, the system is updated as

$$(x,\ y,\ z,\ w) \longrightarrow (x+1,\ y,\ z,\ w).$$

Note that if the null reaction in (13.1) is picked, nothing occurs.

5. Update the time: $t_0 \to t_0 + t$. Go back to 1 and re-iterate.

### 14. Simulation of a 10 component system with a regulatory cascade

To demonstrate the invariant of Eq. (2) in a system with many components, we simulated the following birth-death process. The following reactions describe the dynamics of the components not affected by $X$:

$$\begin{aligned} &z_1 \xrightarrow{1/(1+z_2^2)} z_1+1, \quad z_2 \xrightarrow{1/(1+z_3^2)} z_2+1, \quad z_3 \xrightarrow{1/(1+z_1^2)} z_3+1, \quad z_4 \xrightarrow{1/(1+z_1^2)} z_4+1, \\ &z_5 \xrightarrow{z_3} z_5+1, \quad z_6 \xrightarrow{z_5} z_6+1, \quad z_7 \xrightarrow{z_1/(1+z_1)} z_7+1, \quad z_7 \xrightarrow{z_7/2} z_7-1. \end{aligned} \tag{14.1}$$

The reactions that govern the $X$ dynamics, as well as the passive reporter $Y$, are given by

$$\begin{aligned} &x \xrightarrow{2z_1/(1+x)} x+1, \quad y \xrightarrow{z_1/(1+x)} y+1, \\ &x \xrightarrow{x} x-1, \quad y \xrightarrow{y} y-1. \end{aligned} \tag{14.2}$$

Here, the production rate of $X$ and $Y$ depends linearly on $Z_1$, but is repressed through a negative feedback loop by $X$. The components that are affected by $X$ form a regulatory cascade with non-linear hill function rates. The reactions that describe the dynamics of the components affected by $X$ are:

$$\begin{aligned} &z_8 \xrightarrow{x^3/\left(1+(x/2)^3\right)} z_8+1, \quad z_9 \xrightarrow{z_8^3/\left(1+(z_8/2)^3\right)} z_9+1, \quad z_{10} \xrightarrow{z_9^3/\left(1+(z_9/2)^3\right)} z_{10}+1, \\ &z_8 \xrightarrow{z_8} z_8-1, \quad z_9 \xrightarrow{z_9} z_9-1, \quad z_{10} \xrightarrow{z_{10}} z_{10}-1. \end{aligned} \tag{14.3}$$

In addition, we simulate growing and dividing cells, where the cell volume $V(t)$ divides by a factor of 2 at each cell division, with a fixed division time of $t_d = 1$. Between two subsequent divisions that occur at times $\tau_i$ and $\tau_{i+1}$ respectively, the volume grows exponentially: $V(t) = 2^{(t-\tau_i)}$. At each division, the abundances of each molecule from the reaction network above are reduced according to a binomial distribution with probability 0.5.

The reaction network topology for this system is shown in Fig. 13A. The normalized covariances computed from simulations of this network are plotted in Fig. 13B. All components not affected by $X$ satisfy the invariant of Eq. (2), whereas the three components that are affected by $X$ display a clear violation of the invariant.

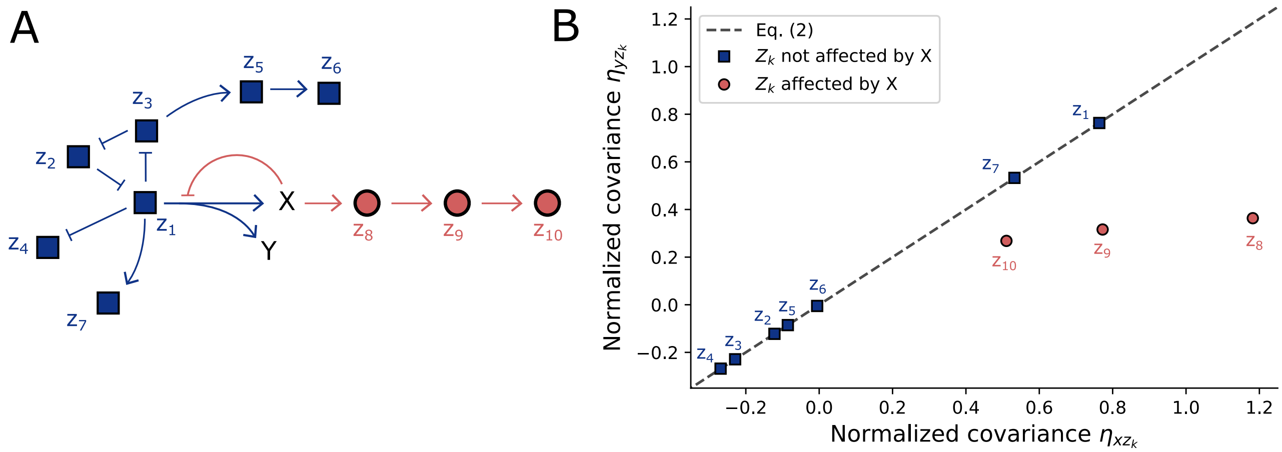


Figure S20. **Eq. (2) is satisfied in an example network made of 10 components.** **A)** The reaction network topology for the system given by Eqs. (14.1), (14.2), and (14.3). Blue squares are components not affected by $X$. Red circles are affected by $X$. **B)** Plotted are the normalized covariances between the *concentrations* of $X$, $Y$ with those of the other components in the network. Components that are not affected by $X$ (blue squares) satisfy the invariant of Eq. (2). Components affected by $X$ (red circles) violate the invariant. Note, the further down the cascade regulated by $X$, the closer the normalized covariances move towards the dashed line. This suggests that the distance from the dashed line is inversely proportional to the relative distances of components down regulatory cascades. Simulations were performed according to the algorithm described in Sec. 13. Normalized covariances were computed by integrating over the trajectories to obtain time averages for first and second moments, waiting for 10,000 cell divisions to occur before integrating to allow the system to reach stationarity. Each computed normalized covariance corresponds to the average of 40 independent simulations that ran for $10^6$ cell divisions. Estimated 95% confidence intervals (which are too minuscule to see) for the normalized covariances correspond to two times the standard error of the mean over these 40 simulations. Using error propagation, in each of the components not affected by $X$, the ratio $\eta_{xz_i}/\eta_{yz_i}$ produced 95% confidence intervals that encompassed the predicted value of 1. See Materials and Methods for additional simulation details.

In Fig. 13B we find that, of the components that are affected by $X$, the normalized covariances with $Z_8$ lie furthest away from the dashed line given by Eq. (2), whereas the normalized covariances with $Z_{10}$ lie closest. This suggests that the "distance" down a regulatory cascade can scale inversely with the degree to which the normalized covariances violate Eq. (2).

Note that the strength of an interaction is not the sole factor that affects the degree to which Eq. (2) is violated. In particular, though the dual reporters $X$ and $Y$ are identically regulated, they will differ due to the probabilistic nature of the reactions governing their dynamics. These "intrinsic fluctuations" are what we exploit to detect causal interactions: when $X$ affects a component of interest $Z$, but $Y$ does not, then the intrinsic fluctuations in $X$ will propagate to $Z$, but those from $Y$ will not, thus creating an asymmetry between the normalized covariances. Therefore, the size of these intrinsic fluctuations affects the difference between the normalized covariances when there is a causal interaction. As a result, the degree to which Eq. (2) is violated cannot equate to an absolute measure for the distance down a regulatory cascade, because both the strength of the interaction and the size of the intrinsic fluctuations in $X$ affect the degree of violation.

However, it might be possible to infer the *relative* distance down a cascade. That is, in a given network, the normalized covariances between $X$, $Y$ and two other components of interest $Z_i$, $Z_j$ that are affected by $X$ can be compared. If the asymmetry between $\eta_{xz_i}$ and $\eta_{yz_i}$ is larger than the asymmetry between $\eta_{xz_j}$ and $\eta_{yz_j}$, then we might be able to conclude that $X$ affects $Z_i$ with a stronger interaction than the interaction from $X$ to $Z_j$, because here the intrinsic fluctuations in $X$ are the same in both cases.

Note that in the particular example in Fig. 13A, the definition of "distance" down the regulatory cascade is clear. However, if there exists feedback from any of the components in the cascade back onto $X$ and $Y$, then this definition is not well defined.

#### 15. Fluctuations in plasmid copy numbers can reduce the degree of violation of Eq. (2)

All 4 of the positive control synthetic circuits used in this study have the target *geneZ* placed on a plasmid. All genes placed on the same plasmid will be impacted by the shared fluctuations that originate from plasmid copy number fluctuations. If the plasmid fluctuations are large enough, they can reduce any differences between $\eta_{xz}$ and $\eta_{yz}$ that result from a causal interaction from $X$ to $Z$. To demonstrate this, we simulated the following two birth-death processes. The following reactions describe the dynamics of an open-loop regulatory cascade:

$$\begin{aligned} &w \xrightarrow{p\lambda} w+1, \quad x \xrightarrow{p\lambda/\left(1+\left(\frac{w^2}{K}\right)\right)} x+1, \quad y \xrightarrow{p\lambda/\left(1+\left(\frac{w}{K}\right)^2\right)} y+1, \quad z \xrightarrow{p\lambda/\left(1+\left(\frac{z}{K}\right)^2\right)} z+1, \\ &w \xrightarrow{w/\tau} w-1, \quad x \xrightarrow{x/\tau} x-1, \quad y \xrightarrow{y/\tau} y-1, \quad z \xrightarrow{x/\tau} z-1. \end{aligned} \tag{15.1}$$

Here, $W$ represses the production $X$ and $Y$, while $X$ represses the production of $Z$. The production rates are proportional to the plasmid copy number $p$ which can also fluctuate. The following reactions describe the dynamics of a closed-loop regulatory cascade:

$$\begin{aligned} &w \xrightarrow{p\lambda/\left(1+\left(\frac{z}{K}\right)^3\right)} w+1, \quad x \xrightarrow{p\lambda/\left(1+\left(\frac{w}{K}\right)^3\right)} x+1, \quad y \xrightarrow{p\lambda/\left(1+\left(\frac{w}{K}\right)^3\right)} y+1, \quad z \xrightarrow{p\lambda/\left(1+\left(\frac{z}{K}\right)^3\right)} z+1, \\ &w \xrightarrow{w/\tau} w-1, \quad x \xrightarrow{x/\tau} x-1, \quad y \xrightarrow{y/\tau} y-1, \quad z \xrightarrow{x/\tau} z-1. \end{aligned} \tag{15.2}$$

Here, $W$ represses the production $X$ and $Y$, $X$ represses the production of $Z$, and $Z$ represses the production of $W$. The production rates are proportional to the plasmid copy number $p$ which can also fluctuate. We simulate the above two systems in two cases. In the first case, there are no plasmid copy number fluctuations, setting $p = 1$ at all times. This corresponds to the case where the genes are all located on the chromosome. In the second case, there are plasmid copy number fluctuations with dynamics modelled by the following birth-death process:

$$p \xrightarrow{\lambda_p} p+1, \quad p \xrightarrow{p/\tau_p} p-1. \tag{15.3}$$

We randomly sample parameters for these systems and find that in the majority of cases, there is a larger degree of violation of Eq. (2) in the case without plasmid fluctuations, see Fig. 13.

#### 16. The invariant relation holds in the face of measurement noise

Here we show that Eq. (2) holds when different types of measurement noise models are added to the class of dual reporter systems given by Eqs. (4.3) and (4.4).

##### A. Multiplicative noise

Let $M_x$ and $M_y$ be identically distributed random variables, independent of $x$ and $y$, that model a multiplicative noise factor for $X$ and $Y$ respectively. If $x_m$ and $y_m$ are the measured abundances of $X$ and $Y$ respectively, then we have

$$x_m = M_x x \quad \& \quad y_m = M_y y, \tag{16.1}$$

where $x$ and $y$ are the true abundances of $X$ and $Y$ respectively. Let $Z_k$ be a component not affected by $X$ or $Y$. If there is measurement noise that affects the $Z_k$ measurement, then we let it be arbitrary and include it in the $z_k$ variable. We only assume that any dependence of $M_x$ and $M_y$ on $z_k$ that occurs as a result of measurement noise is the same, i.e., $E[M_x|z_k] = E[M_y|z_k]$. We use the law of total expectation as follows

$$\langle M_x x z_k \rangle = E\Big[E\big[M_x x z_k | \mathbf{z}^c_{\text{aff}}[-\infty, t], M_x = m\big]\Big] = E\Big[m_x z_k(t) E\big[x|\mathbf{z}^c_{\text{aff}}[-\infty, t]\big]\Big] = E\Big[m z_k(t)\bar{x}(t)\Big],$$

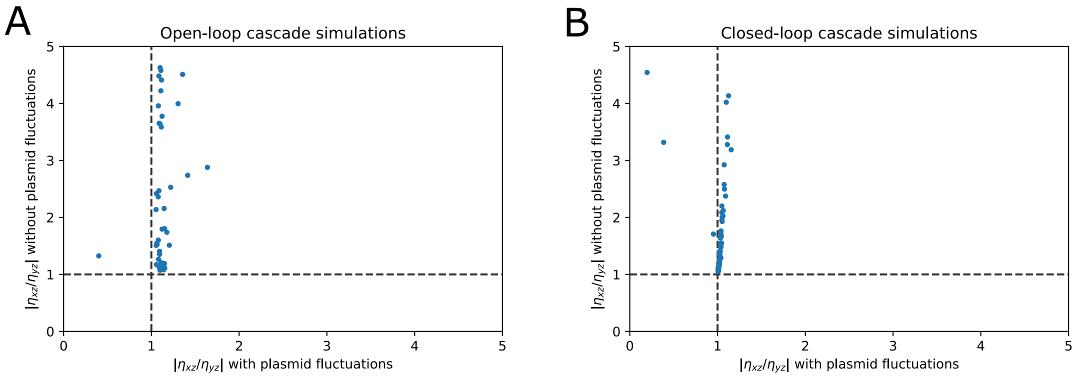


Figure S21. **Fluctuations in plasmid copy numbers reduce the degree of violation of Eq. (2) in simulated test cases. A)** Results of the simulations of the open-loop cascade system given by Eq. (15.1). For each randomly sampled set of parameters, the system was simulated without plasmid fluctuations ($p = 1$ is constant) and with plasmid fluctuations (with dynamics given by Eq. (15.3)). The degree of violation of the invariant of Eq. (2) was computed as the absolute value of the ratio of the normalized covariances $|\eta_{xz}/\eta_{yz}|$. As $X$ in this system represses the $Z$ production rate, there can be a deviation from $|\eta_{xz}/\eta_{yz}| = 1$. We find that the degree of violation of the invariant tends to be bigger in the case without plasmid fluctuations. **B)** Similar to **A** but for simulations of the closed-loop cascade system given by Eq. (15.2). We find that the degree of violation of the invariant tends to be bigger in the case without plasmid fluctuations. For both the open-loop and closed-loop systems, 100 sets of parameters were chosen randomly, with $\tau = 1$, $\lambda = 50r_1$ and $K = 10r_2$ where $r_i$ are uniformly distributed random numbers in $(0, 1)$. Plasmid copy number fluctuations were simulated with $\tau_p = 5r_3$ and $\lambda_p = 5/\tau_p$ which sets the average copy number to 5. Each simulation was performed using the Gillespie alorithm and ran until each reaction occurred at least $10^5$ times.

where $z_k(t)$ is the determined $z_k$ trajectory resulting from conditioning on the history of $\mathbf{z}_k^c$, in the first step we condition on the history of the components not affected by $X$ or $Y$ along with the value of $M_x$, and where the expectation on the right is taken over all the possible histories of $\mathbf{z}_{\text{aff}}^c$ and all the possible values of $M_x$. Similarly, we have $\langle M_y y z_k \rangle = E[m z_k(t)\bar{y}(t)]$. According to the paragraph above Eq. (5.16) we have $\bar{x}(t) = \bar{y}(t)/\alpha$, therefor since $M_x$ and $M_y$ are identically distributed we have

$$\langle M_x x z_k \rangle = \langle M_y y z_k \rangle/\alpha.$$

Similarly we can derive

$$\langle M_x x \rangle = \langle M_y y \rangle/\alpha.$$

It follows from the above two equations and Eq. (16.1) that $\eta_{x_m z_k} = \eta_{y_m z_k}$.

#### B. Additive noise

Let $A_x$ and $A_y$ be random variables, with $\langle A_x \rangle = \langle A_y \rangle = 0$, that model an additive noise term for $X$ and $Y$ respectively. If $x_m$ and $y_m$ are the measured abundances of $X$ and $Y$ respectively, then we have

$$x_m = x + A_x \quad \& \quad y_m = y + A_y, \tag{16.2}$$

where $x$ and $y$ are the true abundances of $X$ and $Y$ respectively. It follows that

$$\langle x_m \rangle = \langle x \rangle \quad \& \quad \langle y_m \rangle = \langle y \rangle.$$

Let $Z_k$ be another cellular component. We assume that the additive noise terms in $X$ and $Y$ are statistically independent of $z_k$. We thus have

$$\mathrm{Cov}(x_m, z_k) = \mathrm{Cov}(x, z_k) \quad \& \quad \mathrm{Cov}(y_m, z_k) = \mathrm{Cov}(y, z_k).$$

It follows that $\eta_{x_m z_k} = \eta_{x z_k}$ and $\eta_{y_m z_k} = \eta_{y z_k}$. If $Z_k$ is a component not affected by $X$ or $Y$, then the invariant relation of Eq. (2) must hold, and so $\eta_{x_m z_k} = \eta_{y_m z_k}$.

#### C. Binomial readout and undercounting noise

Here we show how stochastic undercounting affects the invariant relation of Eq. (2). This is motivated by the fact that common experimental methods can lead to a systematic undercounting of the mRNA-levels and protein-levels. For example, in fluorescence in situ hybridization fluorescent probes only bind to mRNA molecules with a fixed probability. We would thus like to know how the invariant relation of Eq. (2) changes when the reporter abundances $X$ and $Y$ are detected with fixed probabilities $p_x$ and $p_y$ respectively. We let $x_m$ and $y_m$ correspond to the measured $X$ and $Y$ abundances respectively. They are defined as

$$x_m = B(x, p_x) \quad \& \quad y_m = B(y, p_y),$$

where $B(n, p)$ is a binomial distribution with $n$ trials with probability $p$. That is, when each molecule is detect with a fixed probability, the resulting measurement is a binomial readout of the true abundance. Taking the expectation, we have

$$\langle x_m \rangle = E[B(x, p_x)] = E\big[E[B(x, p_x|x)]\big] = E[p_x x] = p_x \langle x \rangle.$$

Now calculating the covariance with cellular component $Z_k$, we have

$$\langle x_m z \rangle = E[B(x, p_x) z_k] = E\Big[E\big[B(x, p_x) z_k | x, z_k\big]\Big] = E\Big[z_k E[B(x, p_x)|x]\Big] = E[p_x x z_k] = p_x \langle xz \rangle$$
$$\Rightarrow \mathrm{Cov}(x_m, z_k) = \langle x_m z_k \rangle - \langle x_m \rangle \langle z_k \rangle = p_x \mathrm{Cov}(x, z_k).$$

As a result, we have

$$\eta_{x_m z} = \eta_{xz} \quad \& \quad \eta_{y_m z} = \eta_{yz},$$

where the second equation follows by symmetry. Therefore, the normalized covariances are unaffected by the molecule detection probabilities. The invariant relation of Eq. (2) thus holds in the face of systematic undercounting of molecules, allowing for arbitrary (and different) detection probabilities for $X$ and $Y$.

#### D. Poisson-Gaussian noise model

Here we consider the case where measurement noise is introduced in fluorescent microscopy images. Fluorscecent imaging noise is often modelled by the Poisson-Gaussian noise model which we consider here [18]. If $\tilde{x}_m$ and $\tilde{y}_m$ are the measured intensities of the $X$ and $Y$ protein fluorescence, then we have

$$\tilde{x}_m = \frac{1}{\chi} \mathcal{P}(\chi \tilde{x}) + \mathcal{N}(0, b),$$

where $\chi \geq 0$ and $b \geq 0$ are arbitrary constants, $\mathcal{P}$ is the Poisson distribution, $\mathcal{N}$ is the normal (Gaussian) distribution, and $\tilde{x}$ is the original fluorescent signal which is proportional to the abundance $x$. Taking the expectation, we have

$$\begin{aligned} \langle \tilde{x}_m \rangle &= E\left[\frac{1}{\chi} \mathcal{P}(\chi \tilde{x})\right] + E\left[\mathcal{N}(0, b)\right] \\ &= \frac{1}{\chi} E\left[E[\mathcal{P}(\chi \tilde{x})|\tilde{x}]\right] + 0 \\ &= \frac{1}{\chi} E\left[\chi \tilde{x}\right] = \langle \tilde{x} \rangle = f_x \langle x \rangle, \end{aligned} \tag{16.3}$$

where $f_x$ is an unknown proportionality constant between the $\tilde{x}$ signal and the protein abundance. That is, the expectation of the measured signal is the same as the expectation of the original signal. We now consider cellular component $Z_k$, which can correspond to the signal of another fluorescent protein in the cell, and can have arbitrary measurement noise. Assuming that the measurement noise in $Z_k$ is independent of the measurement noise in $X$ and $Y$, which could be the case if three different images are taken in three different imaging channels, we take the covariance

$$\begin{aligned} \langle \tilde{x}_m z_k \rangle =& E\left[\frac{1}{\chi} z_k \mathcal{P}(\chi x)\right] + E\left[z \mathcal{N}(0, b) z_k\right] \\ =& \frac{1}{\chi} E\left[E\left[z_k \mathcal{P}(\chi \tilde{x})|\tilde{x}, z_k\right]\right] + E\left[E\left[z_k \mathcal{N}(0, b)|\tilde{x}, z_k\right]\right] \\ =& \frac{1}{\chi} E\left[\chi \tilde{x} z_k\right] + 0 = \langle \tilde{x} z_k \rangle. \end{aligned}$$

Therefor

$$\mathrm{Cov}(\tilde{x}_m, z_k) = \mathrm{Cov}(\tilde{x}, z_k) = f_x \mathrm{Cov}(x, z_k), \tag{16.4}$$

where $f_x$ is an unknown proportionality constant between the $\tilde{x}$ signal and the protein abundance. Putting equations (16.3) and (16.4) together, we have

$$\eta_{\tilde{x}_m z_k} = \eta_{x z_k} \quad \& \quad \eta_{\tilde{y}_m z_k} = \eta_{y z_k},$$

where the second equation follows by symmetry. The normalized covariances are thus not affected by the Poisson-Gaussian noise, and the invariant equation of Eq. (2) can still be exploited.

#### E. Segmentation noise

Let $\tilde{x}_m$ be the signal of the fluorescent $X$ proteins that we measure. The image is made up of pixels which we index with $p$, where the $p$-th pixel has intensity $I_p$. Each cell is segmented, and has cell area $A$ and segmentation area $S$. The measured signal is thus

$$\tilde{x}_m = \sum_{\substack{p\in S\\ p\in A}} I_p + \sum_{\substack{p\in S\\ p\notin A}} I_p.$$

The second term corresponds to additive noise from whenever segmentation fits an area larger than the cell area. This corresponds to fluctuations in the image background. Assuming that the background can be removed and that these fluctuations are negligible compared to the fluorescent protein signal, we can set this term to zero. Moreover, assuming that the fluorescent proteins X are evenly distributed in the cell, then the third term on the right is just the fraction of the total signal that is in the segmentation area

$$\sum_{\substack{p\in A\\ p\in S}} I_p = f_x x \theta,$$

where $\theta = S/A$ when $S < A$ and $\theta = 1$ when $S \geq A$, and $f_x$ is an unknown proportionality constant between the fluorescence and the abundance. We thus have

$$\tilde{x}_m = f_x \theta x,$$

We now let $\theta$ be a stochastic variable that we include in the cloud of variables not affected by $X$, i.e., the segmentation noise is not affected by $x$ through an unspecified Markov chain. In that case, when we condition on the history of upstream variables in $X$, we can still write down Eq. (5.5), and we still have

$$\bar{x}(t) = \bar{y}(t)/\alpha.$$

However, now we are also conditioning on the histories of $\theta$, which means that

$$\begin{aligned}
f_x\theta(t)\bar{x}(t) =& f_y\theta(t)\bar{y}(t)/(\alpha f_y/f_x) \\
&\Rightarrow E\Big[\tilde{x}_m|\mathbf{z}^c_{\text{aff}}[-\infty,t]\Big] = E\Big[\tilde{y}_m|\mathbf{z}^c_{\text{aff}}[-\infty,t]\Big]/(\alpha f_y/f_x) \\
&\Rightarrow E\Big[E\big[\tilde{x}_m|\mathbf{z}^c_{\text{aff}}[-\infty,t]\big]\Big] = E\Big[E\big[\tilde{y}_m|\mathbf{z}^c_{\text{aff}}[-\infty,t]\big]\Big]/(\alpha f_y/f_x) \\
&\Rightarrow \langle\tilde{x}_m\rangle = \langle\tilde{y}_m\rangle/(\alpha f_y/f_x).
\end{aligned}$$

Moreover, we have

$$\langle\tilde{x}_m z_k\rangle = E\Big[E\big[f_x\theta x z_k|\mathbf{z}^c_{\text{aff}}[-\infty,t]\big]\Big] = f_x E\Big[\theta(t)\bar{x}(t)z_k(t)\Big] = \langle\tilde{y}_m z_k\rangle/(\alpha f_y/f_x).$$

We thus have $\eta_{\tilde{x}_m z_k} = \eta_{\tilde{y}_m z_k}$, and Eq. (2) holds in the face of the segmentation noise modelled here.

### 17. Estimating confidence intervals

The corrections from the preceding sections rely on the distribution of measurements. As a result, sampling error affects the corrections and their accuracy, along with the final estimators of the normalized covariances. Here we show two methods we employed to estimate the confidence intervals for the normalized covariances, taking into account sampling error and its effect on the corrections discussed in the preceding sections. The computed normalized covariances and their confidence intervals for both methods are shown in Fig. S16. The two methods give similar results (Figs. 3D,E and S6).

#### A. Sampling with replacement (bootstrapping)

The goal is to estimate the errorbars for a normalized covariance measurement of a strain of cells in a mother machine experiment. Here we use bootstrapping, where the data corrections and the normalized covariances are computed over many samples of the data, allowing for replacement in sampling.

If the experiment produces N fluorescence time-traces of cells with plasmid, and M fluorescence time-traces of cells that lost the plasmid, the following pipeline computes the normalized covariance between the CFP and RFP proteins (the YFP and RFP case follows anagolously), see Fig. S16A.

1. Take a random sample of size N of the cells with plasmid, allowing for replacement. Take another random sample of size M of the cells that lost the plasmid, allowing for replacement.
2. Use the sample of size N to compute the temporal drift curves of the CFP and RFP channels, and drift correct the data from both samples according to the procedure outlined in temporal drift section above.
3. Compute the uneven illumination correction curves of the CFP and YFP channels, using the sample of size N, as outlined in the uneven illumination section above. Correct the data from both samples with the uneven illumination correction curves.
4. From the sample of size M (lost plasmid), pool data into a single distribution and compute the average CFP fluorescence. This corresponds to the autofluorescence and media fluorescence of the CFP channel.
5. Remove the autofluorescence and media fluorescence (obtained in the previous step) from all the CFP measurements from the sample of size N.
6. From the sample of size N, pool data into a single distribution and compute the normalized covariance between CFP and RFP of the sample $\eta^{sample}$ .
7. Go back to step 1 and repeat 100 times, saving each $\eta_{sample}$
8. When 100 normalized covariances $\eta_{sample}$ have been computed, take the average as the final $\eta$ estimate, with confidence intervals given by 2 times the standard deviation of the 100 normalized covariances $\eta_{sample}$.

#### B. Sampling without replacement (splitting data)

The goal is to estimate the errorbars for a normalized covariance measurement of a strain of cells in a mother machine experiment. Here we divide the data from an experiment into $N_{samples}$ disjoint sets. The data corrections and the normalized covariances are computed in each set.

If the experiment produces N fluorescence time-traces of cells with plasmid, and M fluorescence time-traces of cells that lost the plasmid, the following pipeline computes the normalized covariance between the CFP and RFP proteins (the YFP and RFP case follows anagolously), see Fig. S16B.

1. Randomly divide the N cell traces into $N_{samples} = \min(M, 10)$ disjoint samples (each cell is in one and only one sample). Also randomly split the M cell traces with lost plasmid into $N_{samples}$ disjoint sets. This ensures that each disjoint set has at least 1 cell that lost the plasmid to estimate the autofluorescence, but we cap the number of sets to 10 to allow for enough cells in each set. Most strains had more than 10 cells that lost the plasmid in a single mother machine experiment (the lowest was 7).
2. Take a sample of cells with plasmid and a sample of cells that lost plasmid. Compute the temporal drift curves of the CFP and RFP channels using the former sample, and drift correct the data from both samples according to the procedure outlined in temporal drift section above.

3. Compute the uneven illumination correction curves of the CFP and YFP channels, using the sample of cells with plasmid, as outlined in the uneven illumination section above. Correct the data from both samples with the uneven illumination correction curves.

4. From the sample of size M (lost plasmid), pool data into a single distribution and compute the average CFP fluorescence. This corresponds to the autofluorescence and media fluorescence of the CFP channel.

5. Remove the autofluorescence and media fluorescence (obtained in the previous step) from all the CFP measurements from the sample of cells with plasmid.

6. From the sample of cells with plasmid, pool data into a single distribution and compute the normalized covariance between CFP and RFP of the sample $\eta^{sample}$ .

7. Go back to step 2 and repeat until $\eta^{sample}$ has been computed for each of the $N_{samples}$ samples.

8. When the $N_{samples}$ normalized covariances $\eta_{sample}$ have been computed, take the average as the final $\eta$ estimate, with confidence intervals given by 2 times the standard error of mean. Note that the denominator of the standard error is given by $\sqrt{N_{samples} - 1}$ because a degree of freedom is used to compute the average.

---
